# Cell Tracking Profiler: a user-driven analysis framework for evaluating 4D live cell imaging data

**DOI:** 10.1101/859397

**Authors:** Claire Mitchell, Lauryanne Caroff, Alessandra Vigilante, Jose Alonso Solis-Lemus, Constantino Carlos Reyes-Aldasoro, Fabrice de Chaumont, Alexandre Dufour, Stephane Dallongeville, Jean-Christophe Olivo-Marin, Robert Knight

**Affiliations:** Centre for Craniofacial and Regenerative Biology, King’s College London, Tower Wing, Guy’s Hospital, London, U.K; School of Mathematics, Computer Science and Engineering, City, University of London, Tait Building, Northampton Square, London, U.K; Bioimage Analysis Unit, Institut Pasteur, Paris, France; Centre for Stem Cells and Regenerative Medicine, King’s College London, Tower Wing, Guy’s Hospital, London, U.K

**Keywords:** muscle, zebrafish, cell tracking, segmentation, in vivo imaging, Phagosight, Icy, Imaris

## Abstract

Accurate measurements of cell morphology and behaviour are fundamentally important for understanding how disease, molecules and drugs affect cell function in vivo. Using muscle stem cell (muSC) responses to injury in zebrafish as our biological paradigm we have established a ground truth for muSC cell behaviour. This revealed that variability in segmentation and tracking algorithms from commonly used programs are error-prone, leading us to develop a fast semi-automated image analysis pipeline that allows user defined parameters for segmentation and correction of cell tracking. Cell Tracking Profiler (CTP) operates through the freely available Icy platform, and allows user-managed cell tracking from 3D time-lapsed datasets to provide measures of cell shape and movement. Using dimensionality reduction methods, multiple correlation and regression analyses we identify myosin II-dependent parameters of muSC behaviour during regeneration. CTP and the associated statistical tools we have developed thus provide a powerful framework for analysing complex cell behaviour *in vivo* from 4D datasets.

**Summary:** Analysis of cell shape and movement from 3D time-lapsed datasets is currently very challenging. We therefore designed Cell Tracking Profiler for analysing cell behaviour from complex datasets and demonstrate its effectiveness by analysing stem cell behaviour during muscle regeneration in zebrafish.

## Introduction

An ability to measure parameters of cell shape and movement, then to interpret the meaning of such changes, is of fundamental importance for understanding how cells are regulated *in vivo* or in tissue engineering constructs. Advances in *in vivo* and intravital imaging techniques have resulted in the generation of large volumetric datasets in which cells are labelled with multiple fluorescent labels. To characterise cell behaviour from such datasets without extensive distortion of the primary data represents a fundamental problem in image analysis, as cells *in vivo* often show extensive changes in their shape, move rapidly and can be hard to discriminate from adjacent cells if in close proximity (Driscoll and Danuser, 2015).

A number of programs are available for automated analysis of cell behaviour from 3D datasets. These include commercial programs such as Imaris (Bitplane AG) and AMIRA (FEI, Inc.) and open source programs such as ImageJ / Fiji (Collins, 2007; Schindelin et al., 2012), Icy (de Chaumont et al., 2012), Phagosight (Henry et al., 2013), tTy/ qTfy (Hilsenbeck et al., 2016). These use a variety of algorithms to segment and track cells, ranging from user-driven annotation to automated processes with minimal user intervention.

All image analysis programs display similar limitations when attempting to segment 3D datasets in which cells display a uniform labelling including: i) an inability to accurately identify the boundaries of cells, ii) problems in coping with diverse cell sizes and shapes and iii) being able to identify discrete boundaries between adjacent cells (Sbalzarini, 2013). These issues become important when tracking cells, as the segmentation efficiency will dictate how well a cell can be tracked. Further, commercially available software packages do not make their code available, limiting options for users to modify how images are processed.

Several studies have compared software packages for their effectiveness in cell segmentation or tracking (Maska et al., 2014; Ulman et al., 2017). Performance of most software packages has been poor when analysing complex *in vivo* 3D imaging data, as these either rely on specific labels (nuclear or membrane), do not accurately segment cells in 3D, or require computationally expensive methods for segmenting. Given the importance of cell behaviour for effective regeneration, we performed a benchmark study of several available programs (Imaris, Icy and Phagosight) to evaluate their ability to accurately describe migratory muscle stem cells (muSCs) in the myotome of regenerating zebrafish (Knappe et al., 2015; Roy et al., 2017). We found that a fundamental limitation of the programs we tested was the segmentation efficiency and ease of correcting inappropriate tracking. To facilitate analysis of 3D time-lapse data we have therefore designed Cell Tracking Profiler (CTP), an image analysis workflow around the freely available Icy package (de Chaumont et al., 2012) that allows user input to control segmentation and correct tracking of cells. Using a variety of statistical tools we evaluated how outputs from CTP can be analysed to identify parameters of cell shape and movement. Our comparisons against other image analysis packages reveal that CTP offers a powerful and intuitive user-led tool that can provide accurate measures of complex cell behaviour. As an example we used CTP to identify specific changes to muSC shape and movement in response to inhibition of RhoA kinase (ROCK) and myosin II.

A major impediment for interpreting how changes to cell shape and movement are related to specific variables, such as a drug or genetic manipulation, is whether they respond in a coordinated fashion. We evaluated the usefulness of dimensionality reduction methods to address this issue and show that PEER (probabilistic estimation of expression residuals) (Stegle et al., 2012) can be used to identify the response of multiple parameters of cell shape and movement to a variable. Using PEER we reveal that muSC movement shows quantitatively different responses to inhibition of ROCK relative to myosin II. Dissection of individual parameter responses further reveals that specific parameters of muSC responses to tissue injury are altered by inhibition of myosin II activity. Our development of this workflow and proof of principle provides the basis for a medium throughout approach to analysis of time-lapsed datasets from a variety of biological systems. The statistical approaches we have described additionally provide a powerful method for dissecting the effects of several variables in regulating cell behaviour.

## Results

### Morphological changes occur to muSCs during migration to sites of injury

We wished to measure cell shape and behaviour of muSCs, a complex, migratory cell type, in order to determine how they are regulated during tissue repair. Using a zebrafish injury model in which muSCs are labelled with green fluorescent protein (GFP) driven by the *pax7a* promoter (*pax7a:egfp*) (Knappe et al., 2015) we aimed to segment and track cells from time-lapse datasets as accurately as possible. As is common for many transgenic animals, GFP is distributed throughout the cytoplasm of muSCs in *pax7a:egfp* larvae, making it difficult to automatically detect cell boundaries using standard thresholding approaches. To therefore establish a ground truth for cell shape, a manual segmentation of selected cells was performed by hand from three ‘baseline’ datasets (datasets 1-3). This revealed a diversity of muSC sizes and shapes within each individual dataset (Figure 1A). By examining the temporal profile of muSCs in the injured myotome, it was apparent that shape and size for each cell changed over time in a non-uniform manner (Figure 1).

**Figure 1:**
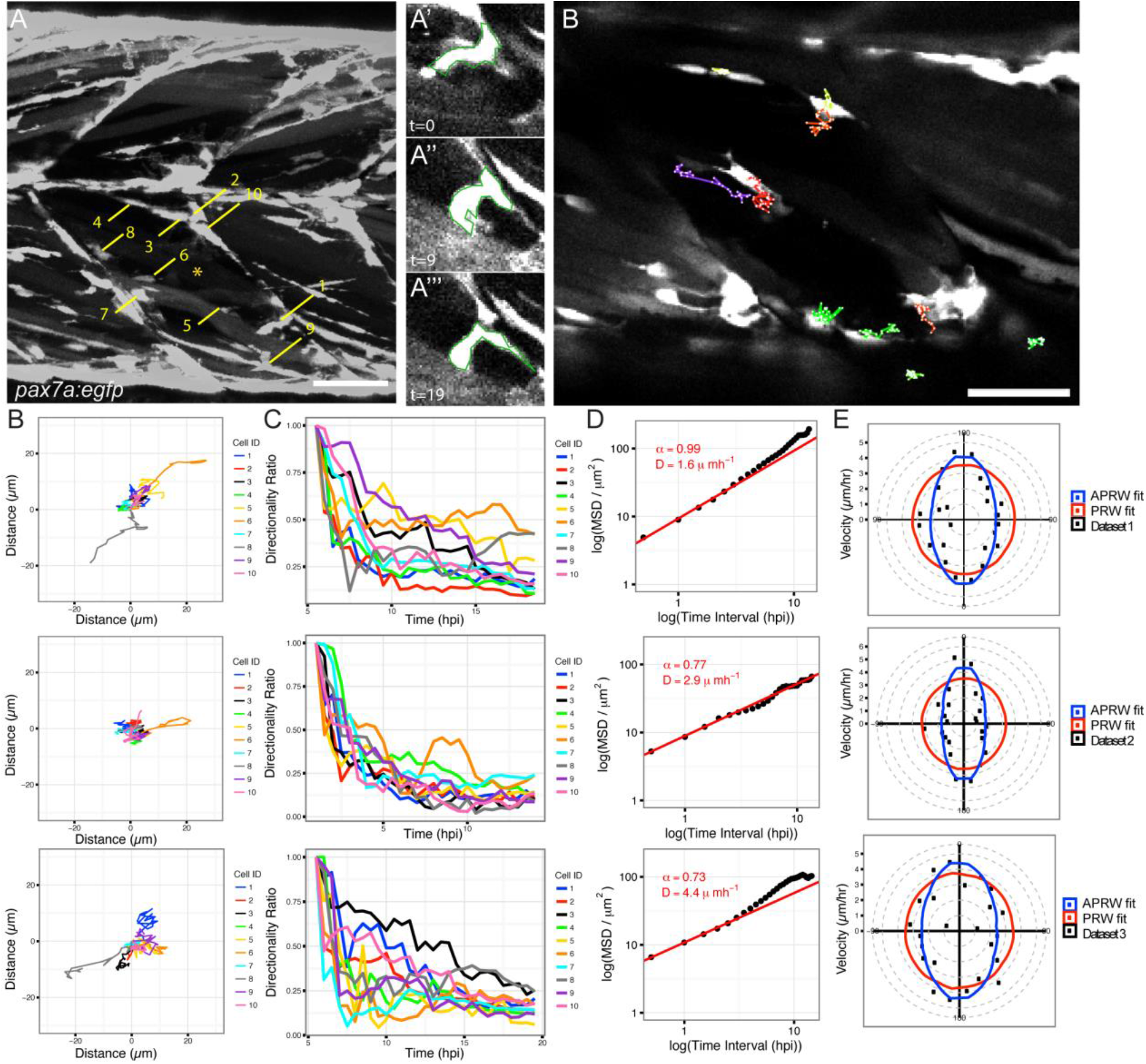
Muscle stem cells in transgenic pax7a:egfp zebrafish larvae show dynamic shape changes as they respond to muscle injury in a time-lapsed movie (dataset 1, acquisition interval 30 minutes). Manual segmentation of 10 cells (numbered, A) in an injured myotome (*) was performed at three time-points (time 0,9,19). Cell 1 changes shape over time (A’ A”, A’”). Selected cells were manually tracked over time using MTrackJ (B). Tracks were mapped to a common origin (C) for 10 selected cells from datasets 1-3 and the directionality ratio plotted (D). Mean Squared Displacement (MSD) was used to calculate D (diffusion constant of cells) and plotted as a logged value against the log of time (E). A line of best fit (red) was generated from which the α parameter was calculated. Cell movement was fitted against models of persistent random walk (PRW) and anisotropic PRW (APRW), with goodness of fit of the data to either model described by an R-squared value. Scale bar 50µm (A,B).

To define how muSCs migrate towards a muscle injury, we established a ground truth by tracking GFP expressing cells in *pax7a:egfp* larvae using the ImageJ manual tracker plugin MTrackJ (Figure 1B-C, Video 1). Each cell was tracked and parameters for duration, point speed and directionality measured (Table 1). To determine if muSCs responding to injury showed rapid movement we evaluated instantaneous speed. This is a calculation of the speed of cell movement in-between time-points and can be used to infer how fast cells move regardless of their directionality. Cells showed considerable variability in their behaviour and no clear trend was discernable. We then tested whether muSCs showed persistence in their movement after injury by plotting the directionality ratio relative to time (Gorelik and Gautreau, 2014). A feature of this measure is that values decay from 1 as a function of their trajectory relative to their initial trajectory, such that cells showing a more persistent movement will show values closer to 1 over time. Variability in cell behaviour of all three datasets was apparent, with some cells showing high persistence lasting for some time (e.g. cells 6 and 9 in dataset 1 and cell 3 in dataset 3), whereas others showed a rapid drop in persistence e.g. cell 1 in dataset 1 (Figure 1D). We calculated the directional autocorrelation of cells to assess whether cell shape is related to the direction of cell migration (Gorelik and Gautreau, 2014), but observed no clear trend over time. An analysis of cell trajectories by Mean Square Displacement (MSD) indicated cell movement was sub-diffusive with alpha parameter values calculated as <1 and diffusion constant (D) > 1 (Figure 1E). MSD assumes cell movement fits a model of persistent random walk (PRW), but recent descriptions of cell behaviour in 3D have suggested that cells move in an anisotropic mode (Wu et al., 2015). We therefore evaluated whether angular velocity of MuSCs fitted better to a model of PRW compared to an anisotropic persistent random walk (APRW, Fig 1F). In all 3 datasets we observed that the velocity of the tracked cells showed a better fit to the APRW model than to a PRW model (Table 1).

**Table 1:**
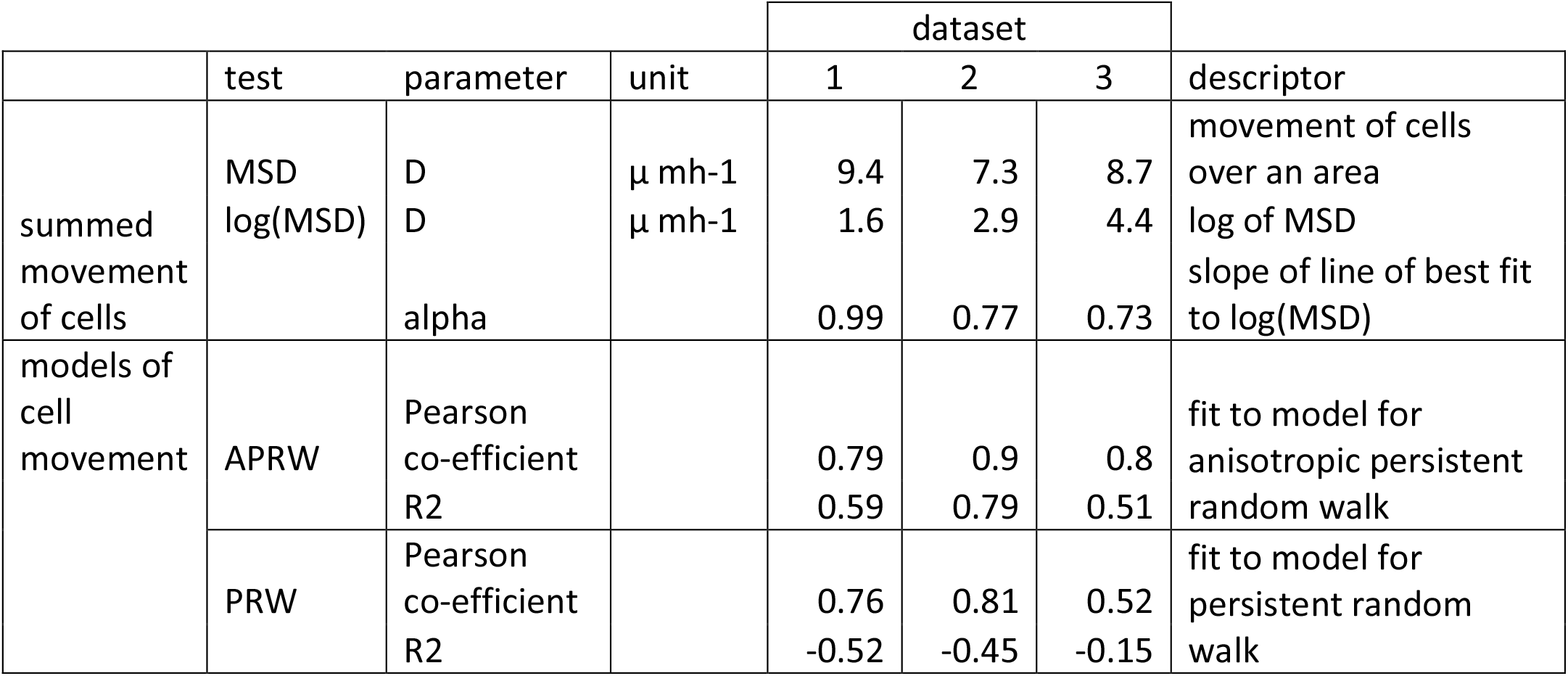
Descriptions of cell movement from datasets 1-3 relative to models of cell movement. Mean squared displacement (MSD) is represented by parameter D (diffusion) and the alpha parameter (a) calculated from the slope of a line of best fit to plots of logged MSD relative to logged time. Fits of cell movement relative to models of persistent random walk (PRW) and anisotropic persistent random walk (APRW) are represented by Pearson co-efficient values and R^2^ values.

### Cell shape affects automated segmentation efficiency

To explore the potential and limits of automated segmentation to identify and measure muSCs we decided to compare three programs that have been used to analyse migratory cell behaviour in a variety of contexts: Imaris (Bitplane AG), Icy (de Chaumont et al., 2012) and Phagosight (Henry et al., 2013). We first evaluated how well each program could segment muSCs cells from datasets 1-3 relative to the ground truth obtained by manual segmentation. The criteria used to compare efficiency of these segmentation processes were 1) how many cells could be identified at three specified time points, 2) how many of a pre-defined population of selected cells of interest at each time point could be identified and 3) similarity of segmented cell volume to volume defined by manual segmentation.

Cell segmentation was performed using Imaris, Icy and Phagosight as described in the Methods section. Optimal parameters for segmentation of cells for datasets 1-3 were determined by comparing to a ground truth, defined by manual quantification or segmentation of cells at three defined times during the time-lapse. Segmentation effectiveness was scored relative to the ground truth at the same time-point following changes to critical parameters. We found that segmentation of cells by Imaris, Icy or Phagosight using default parameters resulted in many more cells identified when compared to the ground truth. Optimisation of segmentation parameters reduced this number, but for datasets 2-3 all three programs still over-segmented more cells than were observed in the ground truth (Figure 2A). Overall, the number of cells identified by Icy was most similar to the ground truth. A comparison of the volumes of the segmented cells showed that Phagosight segmented cells with a wide spread of volumes, in contrast to Icy and Imaris (Figure 2B).

**Figure 2:**
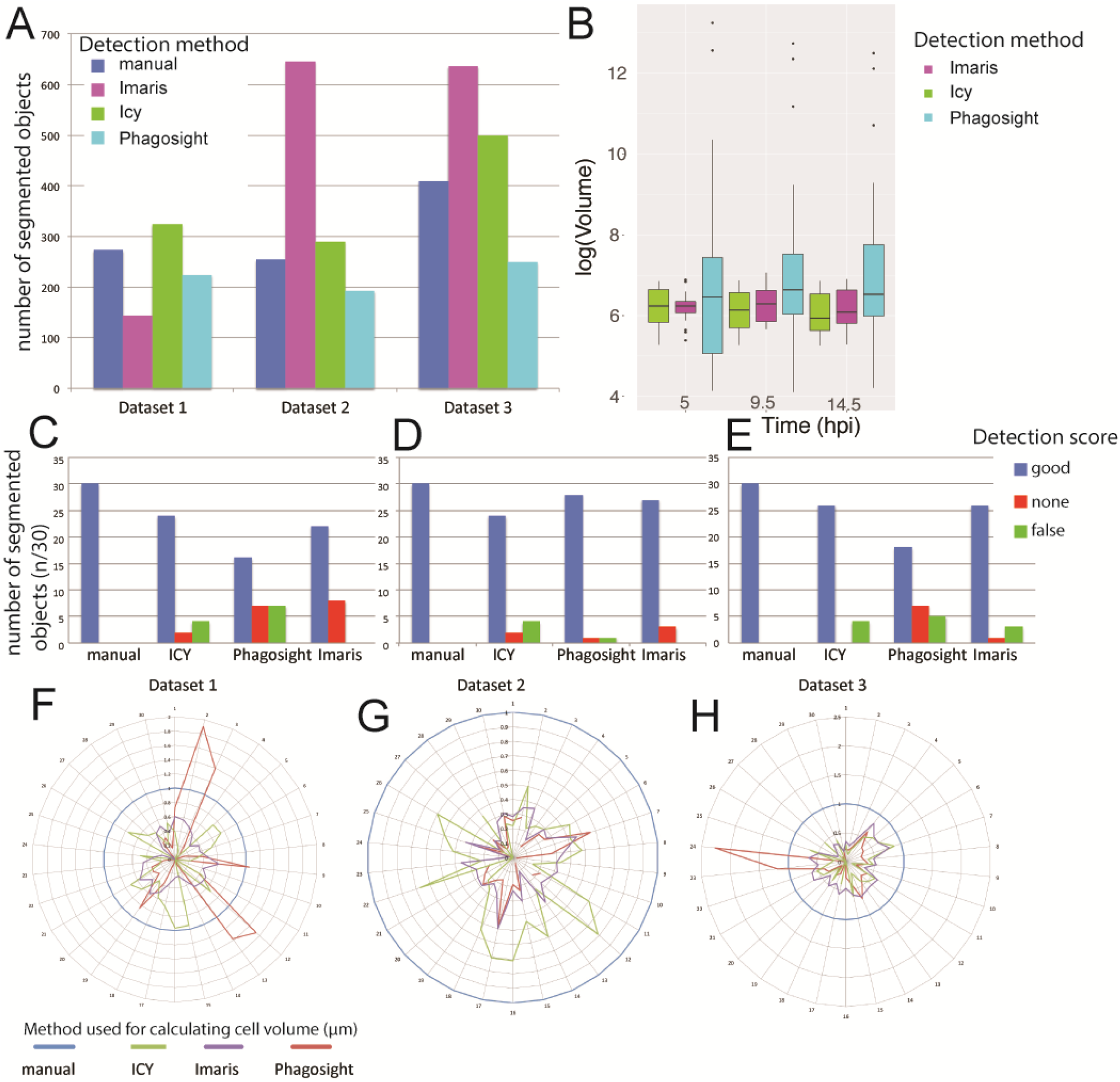
Evaluation of segmentation accuracy by Imaris, Icy and Phagosight relative to a ground truth. The number of segmented objects by Imaris (pink), Icy (green) and Phagosight (cyan) were compared to a manually defined ground truth (dark blue) from datasets 1-3 (A). The volumes of the segmented objects from dataset 1 were compared relative to a ground truth by plotting average values with standard deviation at 5, 9.5 and 14.5 hours post injury (hpi) for each method (B). The effectiveness of each method for detecting a selected cohort of 10 cells at 3 time-points (30 objects) was scored as good (blue), none (red) or false (green) for each dataset (C-E). The volumes of the selected 30 objects segmented by each method were compared by star plots in which the outputs of the three programs were normalised to the ground truth (blue) for each dataset (F-H).

We then assessed how well each program could identify the ten cells of interest that we had defined by manual annotation from each dataset. Datasets 1-3 were segmented using optimal parameters then the detection of the cells at the three defined time-points was defined as good (a single cell), poorly detected (cell detected, but not resolved as a single object or merged with another cell) or not detected. Each program showed differences in their ability to segment cells from the individual datasets (Figure 2C-E). Imaris and Icy were more effective at segmenting cells than Phagosight for datasets 1 and 3, but not dataset 2. Imaris had a higher error rate than Icy, which was more conservative and failed to segment cells that were poorly detected.

To evaluate the accuracy of segmentation by Icy, Imaris and Phagosight we compared volumes against the manually defined ground truth. Focusing on the ten cells of interest at one time-point, we noted that overall, all three programs failed to segment the entire cell volume relative to manual segmentation (Figure 2F-H). Icy was the most effective at segmenting cells from the datasets overall, although Imaris showed more consistency. In contrast, Phagosight showed relatively variable results. Again, this differed between datasets, but we consistently observed inappropriate segmentation by Phagosight, as several cells were segmented with much larger volumes than the ground truth.

### Analysing cell migration

In order to accurately describe cell behaviour during tissue repair, it is necessary to track them accurately through a four-dimensional space. Given the time-intensive nature of manually annotating and tracking cells an automated method is necessary. Icy, Imaris and Phagosight use different approaches to track cells: the Spot Tracking tool in Icy evaluates past and futures frames to predict the position of a cell over time; Phagosight employs a keyhole algorithm that makes predictions about cell movement from past and future frames in a restricted model of directional movement (Reyes-Aldasoro et al., 2008); Imaris offers several algorithms for tracking cells through the Surfaces tool.

To evaluate the accuracy of Imaris, Icy and Phagosight in tracking muSCs, we compared outputs from these programs to the ground state obtained by manually tracking cells (Figure S1A-C). Tracking for all three programs was performed using an optimally segmented labelled image for each dataset generated by the Icy tool HK Means. These acted as a reference image to launch the tracking process with the different software programs and allowed a direct comparison between them. To evaluate the error in these segmented reference images we assessed whether our selected ten cells were present at each time point (Figure S1D). This revealed that some of the cells were not identified at each time point, providing a potential source of error for subsequent tracking.

Tracking efficiency by Imaris, Icy and Phagosight was assessed by a manual scoring of how well they could identify tracks for the ten cells of interest segmented by HK Means. Effectiveness of tracking was scored by whether a segmented cell was linked between time-points (+1), if this was inappropriate (i.e. to another cell, −2) or was broken between subsequent time-points (−1).

Comparing the three programs for their ability to track cells reveals that on average, Icy was able to generate longer tracks and also generated fewer tracks than Imaris or Phagosight. The performance of Imaris was nearly as good as Icy and was better than Phagosight.

Accuracy of cell tracking by each program was evaluated by the Track Performance Tool. This compares tracking of cells from the same dataset by performing a pairwise comparison to a reference dataset (Chenouard et al., 2014). Using the manually tracked data as the reference, alpha, beta and Jaccard similarity coefficients of fit were obtained for each dataset (Figure S1E). These coefficients describe localisation and tracking errors when tracks are compared against the ground truth (alpha), and factoring in non-paired (spurious) tracks (beta); the Jaccard similarity coefficient describes the fit of the tracking data only. Using these coefficients, Imaris was observed to be the most effective at tracking cells relative to the ground truth for dataset 1 (Video 1) and 2, whereas Icy performed better for dataset 3.

### Cell Tracking Profiler: a semi-automated pipeline for segmenting and tracking cells

In order to optimally segment and track cells in a semi-automated manner, we built a Java-based pipeline that allowed tuneable segmentation and fast, accurate tracking together with an ability to manipulate and correct tracks. Outputs from Cell Tracking Profiler (CTP) include parameters of cell shape (volume, surface, sphericity, convexity) and movement (displacement, distance, directionality, instantaneous speed) for each cell over time (Figure 3A). To facilitate post-processing of date we wrote a MATLAB script, CTP2R, to process outputs from CTP and select regions of interest (ROIs) to analyse. CTP2R calculates mean squared displacement and directional autocorrelation of cells for one or two user-defined ROIs and writes values for each cell into a spreadsheet that can be opened and plotted in R. (Figure 3B).

**Figure 3:**
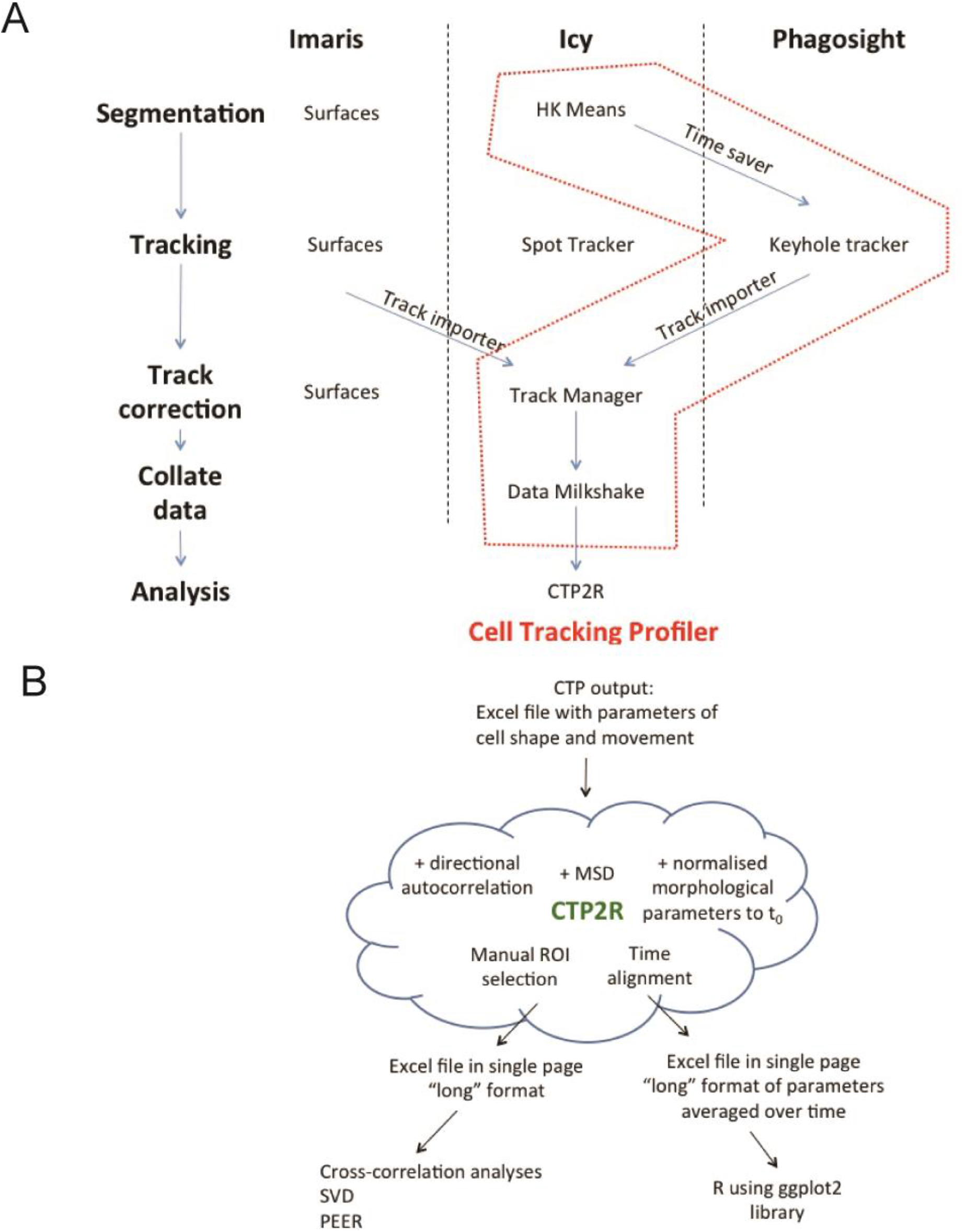
Cell Tracking Profiler (CTP) forms a practical workflow for managing cell segmentation and tracking and can feed outputs into a variety of tools for statistical analysis. CTP runs in Icy under ‘plugins’. Users define segmentation parameters for processing of datasets by HK-Means. Outputs, including images and measurements of cells were saved by ‘Time-Saver’ into two files. Objects are tracked by the Keyhole Tracker program in MATLAB runtime then imported into the Icy plugin Track Manager for editing. The Milkshake plugin then collates information from tracking with values of cell shape to produce an Excel sheet (A). This can be analysed using a MATLAB script, CTP2R, which can normalise data to time-point 0 (t_0_), generate values of directional autocorrelation, mean squared displacement (MSD), select a region of interest (ROI) and average parameter values over time (B).

Our criteria for building CTP were the need for: 1) speed of segmentation and tracking, 2) ability to modify cell tracks, 3) linking of cell shape and movement parameters, 4) freely available code on a robust platform. Based on these requirements, CTP was built to run from Icy in a Java environment. It incorporates a modified version of HK Means (myHKMeans) that segments cells from 3 separate channels with user input to define optimal values suitable for the range of cell shape. Cell shape parameters of segmented cells are extracted, then, cells are tracked using Keyhole Tracker. Tracked cells were evaluated by eye using the ‘Track Manager’ tool and corrected manually, with a final output file being produced containing parameters of cell movement and shape by utilising a new plugin called Milkshake (Figure 3A).

We investigated the ability of CTP to segment and track all cells in datasets 1-3 relative to an uninjured control animal (dataset 0, Video 2). Following editing of tracks using Track Manager cells could be tracked throughout the movie and corresponding values of shape and movement obtained for statistical analysis.

### Analysis of muSC behaviour using CTP

A key requisite of CTP was that it could track cells and generate outputs of cell shape that allow statistical analysis. In order to determine whether there were significant changes to cell behaviour and shape in response to injury we selected regions of interest using CTP2R. This allowed us to compare values for cells in injured myotomes relative to those in uninjured myotomes (Figure 4A). There were significant differences in parameters of shape (volume, surface area, sphericity, convexity) between cells in injured myotomes from datasets 2 and 3 relative to cells in an uninjured control animal captured in dataset 0 (Fig 4B). Similarly, tracking of cells within injured myotomes from datasets 2-3 showed a more directional and rapid movement than cells from uninjured animal in dataset 0 (Fig. 4C). In contrast, dataset 1 showed no significant differences in cell shape (roundness) between cells in injured myotomes relative to those in dataset 0 and did not display any differences in cell movement.

**Figure 4:**
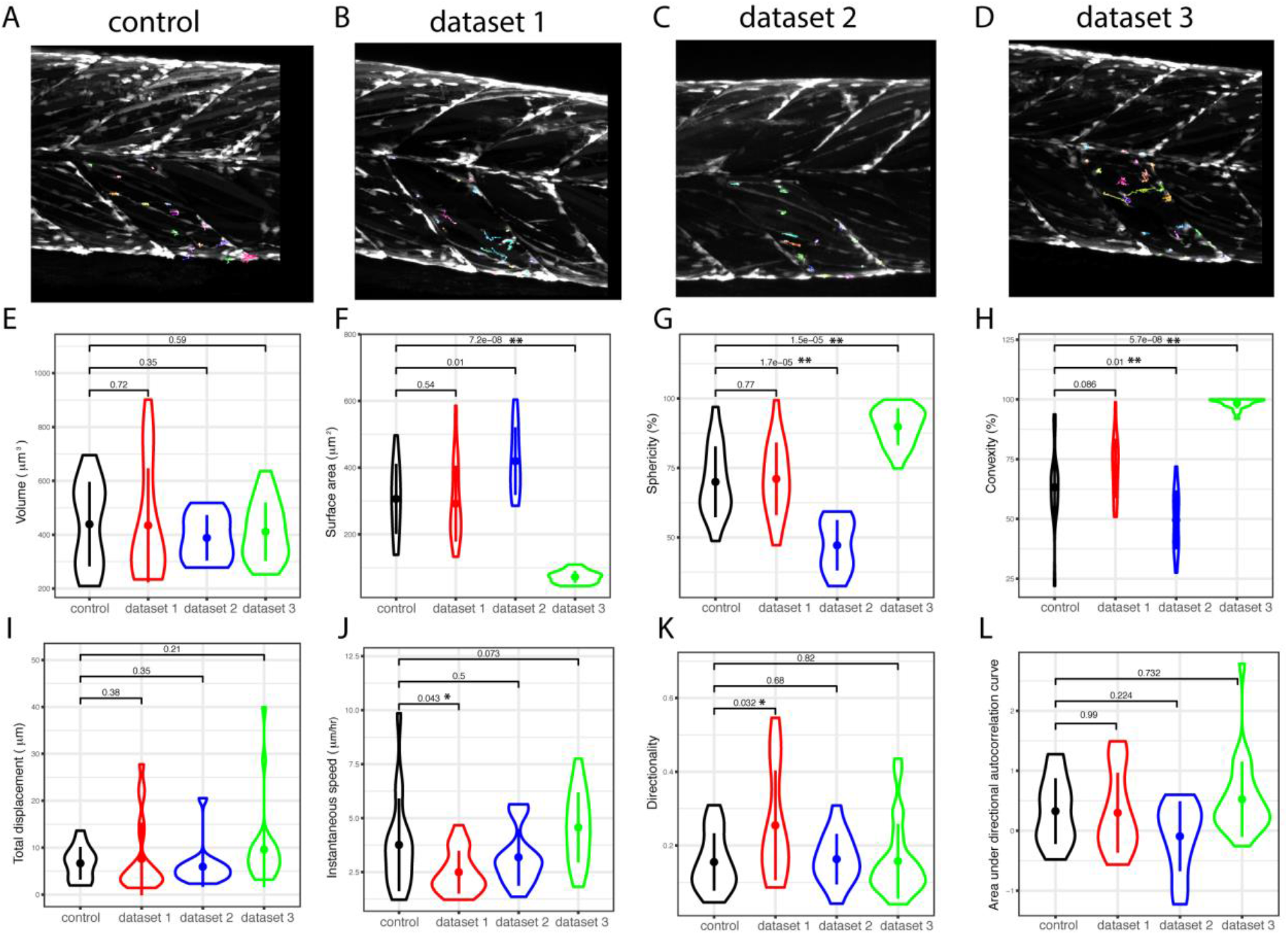
Tracking and processing of datasets 0, 1, 2 and 3 from injured larvae using CTP 2.0. Time-lapse images from injured (datasets 1-3) and an uninjured larva (control, dataset 0) (A-D) were processed by CTP 2.0 using optimal parameters for segmentation and tracks corrected using Track Manager. Cells in the 11^th^ ventral myotome were selected using CTP2R (coloured tracks) and measures of shape (cell volume, surface area, sphericity, convexity, E-H) and movement (total displacement, instantaneous speed, directionality, I-K) plotted as Violin plots. Directional autocorrelation was compared between datasets by by measuring the area from beneath the curve of plotted values (L). Significance of difference between datasets for each parameter was calculated using a 1-way ANOVA and pairwise comparisons performed using a Tukey posthoc test with significance denoted as p<0.05 (*) or p<0.01(**).

In order to determine whether CTP was able to extract sufficient detail of cell shape and movement to reveal the effect of a perturbation, we analysed the behaviour of cells exposed to RhoA kinase (ROCK) inhibitors (Fig. 5A-C). A comparison of cell shape and movement revealed clear differences between animals treated with DMSO vehicle control (dataset 4, Video 3) relative to those treated by 15 µM Y-27632 (dataset 5, Video 4) or 10 µM ROCKOUT (dataset 6, Video 5, Table S1). Measures of cell shape (volume, sphericity, convexity, roundness) were significantly altered by inhibition of ROCK (p<0.01, Figure 5D). Some parameters of movement (displacement, directionality, directional autocorrelation, p<0.001) were also strongly reduced by inhibition of ROCK, although instantaneous speed was less affected (p<0.01, Figure 5E). This suggests that inhibition of ROCK affects muSC shape and perturbs directed cell movement. RhoA signalling is a known regulator of myosin II contractility and hence can control cell migration. To determine whether myosin II function is important for muSC responses to injury we measured cell responses in the presence or absence of 10µM blebbistatin (datasets 7-12, Figure 5F-G, Videos 6-7). We noted that in uninjured myotomes addition of blebbistatin altered cell shape but did not affect cell movement (Fig 5H-I). In the presence of an injury, parameters of cell movement, but not shape, were significantly affected (p>0.05, Table S2).

**Figure 5:**
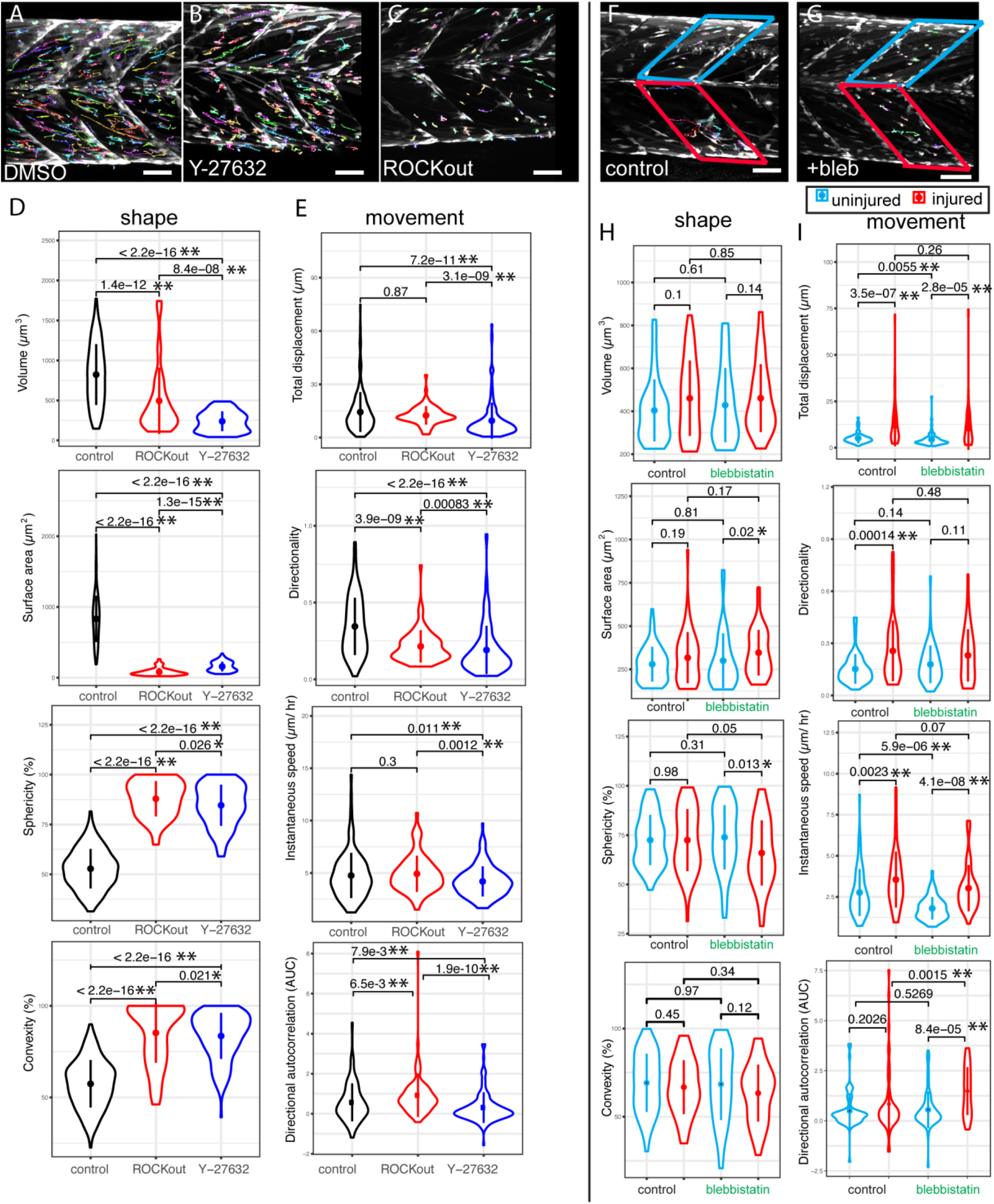
Outputs from CTP from analyses of datasets generated from animals exposed to small molecule inhibitors of ROCK (A-E) or of myosin II (F-I). GFP+ cells were tracked (coloured lines) from time-lapses of pax7a:egfp larvae exposed to DMSO (A), Y-27632 (B) or ROCKOUT (C). Violin plots of cell shape (D) and movement (E) were generated from CTP outputs after correction of tracking and significance of differences tested by Kruskal-Wallis tests and pairwise comparisons performed with a Dunn post-hoc test. Tracking of GFP+ cells in injured (red box) and uninjured (green) myotomes of *pax7a:egfp* larvae in the absence (F) or presence of 10 µM blebbistatin (G) was performed (n=3 for each condition). Violin plots of shape (H) and movement (I) were generated from CTP outputs for cells in injured myotomes (red) and adjacent uninjured myotomes (blue) in larvae exposed to blebbistatin or water. Significance of differences in parameters of shape and movement were evaluated using Kruskal-Wallis tests and pairwise comparisons tested by Dunn post-hoc tests with comparisons showing p<0.05 (*) and p<0.01(**) indicated. Scale bars: 50 µM (A,B,C,F,G).

### A statistical approach for defining muSC responses to injury

Having identified a number of muSC shape and movement parameters that are altered due to specific interventions, we wished to identify a statistical approach that would allow us to discriminate changes to cell responses when multiple parameters are simultaneously affected. Importantly, we also wanted to account for differences relative to time, as many parameters changed over the timecourse of the experiment. Principal component analysis (PCA) revealed that cell shape parameters were strongly affected by inhibition of ROCK (Figure 6A). Variables representing parameters of shape show closest proximity to the circle of correlations and hence are the most important in explaining the variability in the dataset. In contrast, PCA analysis of cells exposed to blebbistatin in injured or uninjured muscle showed a clear segregation into three groups including variables for parameters of roundness (convexity, sphericity, roundness), flatness (surface area, maximum Feret diameter) or movement (Figure 6B). All three groups of variables seem equally important for explaining changes as a consequence of blebbistatin treatment and injury relative to their distribution.

**Figure 6:**
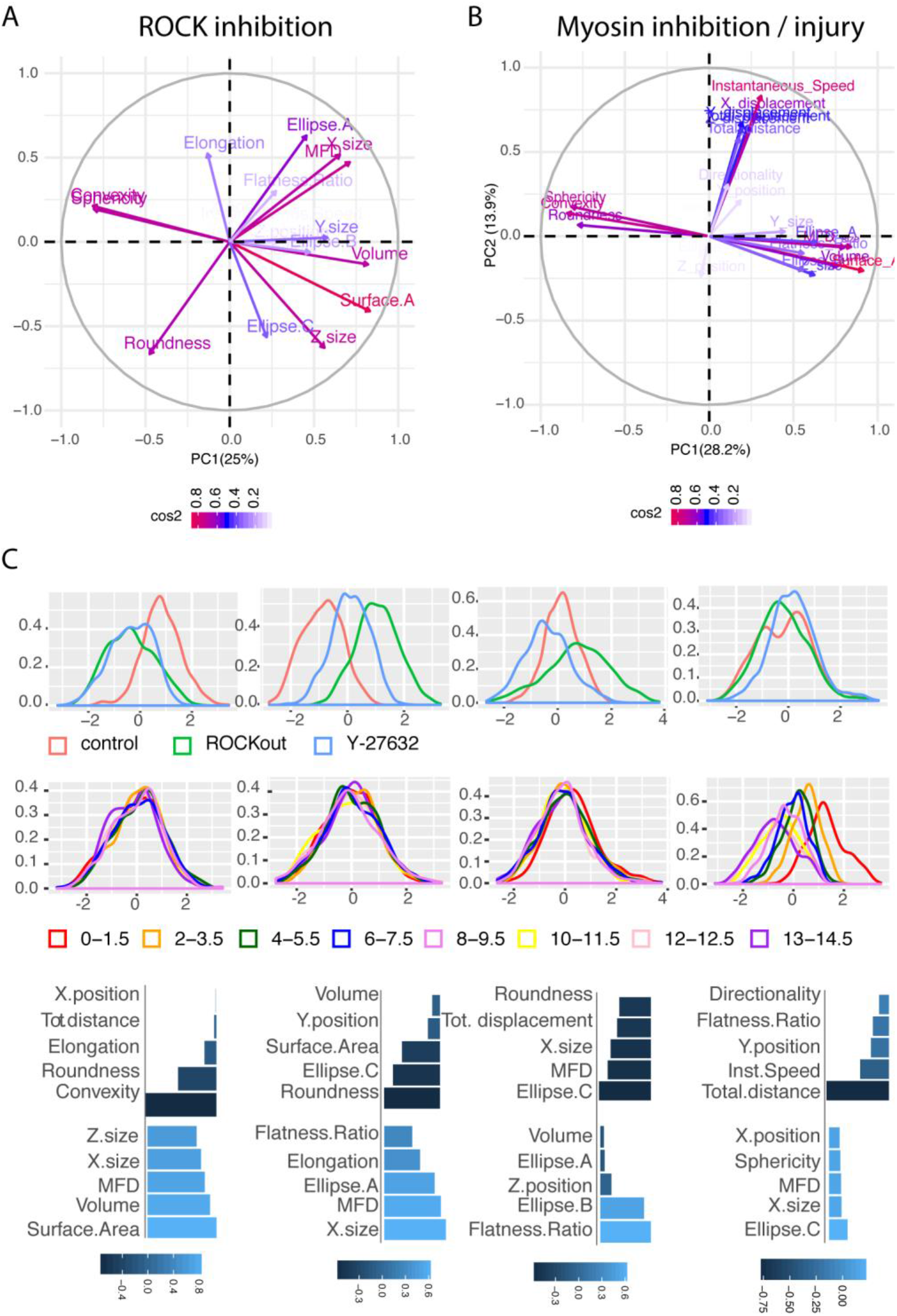
Dimensionality reduction analyses of CTP outputs from larvae treated with ROCK inhibitors or by blebbistatin in the presence or absence of injury. PCA variable plots for datasets 4-6 (A, ROCK inhibitors) and 7-12 (B, obtained from injured or uninjured myotomes in which animals were exposed to blebbistatin) revealing relationships between variables. Positively correlated variables are grouped together and negatively correlated variables are positioned on opposite sides of the plot origin (opposed quadrants). Colours correspond to the cos2 value (the squared loadings for variables - if a variable is perfectly represented by only two components, the sum of the cos2 is equal to one) revealing the importance of a given principal component for behaviour of a variable. Distance between variables and the origin indicates the quality of the variables for explaining the data on the factor map. PEER plots showing the distribution of values for the synthetic factors obtained with PEER dimensionality reduction as a function of density (y axis) relative to 0 (y axis) for datasets 4-6 of larvae treated with ROCK inhibitors (C). The effect of the external perturbation (drug, C) and the effect of the time (hours after commencement of time-lapse, D) are shown by changes to the distribution of the factor. Loading of phenotypic raw features on each factor reveals how much changes to each parameter contributes to the factor with negative and positive values (shaded blue scale) reflecting the importance of that parameter (E).

In order to explore how ROCK inhibition and technical or biological components (covariates) contributed to the observed variation in cell phenotypes, we applied a dimensionality reduction approach called Probabilistic Estimation of Expression Residuals (PEER) (Stegle et al., 2012). PEER was originally implemented for gene expression data and then applied to multidimensional reduction of phenotypic data (Vigilante et al., 2018). It is a collection of Bayesian approaches that takes as input measurements and covariates and then extract factors that explain hidden portions of the variability. This is fundamental because unobserved, hidden factors, such as cell culture conditions can have an influence on large numbers of cells and many studies have demonstrated the importance of accounting for hidden factors to achieve a stronger statistical discrimination signal (Kang et al., 2008; Leek and Storey, 2007; Stegle et al., 2010).

Using PEER, 21 phenotypic measurements and covariates (i.e. time, cell identity and presence or absence of drug) were used as inputs; factors that explain portions of the variance were generated and used to explore cell phenotypes. The optimal number of PEER factors was set to 5 according to the automatic relevance detection (ARD) parameters, which reveal those dimensions needed to model the variation in the data. PEER Factors that revealed cell responses to ROCK inhibition (Factor 1) and discriminated between responses to Y-27632 and ROCKOUT (Factors 2 and 3, Figure 6C) were identified. By mapping the PEER factors against time as a covariate, we noted found that these factors show an altered distribution over time, suggestive of changes to cell shape or behaviour across the timecourse of the experiment (e.g. Factor 4 shows a changing distribution from 0-1.5 h relative to 13-14.5 h, Fig. 6D). Mapping of the contribution of shape and movement parameters to PEER factor 1 revealed that cell shape (volume, cell shape) can be used to show differential responses to Y-27632 and ROCKOUT relative to control cells, but cell displacement and instantaneous speed did not (Figure 6E). In contrast, Factor 2 segregated all three conditions and was influenced most strongly by cell shape (mean Feret diameter, x-size, elongation, Figure 6E). This suggests that cell shape is most different between the three conditions and concurs with the analyses of single parameters (see Figure 5).

We applied a similar approach to probe how the responses of muSCs to injury were affected by blebbistatin. We identified five PEER factors that can mostly explain the variance of the data and found that factor 2 identifies differences in cell responses from injured and uninjured myotomes, or in the presence or absence of blebbistatin (Figure 7A). Factor 2 is strongly influenced by cell movement (instantaneous speed, XYZ displacement), indicating the importance of both injury and blebbistatin on cell migration (Fig. 7D). In contrast, factor 1, which has a strong cell shape contribution (area, volume, mean feret diameter) was not obviously altered by blebbistatin, but did respond to injury (Figure 7A, D). Strikingly, these PEER factors showed differing responses to the combination of blebbistatin and injury over time (Figure 7B). Factor 1 revealed that the addition of blebbistatin resulted in injured cells assuming a profile at later times that was more similar to cells not treated by blebbistatin at early stages (Figure 7C, compare B_I and NB_I). In contrast, factor 2 showed a difference in the response of cells to injury in the presence or absence of blebbistatin at late time points (Figure 7C,E, compare B_I and NB_I).

**Figure 7:**
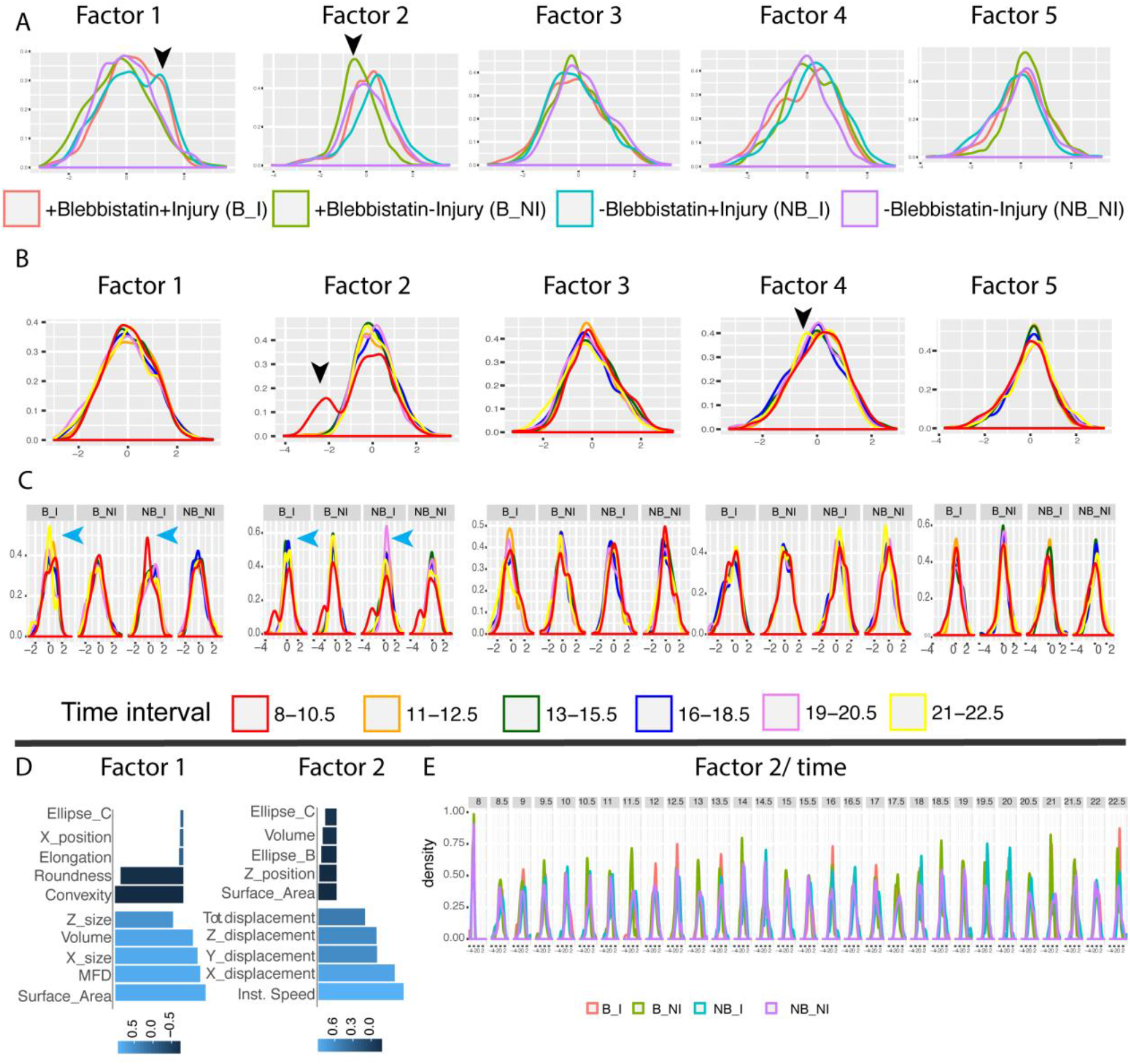
PEER factor distribution for datasets 7-12 relative to independent variables blebbistatin, injury and time. PEER Factor 1 distribution differs between cells in injured myotomes dependent on whether blebbistatin is added or not (arrowhead to blue line, A). In contrast, Factor 2 shows clear differences due to addition of blebbistatin in an absence of injury (arrowhead to green line, A). Factors 2 and 4 show different distributions as a consequence of time (hours post injury, hpi) with early (red, 8-10.5 hpi) and later (yellow, 21-22.5 hpi) time intervals affecting factors differently (arrowheads, B). Mapping of time relative to the four classes of manipulations (Blebbistatin +Injury: B_I, Blebbistatin No Injury: B_NI, No Blebbistatin +Injury: NB_I, No Blebbistatin No Injury: NB_NI) for each factor (C). The distribution of factor 1 is affected by time as the peak distribution for class B_I occurs between 21-22.5 hours post injury (yellow line) and for class NB_I at 8-10.5 hours post injury (red line, blue arrowheads). Similarly Factor 2 shows a temporal shift in distribution between B_I and NB_I (blue arrowheads). Loading of phenotypic raw features on factor 1 and 2 reveals how much the change to each parameter weights the factor (D). Negative and positive values are represented by a scale (blue) with pale blue showing a positive contribution and dark blue indicative of not contributing to the factor. The distribution of factor 2 changes for each class (coloured line) over time (hours after injury) reflecting differences in the responses of contributing parameters (E).

To better understand how specific parameters are related we asked whether there are pairs of parameters that show a correlated change in response to injury or addition of blebbistatin. We found positive correlations between parameters of cell movement or shape regardless of blebbistatin treatment or injury including those between i) surface area and volume, ii) surface area and maximum Feret diameter, iii) sphericity and convexity, iv) convexity and roundness (Figure 8). Correlations between measures of movement were not strong suggesting considerable heterogeneity between cells. In an absence of an injury, correlations between parameters of cell shape were often reduced by addition of blebbistatin (Table S3). A negative correlation between directionality and surface area (−0.08) becomes a positive correlation in the presence of blebbistatin (0.08). Correlations between parameters were often low and it was therefore difficult to extrapolate from these how cell behaviour might be affected by injury or blebbistatin. By visualising measures of significance with the correlation coefficient in a heat map several differences due to injury or addition of blebbistatin became more discernable (Figure 9). Correlations between total displacement and elongation and between total distance and surface area occur in cells responding to injury. These are altered in the presence of blebbistatin, indicating that the relationship between cell shape and movement are altered by inhibition of myosin II function as predicted from a variety of in vitro studies.

**Figure 8:**
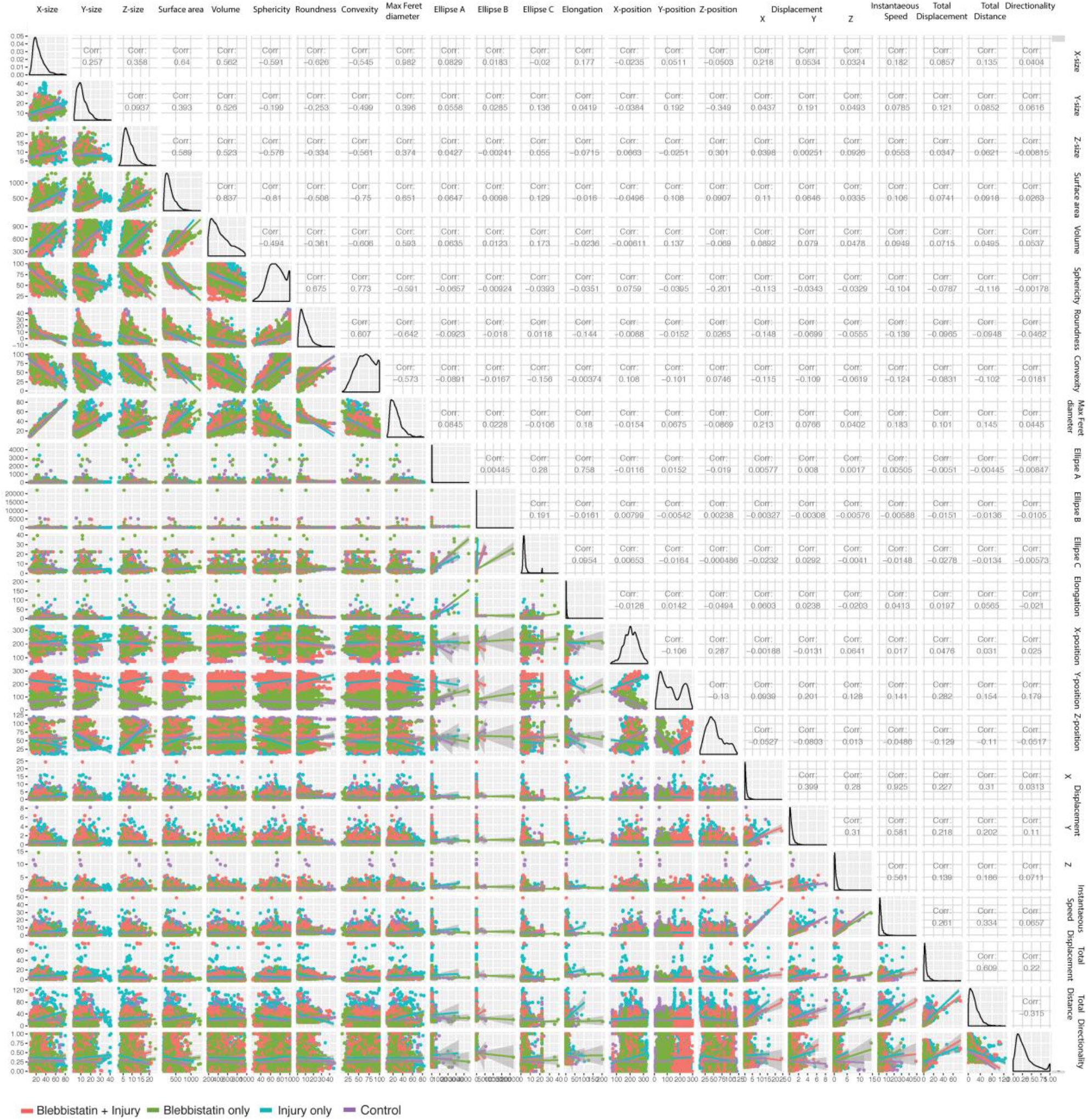
Correlation analyses of different phenotypic measurements of cells from datasets 7-12 (I). Parameters of shape and movement (X and Y axes) were compared across different conditions that were separated into four classes: 1) blebbistatin + injury (red), 2) blebbistatin, no injury (green), 3) no blebbistatin, injury (blue), 4) no blebbistatin, no injury (purple). A line of best fit is shown for all conditions and a corresponding correlation coefficient for the two parameters being compared.

**Figure 9:**
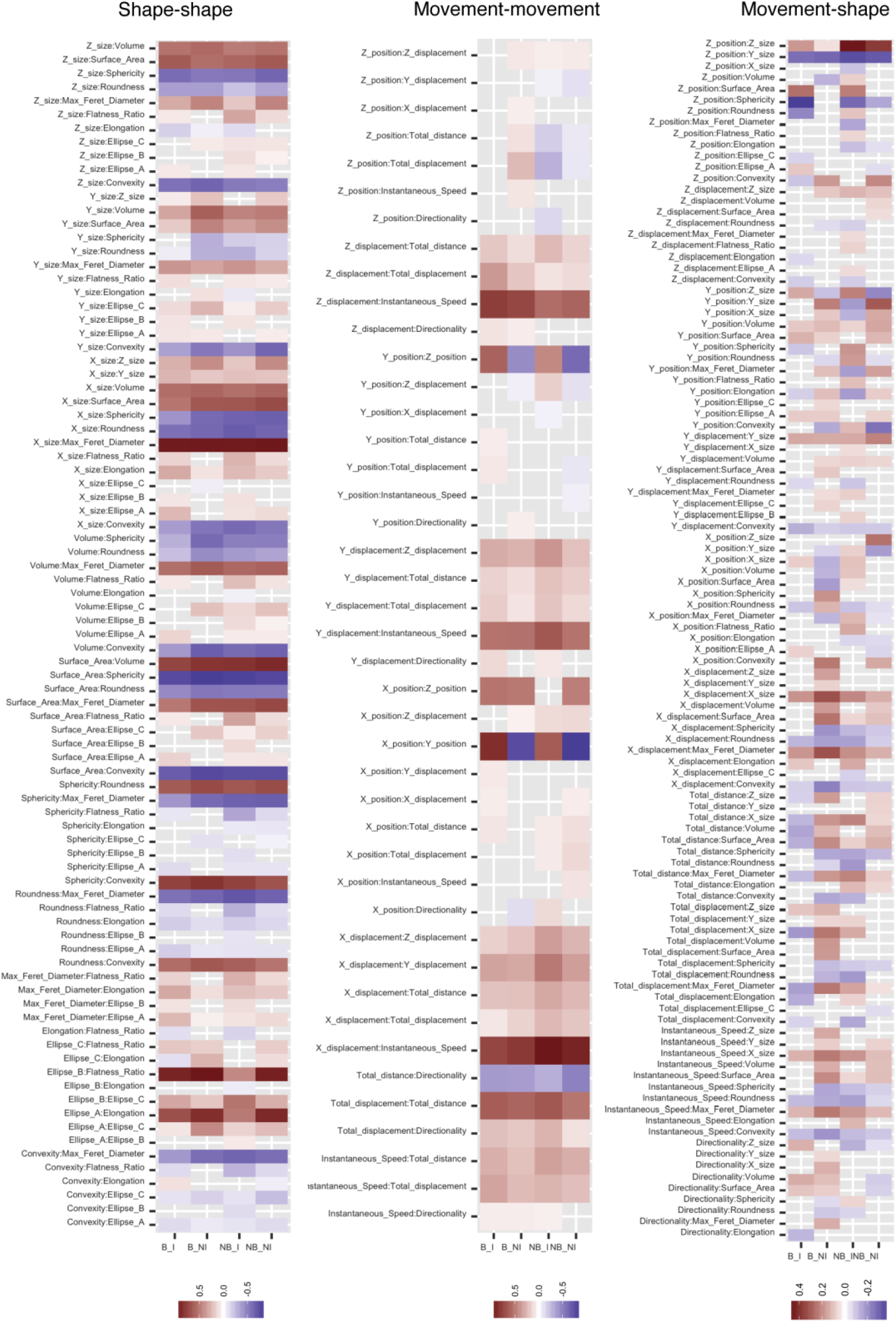
heatmaps showing correlation values for all parameters of shape and movement from datasets 7-12 when compared against each other. Colour represents the correlation value for each class (Blebbistatin +Injury: B_I, Blebbistatin –Injury: B_NI, No Blebbistatin +Injury: NB_I, No Blebbistatin –Injury: NB_NI) and grey boxes correspond to non-significant correlations (p<0.05).

In order to understand whether different variables interact to modify specific parameters of cell shape or movement we employed multiple linear regression to test the relative importance of key variables (injury, blebbistatin, injury, dataset, time) and interaction effects between myosin II inhibition and the induction of an injury (Table S4, Figure 9). No strong associations were identified between dataset origin or with time with changes to parameters of shape or movement. Blebbistatin exposure was strongly associated with changes to parameters of curvature (sphericity, roundness and convexity) as suggested from analyses using PEER. The interaction between blebbistatin and injury was also important for the changes in parameters of curvature although it generally showed an opposite association. Taking convexity as an example, the coefficient of regression for blebbistatin is 0.78, for injury is −0.08 and the interaction between blebbistatin and injury is −0.203 (Table S4). These are all significant associations and tests of residuals and leverage indicate a good fit for the regression model. A general pattern we observed is that correlation coefficients for blebbistatin and injury showed opposite responses relative to parameters of shape, suggesting these variables induce opposing responses to cell shape in muSCs. A similar difference in the association between blebbistatin or injury was observed for parameters of movement (displacement, instantaneous speed). The coefficient of regression for the interaction between blebbistatin and injury was also significant for many parameters and generally showed a similar relationship as seen for the injury variable. This indicates that although the response of cells to injury is strongly blunted by addition of blebbistatin, they still show a profile reflecting a migratory behaviour.

## Discussion

Imaging cell behaviour in vivo or in 3D tissue engineering constructs faces several limitations including the need to often visualise a fluorescent signal at depth in tissue, autofluorescence of the tissue or limitations of the marker used to label the cell. These conspire to make analysis of such time-lapsed datasets challenging and can limit the ability to apply statistics in order to discern changes to cell behaviour. We find this a particular challenge in our goal of imaging fluorescently labelled muSCs during tissue regeneration in zebrafish.

To measure cell shape and movement from time-lapsed movies we have developed an image analysis approach that allows us to segment cells and extract measures in a semi-automated manner. An evaluation of available programs for segmentation and cell tracking revealed that combining the HK Means method for threshold classification of pixels with a keyhole tracking algorithm provided a rapid and effective method for analysing the dynamic movements of muSCs. We incorporated these programs into Cell Tracking Profiler, a package that runs in the freely available Icy platform and allows us to identify the best conditions for accurate segmentation of cells, to track them over time and then manually correct this tracking data. The strength of our approach is that we are able to analyse cell behaviour from complex 4D datasets in which there are few markers available for defining cell shape or identity, making segmentation problematic. Further, by being able to adjust the tracking information manually after the tracking is complete, we are able to effectively define cell migration with no need for extensive optimisation of the tracking process. We then explored a variety of statistical methods for analysing changes to cell shape and movement. A combination of dimensionality reduction approaches (PCA, PEER), correlation and multiple regression analyses were used to reveal opposing effects of injury and blebbistatin on muSC behaviour.

Several packages have been described for analysis of live cell imaging data, including plugins that run in ImageJ/ Fiji or that are written in a variety of programming languages (Driscoll and Danuser, 2015; Hilsenbeck et al., 2016; Rajasekaran et al., 2016). Using these tools requires an understanding of the assumptions underlying the package or some fluency in programming. In contrast, our analysis pipeline utilises Icy as the main interface, making it user-friendly and amenable to modifications as it is open source and does not require programming skills. As proof of principle, we show that user-defined settings for defining segmentation and allowing correction of tracked cells allows a medium throughput analysis of time-lapsed data.

Our comparisons of commonly used software, such as Imaris and Icy, revealed limitations in their accuracy of segmentation and tracking relative to a manually annotated ground truth. One approach to increase the accuracy of analysis is to improve image acquisition parameters. However, this may not be always desirable, due to potential bleaching or phototoxicity of the sample, an inability to acquire information at sufficient resolution rapidly enough, insufficient signal of the fluorescent signal or high background. Our solution to these challenges was to build an analysis pipeline that allows user-led correction and employs statistical models to evaluate the importance of contributing variables. PEER has been previously used to investigate relationships between cell behaviour and gene expression for cells cultured *in vitro* (Stegle et al., 2012). We demonstrate the utility of PEER for dissecting the complexity of cell behaviour *in vivo* and to identify which parameters of cell behaviour are sensitive to specific perturbations. Corroboration of trends observed using PEER came from cross-correlation analyses and multiple linear regression, which allowed us to examine how multiple perturbations affect cell behaviour. Utilising this methodology for classifying the response of a motile stem cell population in response to a drug provides a powerful tool for identifying small molecule regulators of regeneration in complex animal models.

A central facet of our work is to understand how resident stem cell behaviour is regulated in response to injury. Measurements of Mean Squared Displacement or fitting to a model of anisotropic persistent random walk revealed that muSCs move towards muscle injury in a directional manner. This movement likely reflects a discrete mode of behaviour that requires cells to respond to a changing external environment as tissue remodelling occurs. As the parameters used to define cell shape do not represent complex structures it is not possible to identify features including protrusions, blebs or asymmetric features by CTP. This is a potential limitation in describing cell shape that will overlook informative features. Nonetheless we were able to define ROCK-dependent changes to muSC shape and movement which appeared to be uncoupled when examined by PCA and PEER. This segregation may reflect differential requirements for ROCK in controlling cell behaviour by modifying localised actinomyosin contractility, regulating formin during actin bundling, or controlling myosin and actin flow (Konigs et al., 2014; Ridley, 2015; Ruprecht et al., 2015).

ROCK phosphorylates myosin II and myosin II dependent functions have been described to include regulation of actin flow and stabilisation of focal adhesions, critical for mesenchymal cell migration (Aguilar-Cuenca et al., 2014). We noted that migration speed and shape of muSCs are affected by blebbistatin similar to ROCK inhibition. However, based on their differing contribution to the PEER factors, these parameters respond differently to blebbistatin treatment compared to cells treated with ROCK inhibitor. This implies that inhibition of RhoA signalling and myosin II do not result in similar phenotypic outcomes in muSCs, reflecting their diverse roles in controlling cytoskeletal organisation and cell shape.

A variety of myosin II independent cell movements have been described that are dependent on actin flow (Lomakin et al., 2015). Further, many cells are able to switch their mode of cell migration to myosin II-independent modes in both 3D artificial and *in vivo* environments (Liu et al., 2015; Ruprecht et al., 2015). Our observations indicate that muSCs are able to migrate when myosin II activity is inhibited, revealing a similar plasticity in how muSCs can migrate *in vivo*.

In summary, we show that by creating a user-defined semi-automated process for dataset segmentation and cell tracking, we can obtain biological signatures for cell responses to specific molecular manipulations. CTP runs in Icy, an open source platform, providing great scope for further development and integration with other analysis suites. Finally, we demonstrate that using several statistical approaches reveals trends and associations between parameters of cell shape and movement that are differentially affected by multiple variables. Our approach, combining CTP with a variety of statistical tests therefore provides the means for dissecting the consequences of multiple inputs on cell behaviour in complex 4D datasets.

## Methods

### Zebrafish husbandry and imaging

Transgenic 7 days post fertilisation (dpf) *Tg*[*pax7a:egfp*]*; pfeffer* zebrafish larvae were injured as previously described (Knappe et al., 2015) and treated with small molecule inhibitors. Images were acquired at 1-5 µm intervals every 30 minutes using a Nikon C1 confocal microscope equipped with an Argon-ion 488nm laser and a 20x water-dipping objective (NA = 0.7) under a constant temperature of 28.5 °C. All animal work was performed according to local and Home Office regulations under project license PBC5F9B13 with reference to the ARRIVE guidelines for use of animals in scientific experiments (Kilkenny et al., 2010).

### Image Analysis

Time-lapse image sequences were processed by ImageJ to remove movement artifacts using the plugin ‘Correct for 3D drift’ (http://fiji.sc/Correct_3D_drift) and, if necessary, cropped to focus only on relevant regions. No other manipulations were performed to the images prior to analysis. Datasets were selected for analysis that were considered representative and were obtained from larvae showing no obvious signs of ill health. Parameters used for Imaris 8.0 (Bitplane AG), Icy 1.7.3 (de Chaumont et al., 2012) and Phagosight (Henry et al., 2013) are the optimal values of segmentation or cell tracking as defined relative to the ground truth. The ground truth dataset was obtained by: i) cells were counted by Cell Counter (Image J/ Fiji) at three time points for each time-lapse, ii) regions of interest (ROI) were defined for 10 cells per dataset by drawing around the cell in each slice of the z-stack, then merging to generate a 3D object. iii) the selected 10 cells were tracked over time using MTrackJ (Image J/ Fiji). Optimal values for segmentation were identified by i) counting the number of objects identified, ii) scoring the identified objects against the manually defined subset of 10 cells at 3 different points in the time-lapse, iii) evaluating the size of the segmented objects relative to the manually annotated ground truth. Parameter values that best identified cells from the ground truth as discrete objects were selected as optimal.

Cell movement was scored by comparing automatically generated values relative to the manual ground truth. A ground truth was established by tracking the selected 10 cells for each dataset using MTrackJ (Image J/ Fiji). Cell tracking was performed using Surfaces in Imaris, Spot Tracker in Icy and by employing a keyhole tracking algorithm in Phagosight. Measures of tracking used for calculating efficiency were i) the number of tracks identified, ii) the length of continuous tracks, iii) the number of times a track was interrupted, lost or falsely connected. Optimal parameter values were defined as those i) that resulted in the longest contiguous tracks for cells defined in the ground truth, ii) that resulted in the fewest number of false connections (see Supplementary note). Determination of the segmentation and tracking efficiency of Imaris, Icy and Phagosight were performed by calculating alpha, beta and Jaccard co-efficients using the Track Analysis tool in Track Manager (Chenouard et al., 2014).

### Software development

*CellTrackingProfiler* was generated using the Protocols tool in Icy. It is a semi-automated tool allowing segmentation and tracking of cells from 4D datasets. *CellTrackingProfiler* incorporates a modified version of the Icy segmentation plugin *HK-Means* (Dufour et al., 2008), *Keyhole Tracker* (the tracking algorithm from Phagosight was modified to run in Java through a MATLAB Runtime (Henry et al., 2013)) and scripts for extracting information about cell shape, position and movement. Manual correction of tracked cells is then achieved using the Icy *Track Manager* plugin before the corrected cell tracks are linked to information about cell shape by the *Milkshake* tool. The output is a tabulated file that can be opened by spreadsheet managers, such as Excel (Microsoft Corporation) or imported into other image analysis programs. This file describes several parameters of cell shape and movement, including volume, surface area, convexity, sphericity, x/y/z position, distance, displacement, speed for each segmented cell in the dataset for each channel.

To post-process and reformat the data for statistical analysis, the *CTP2R* application was written in MATLAB and compiled as a standalone executable using the MATLAB compiler. The script allows for selection of specific ROIs and additionally calculates directional autocorrelation and the mean squared displacement for each cell (equations from Gorelik and Gautreau, 2014 (Gorelik and Gautreau, 2014)). This post-processed data is then reformatted into a long, single sheet format for easy plotting using the ggplot2 library in R. Normalisation of data was performed by comparing each value to the value for that parameter at t=0 for the time-lapse in question.

To generate a single value to represent the directional autocorrelation for each cell, the area under the curve was calculated within CTP2R using the trapezoidal rule. Tests of fit to models of isotropic or anisotropic persistent random walk (PRW, APRW) were performed using a MATLAB script as previously described in Wu et al, 2015 (Wu et al., 2015).

### Statistical Analysis

Analysis of cell shape and movement produced by CTP2R were performed using R and plotted using the ggplot2 library. Distribution of data was evaluated for normality by Shapiro-Wilks tests. Statistical tests for differences of cell shape and movement were performed by 1-way ANOVA, Wilcoxon-Mann-Whitney tests or Kruskal-Wallis tests as appropriate, dependent on distribution of the data.

Multivariate analysis was performed using principal component analysis (PCA), PEER (Stegle et al., 2012), autocorrelation and multiple linear regression in R. PEER (probablistic estimation of expression residuals) uses an unsupervised linear model to account for global variance components in the data and yields a number of factor components that can be used as synthetic phenotype in further analysis. We tested a wide range of parameter settings for the model (the *k* number), controlling the amount of variance explained by it. We ran PEER with the full pre-normalized dataset with the following parameters: K = 5; covariates = cell ID, drug, treatment and time; maximum iterations = 10,000. PCA analysis and the results visualization was performed using R software and FactoMineR and factoextra packages. Multiple linear regression analysis was performed using the lm() function and visualized through the ggfortify package. A p-value threshold of 0.05 was set to select the statistically significant coefficients.

## Supplementary Materials

**Figure S1:** Tracking efficiency of segmented objects by Imaris (A), Icy (B) and Phagosight (C) was evaluated using dataset 1 following segmentation by HK Means to ensure consistency.

**Table S1:** Statistical significance of differences in parameters of cell movement and shape between control (DMSO) treated, ROCKout treated and Y-27632 treated larvae.

**Table S2:** Statistical significance of differences in parameters of cell movement and shape between injured and uninjured myotomes of control and blebbistatin treated larvae.

**Table S3:** table of correlation values (corr) and p values from cross-correlation tests between parameters of shape and movement for each class of cells analysed from datasets of larvae with variables of blebbistatin treatment and injury.

**Table S4:** multiple regression models for shape and movement parameters from datasets of larvae with variables of blebbistatin treatment and injury.

**Video 1:** Time-lapse movie of 7 dpf pax7a:egfp larva following injury (dataset 1)

**Video 2:** Time-lapse movie of 7 dpf pax7a:egfp larva in absence of injury (dataset 0)

**Video 3:** Time-lapse movie of uninjured 3 dpf pax7a:egfp larva with DMSO (1%) added prior to imaging (dataset 4)

**Video 4:** Time-lapse movie of uninjured 3 dpf pax7a:egfp larva with Y-27632 (15µM) added prior to imaging (dataset 5).

**Video 5:** Time-lapse movie of uninjured 3 dpf pax7a:egfp larva in absence of injury with ROCKout (10µM) added prior to imaging (dataset 6).

**Video 6:** Time-lapse movie of injured and uninjured myotomes in 7 dpf pax7a:egfp larva prior to imaging (dataset 9)

**Video 7:** Time-lapse movie of injured and uninjured myotomes in 7 dpf pax7a:egfp larva treated with blebbistatin (0.8µM) prior to imaging (dataset 10).

## Acknowledgments

We thank Pia Cumine and Stefanie Knappe for help with generation of imaging data, Alexis Gautreau for advice on DiPer, Brian Stramer for helpful discussions and Ron de Bruin for assistance with Excel macros. LC was funded through the Erasmus+ Training scheme. RK was funded by grants from BBSRC (BB/P002390/1) and the Wellcome Trust (202852/Z/16/Z, 108111/Z/15/Z). JASL is funded by a City University doctoral scholarship. CCRA is partly funded by grants from the Leverhulme Trust (RPG-2017-054) and the Australian Research Council (DP170102235).

## Author Contributions

C.M. performed image and statistical analysis, wrote code and contributed to the writing of the paper. L.C. performed analyses of data and wrote code. A.V. performed statistical analyses. J.S.L. developed MATLAB code. C.C.A.R. developed MATLAB code and contributed to the writing of the paper. F.d.C. wrote code for Icy. A.D. wrote code for Icy.S.D. wrote code for Icy. J.C.O.M. provided computing infrastructure for development of Icy and contributed to the manuscript. RK designed and coordinated the project, performed image acquisition, analyses and writing of the paper.

There are no competing interests for any author of this work.

