## Supplementary material for "Cell Tracking Profiler: a user-driven analysis framework for evaluating 4D live cell imaging data"

Figure S1: Tracking efficiency of segmented objects by Imaris (A), Icy (B) and Phagosight (C) was evaluated using input as dataset 1 following segmentation by HK Means. Tracks for cells are overlain on a slice from a single time-point (A-C) and were compared against the ground truth generated by manual tracking. Scoring of tracking was performed by counting the number of track breaks, loss or inappropriate links for each program for all three datasets (D). Accuracy of tracking for each program was determined by calculating alpha, beta and Jaccard coefficients of fit (E) relative to the ground truth.
Scale bar 50 μm.

Table S1: Statistical significance of differences in parameters of cell movement and shape between control (DMSO) treated, ROCKout treated and Y-27632 treated larvae. Significance for each parameter between conditions was tested by a Kruskal-Wallis test followed by pairwise comparisons using a Dunn post-hoc test.

Table S2: Statistical significance of differences in parameters of cell movement and shape between injured and uninjured myotomes of control and blebbistatin treated larvae. Significance for differences in a parameter between conditions was tested by a Kruskal-Wallis test followed by pairwise comparisons using a Dunn post-hoc test.

Table S3: table of correlation values (corr) and p values from cross-correlation tests between parameters of shape and movement. Values are derived for each class of cells analysed from datasets of larvae with variables of blebbistatin treatment and injury. Classes are defined as: Blebbistatin +Injury (B_I), Blebbistatin –Injury (B_NI), No Blebbistatin +Injury (NB_I), No Blebbistatin –Injury (NB_NI). Significance of correlations between parameters is shown for uninjured animals in an absence of blebbistatin (controls) with $p<0.05$ (*) or $p>0.05$ (NS) and correlations are colour coded to show positive (yellow) or negative (orange) correlations.

Table S4: multiple regression models for shape and movement parameters from datasets of larvae with variables of blebbistatin treatment and injury. Changes to parameters of shape and movement were fitted against models incorporating independent variables of time, injury, addition of blebbistatin, the dataset or injury-blebbistatin interactions using multiple linear regression. The coefficient of fit and significance value is shown for each parameter relative to the variable tested. Correlations showing a significance of $p<0.05$ (*) and a coefficient of fit >±0.4 were selected and negative correlations (>-0.4) highlighted in orange and significant positive correlations in yellow (>0.4).

Video 1: maximum intensity projection of a confocal imaging time-lapse sequence of a 7 dpf pax7a:egfp larvae that has been injured (arrowhead) in the ventral myotome (dataset 1). Z-stacks were acquired every 20 minutes from 8 hours after injury (asterisk). Scale bar 50μm.

Video 2: maximum intensity projection of a confocal imaging time-lapse sequence of an uninjured 7 dpf pax7a:egfp larvae (dataset 0). Scale bar 50µm.

Video 3: maximum intensity projection of a confocal imaging time-lapse sequence of an uninjured 3 dpf pax7a:egfp larvae treated with 1% DMSO (dataset 4). Z-stacks were acquired every 20 minutes. Scale bar 50µm.

Video 4: maximum intensity projection of a confocal imaging time-lapse sequence of an uninjured 3 dpf pax7a:egfp larvae treated with 15µM Y-27632 (dataset 5). Z-stacks were acquired every 20 minutes. Scale bar 50µm.

Video 5: maximum intensity projection of a confocal imaging time-lapse sequence of an uninjured 3 dpf pax7a:egfp larvae treated with 10µM ROCKout (dataset 6). Z-stacks were acquired every 20 minutes. Scale bar 50µm.

Video 6: maximum intensity projection of a confocal imaging time-lapse sequence of a pax7a:egfp larvae that has been injured (asterisk) in ventral myotome 11 (dataset 9). Scale bar 50µm.

Video 7: maximum intensity projection of a confocal imaging time-lapse sequence of a pax7a:egfp larvae that has been injured (asterisk) in ventral myotome 11 (arrowhead) and treated with 0.8µM blebbistatin (dataset 10). Scale bar 50µm.

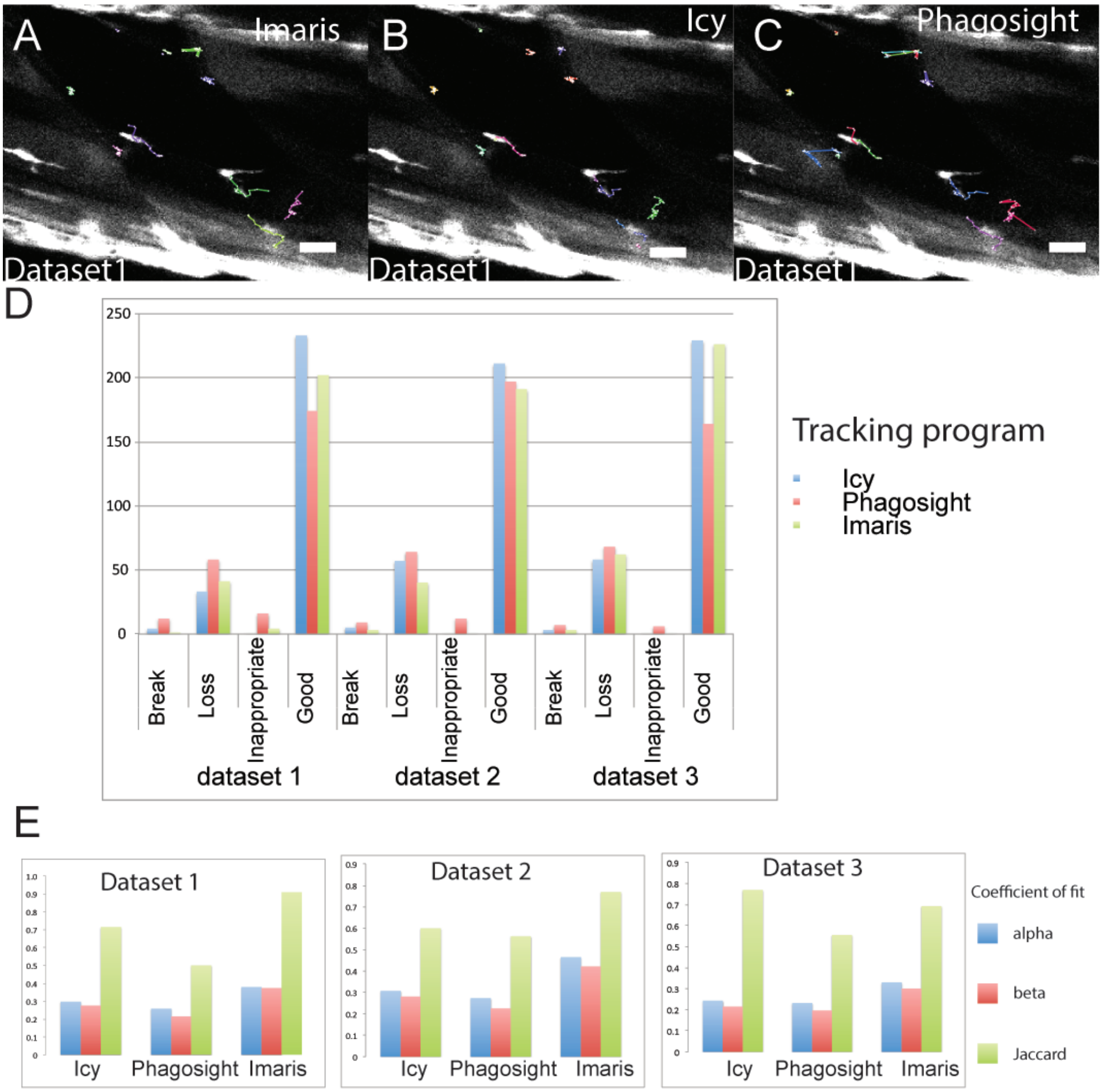

A
Imaris
Dataset1
B
Icy
Dataset1
C
Phagosight
Dataset1
D
250
200
150
100
50
0
Break
Loss
Inappropriate
Good
dataset 1
dataset 2
dataset 3
Tracking program
Icy
Phagosight
Imaris
E
Dataset 1
Dataset 2
Dataset 3
Icy
Phagosight
Imaris
Coefficient of fit
alpha
beta
Jaccard

**Figure S1**

| | Distance (µm) | Displacement (µm) | Directionality | Directional autocorrelation | Instantaneous speed (µm/hr) | Surface Area (µm^2) | Volume (µm^3) | Sphericity (%) | Convexity (%) | Roundness (%) |
|---|---|---|---|---|---|---|---|---|---|---|
| Kruskal-Wallis statistic | 2.41e-10 | 5.58e-12 | < 2.2e-16 | 1.52e-8 | 0.0043 | < 2.2e-16 | < 2.2e-16 | < 2.2e-16 | < 2.2e-16 | < 2.2e-16 |
| Control-ROCKout | 6.6e-10 | 0.87 | 3.9e-09 | 0.00065 | 0.30 | 2e-16 | 1.4e-12 | <2e-16 | <2e-16 | <2e-16 |
| Control-Y27632 | 7.4e-05 | 7.2e-11 | < 2e-16 | 0.00079 | 0.011 | 2e-16 | < 2e-16 | <2e-16 | <2e-16 | 0.0034 |
| ROCKout-Y27632 | 0.0019 | 3.1e-09 | 0.00083 | 1.9e-10 | 0.0012 | 1.3e-15 | 8.4e-08 | 0.026 | 0.021 | <2e-16 |

**Table S1**

| | Distance (µm) | Displacement (µm) | Directionality | Directional autocorrelation | Instantaneous speed (µm/hr) | Surface Area (µm^2) | Volume (µm^3) | Sphericity (%) | Convexity (%) | Roundness (%) |
|---|---|---|---|---|---|---|---|---|---|---|
| KW statistic | 1.85e-09 | 2.42e-10 | 0.0011 | 5.65e-05 | 1.62e-11 | 0.061 | 0.20 | 0.059 | 0.12 | 0.32 |
| 2-3 | 1.29e-06 | 3.18e-05 | 0.1505 | 8.4e-05 | 9.10e-07 | 0.0098 | 0.11 | 0.0038 | 0.018 | 0.097 |
| 1-3 | 0.17 | 0.0264 | 0.1159 | 0.0015 | 0.0531 | 0.092 | 0.41 | 0.023 | 0.072 | 0.35 |
| 1-2 | 2.13e-10 | 2.57e-11 | 0.0068 | 0.6305 | 7.97e-13 | 0.13 | 0.13 | 0.23 | 0.24 | 0.16 |
| 0-3 | 0.11 | 0.0171 | 0.0120 | 1.1e-05 | 0.16 | 0.0078 | 0.038 | 0.024 | 0.014 | 0.39 |
| 0-2 | 3.31e-05 | 0.0131 | 0.0913 | 0.5269 | 5.94e-06 | 0.50 | 0.29 | 0.20 | 0.48 | 0.036 |
| 0-1 | 0.0065 | 2.38e-06 | 0.0001 | 0.2026 | 0.0016 | 0.12 | 0.044 | 0.47 | 0.22 | 0.22 |

### Table S2

**Table S3**

Cross correlation values for parameters of shape and movement across classes.

| | | | | | | Significance |
|---|---|---|---|---|---|---|
| comparison | Parameter 1 | Parameter 2 | cor | p | class | weighting |
| 1 | Convexity_. | Ellipse_A | -0.1342009 | 0.0001043 | B_I | |
| | Convexity_. | Ellipse_A | -0.0771927 | 0.00331192 | B_NI | |
| | Convexity_. | Ellipse_A | -0.0981427 | 0.0003986 | NB_I | |
| | Convexity_. | Ellipse_A | -0.1321582 | 1.66E-07 | NB_NI | * |
| 2 | Convexity_. | Ellipse_B | -0.0482205 | 0.1649023 | B_I | |
| | Convexity_. | Ellipse_B | -0.0083273 | 0.75170806 | B_NI | |
| | Convexity_. | Ellipse_B | -0.1341958 | 1.22E-06 | NB_I | |
| | Convexity_. | Ellipse_B | -0.0445719 | 0.07871035 | NB_NI | NS |
| 3 | Convexity_. | Ellipse_C | -0.1062653 | 0.00215932 | B_I | |
| | Convexity_. | Ellipse_C | -0.170527 | 6.72E-11 | B_NI | |
| | Convexity_. | Ellipse_C | -0.1044182 | 0.00016414 | NB_I | |
| | Convexity_. | Ellipse_C | -0.2406585 | 0 | NB_NI | * |
| 4 | Convexity_. | Elongation | 0.10979097 | 0.00152557 | B_I | |
| | Convexity_. | Elongation | -0.0140268 | 0.59406681 | B_NI | |
| | Convexity_. | Elongation | -0.0145762 | 0.59981208 | NB_I | |
| | Convexity_. | Elongation | -0.0636733 | 0.01197064 | NB_NI | NS |
| 5 | Convexity_. | Flatness_Ratio | -0.1359129 | 8.48E-05 | B_I | |
| | Convexity_. | Flatness_Ratio | -0.0376543 | 0.15239483 | B_NI | |
| | Convexity_. | Flatness_Ratio | -0.2853036 | 0 | NB_I | |
| | Convexity_. | Flatness_Ratio | -0.1440131 | 1.14E-08 | NB_NI | * |
| 6 | Convexity_. | Max_Feret_Diameter | -0.4020119 | 0 | B_I | |
| | Convexity_. | Max_Feret_Diameter | -0.5946732 | 0 | B_NI | |
| | Convexity_. | Max_Feret_Diameter | -0.6353698 | 0 | NB_I | |
| | Convexity_. | Max_Feret_Diameter | -0.5921512 | 0 | NB_NI | * |
| 7 | Directionality | Convexity_. | -0.0343352 | 0.33118219 | B_I | |
| | Directionality | Convexity_. | -0.0443303 | 0.09779324 | B_NI | |
| | Directionality | Convexity_. | -0.0108995 | 0.70556758 | NB_I | |
| | Directionality | Convexity_. | 0.04030877 | 0.11790975 | NB_NI | NS |
| 8 | Directionality | Ellipse_A | -0.0233787 | 0.50826124 | B_I | |
| | Directionality | Ellipse_A | -0.0170323 | 0.52487026 | B_NI | |
| | Directionality | Ellipse_A | 0.01701106 | 0.55539757 | NB_I | |
| | Directionality | Ellipse_A | -0.0120249 | 0.64101225 | NB_NI | NS |
| 9 | Directionality | Ellipse_B | -0.0370326 | 0.29458072 | B_I | |
| | Directionality | Ellipse_B | -0.0177609 | 0.50729234 | B_NI | |
| | Directionality | Ellipse_B | 0.0274358 | 0.34151462 | NB_I | |
| | Directionality | Ellipse_B | 0.0179823 | 0.48560268 | NB_NI | NS |
| 10 | Directionality | Ellipse_C | -0.031027 | 0.37991019 | B_I | |
| | Directionality | Ellipse_C | 0.0086824 | 0.74584954 | B_NI | |
| | Directionality | Ellipse_C | 0.02394454 | 0.406482 | NB_I | |
| | Directionality | Ellipse_C | -0.0336671 | 0.19161724 | NB_NI | NS |
| 11 | Directionality | Elongation | -0.1297293 | 0.00022778 | B_I | |
| | Directionality | Elongation | 0.00485158 | 0.85628233 | B_NI | |
| | Directionality | Elongation | 0.03334789 | 0.2475797 | NB_I | |
| | Directionality | Elongation | -0.0370018 | 0.15121994 | NB_NI | NS |
| 12 | Directionality | Flatness_Ratio | 0.0038512 | 0.91323198 | B_I | |
| | Directionality | Flatness_Ratio | -0.0187837 | 0.48314728 | B_NI | |
| | Directionality | Flatness_Ratio | 0.01548588 | 0.59139619 | NB_I | |
| | Directionality | Flatness_Ratio | 0.031067 | 0.22823667 | NB_NI | NS |
| 13 | Directionality | Max_Feret_Diameter | -0.0406904 | 0.24942993 | B_I | |
| | Directionality | Max_Feret_Diameter | 0.14013021 | 1.46E-07 | B_NI | |
| | Directionality | Max_Feret_Diameter | 0.04903847 | 0.08897542 | NB_I | |
| | Directionality | Max_Feret_Diameter | -0.0162614 | 0.52831816 | NB_NI | NS |
| 14 | Directionality | Roundness_. | -0.0296138 | 0.40200257 | B_I | |
| | Directionality | Roundness_. | -0.0973193 | 0.00027101 | B_NI | |
| | Directionality | Roundness_. | -0.0733883 | 0.01085701 | NB_I | |

| | | | | | | |
|---|---|---|---|---|---|---|
| | Directionality | Roundness_. | 0.0020959 | 0.93522824 | NB_NI | NS |
| 15 | Directionality | Sphericity_. | -0.0376182 | 0.28700581 | B_I | |
| | Directionality | Sphericity_. | -0.0633881 | 0.01785381 | B_NI | |
| | Directionality | Sphericity_. | 0.07073178 | 0.01409525 | NB_I | |
| | Directionality | Sphericity_. | 0.04303997 | 0.09498898 | NB_NI | NS |
| 16 | Directionality | Surface_Area | 0.09968589 | 0.00469175 | B_I | |
| | Directionality | Surface_Area | 0.08173048 | 0.0022425 | B_NI | |
| | Directionality | Surface_Area | -0.0559867 | 0.05211638 | NB_I | |
| | Directionality | Surface_Area | -0.0808041 | 0.00169925 | NB_NI | * |
| 17 | Directionality | Volume | 0.14176981 | 5.54E-05 | B_I | |
| | Directionality | Volume | 0.10787848 | 5.37E-05 | B_NI | |
| | Directionality | Volume | -0.0029108 | 0.91963343 | NB_I | |
| | Directionality | Volume | -0.0896425 | 0.00049601 | NB_NI | * |
| 18 | Directionality | X_size | -0.0277842 | 0.4317186 | B_I | |
| | Directionality | X_size | 0.13454624 | 4.52E-07 | B_NI | |
| | Directionality | X_size | 0.04351831 | 0.13125397 | NB_I | |
| | Directionality | X_size | -0.0095507 | 0.71113295 | NB_NI | NS |
| 19 | Directionality | Y_size | -0.0021362 | 0.95180488 | B_I | |
| | Directionality | Y_size | 0.06467917 | 0.0156499 | B_NI | |
| | Directionality | Y_size | 0.03683987 | 0.20146053 | NB_I | |
| | Directionality | Y_size | -0.0272711 | 0.29022232 | NB_NI | NS |
| 20 | Directionality | Z_size | 0.1343978 | 0.00013352 | B_I | |
| | Directionality | Z_size | 0.05160434 | 0.05389812 | B_NI | |
| | Directionality | Z_size | -0.140616 | 9.66E-07 | NB_I | |
| | Directionality | Z_size | -0.0512889 | 0.04658682 | NB_NI | * |
| 21 | Ellipse_A | Ellipse_B | -0.0081213 | 0.81516826 | B_I | |
| | Ellipse_A | Ellipse_B | 0.00308907 | 0.90657105 | B_NI | |
| | Ellipse_A | Ellipse_B | 0.07908726 | 0.00435739 | NB_I | |
| | Ellipse_A | Ellipse_B | 0.01265245 | 0.61787102 | NB_NI | NS |
| 22 | Ellipse_A | Ellipse_C | 0.09189574 | 0.00803233 | B_I | |
| | Ellipse_A | Ellipse_C | 0.44095296 | 0 | B_NI | |
| | Ellipse_A | Ellipse_C | 0.15713197 | 1.26E-08 | NB_I | |
| | Ellipse_A | Ellipse_C | 0.24057916 | 0 | NB_NI | * |
| 23 | Ellipse_A | Elongation | 0.70130503 | 0 | B_I | |
| | Ellipse_A | Elongation | 0.87796605 | 0 | B_NI | |
| | Ellipse_A | Elongation | 0.50870252 | 0 | NB_I | |
| | Ellipse_A | Elongation | 0.901443 | 0 | NB_NI | * |
| 24 | Ellipse_A | Flatness_Ratio | -0.0087235 | 0.80173691 | B_I | |
| | Ellipse_A | Flatness_Ratio | -0.0019378 | 0.94131018 | B_NI | |
| | Ellipse_A | Flatness_Ratio | 0.01121174 | 0.68653824 | NB_I | |
| | Ellipse_A | Flatness_Ratio | 0.00343685 | 0.89221215 | NB_NI | NS |
| 25 | Ellipse_B | Ellipse_C | 0.32169312 | 0 | B_I | |
| | Ellipse_B | Ellipse_C | 0.19915795 | 2.13E-14 | B_NI | |
| | Ellipse_B | Ellipse_C | 0.53321636 | 0 | NB_I | |
| | Ellipse_B | Ellipse_C | 0.28872606 | 0 | NB_NI | * |
| 26 | Ellipse_B | Elongation | -0.0574443 | 0.09795976 | B_I | |
| | Ellipse_B | Elongation | -0.0100304 | 0.70312941 | B_NI | |
| | Ellipse_B | Elongation | -0.072559 | 0.00892101 | NB_I | |
| | Ellipse_B | Elongation | -0.0114179 | 0.65257318 | NB_NI | NS |
| 27 | Ellipse_B | Flatness_Ratio | 0.93762028 | 0 | B_I | |
| | Ellipse_B | Flatness_Ratio | 0.99648869 | 0 | B_NI | |
| | Ellipse_B | Flatness_Ratio | 0.44592172 | 0 | NB_I | |
| | Ellipse_B | Flatness_Ratio | 0.95222205 | 0 | NB_NI | * |
| 28 | Ellipse_C | Elongation | -0.1162253 | 0.00078857 | B_I | |
| | Ellipse_C | Elongation | 0.27555656 | 0 | B_NI | |
| | Ellipse_C | Elongation | -0.0483337 | 0.08173684 | NB_I | |
| | Ellipse_C | Elongation | 0.12035219 | 1.91E-06 | NB_NI | * |
| 29 | Ellipse_C | Flatness_Ratio | 0.20437445 | 2.76E-09 | B_I | |

| | | | | | | |
|---|---|---|---|---|---|---|
| | Ellipse_C | Flatness_Ratio | 0.173253 | 3.30E-11 | B_NI | |
| | Ellipse_C | Flatness_Ratio | -0.0189737 | 0.4946166 | NB_I | |
| | Ellipse_C | Flatness_Ratio | 0.18198234 | 4.64E-13 | NB_NI | * |
| 30 | Elongation | Flatness_Ratio | -0.1185759 | 0.00061448 | B_I | |
| | Elongation | Flatness_Ratio | -0.0199463 | 0.44850901 | B_NI | |
| | Elongation | Flatness_Ratio | -0.1631567 | 3.38E-09 | NB_I | |
| | Elongation | Flatness_Ratio | -0.0472429 | 0.06236496 | NB_NI | NS |
| 31 | Instantaneous_Speed | Convexity_. | -0.1171073 | 0.00088489 | B_I | |
| | Instantaneous_Speed | Convexity_. | -0.1979037 | 8.55E-14 | B_NI | |
| | Instantaneous_Speed | Convexity_. | -0.1187167 | 3.63E-05 | NB_I | |
| | Instantaneous_Speed | Convexity_. | -0.0986025 | 0.00012677 | NB_NI | * |
| 32 | Instantaneous_Speed | Directionality | 0.06242842 | 0.03114585 | B_I | |
| | Instantaneous_Speed | Directionality | 0.06400982 | 0.00788369 | B_NI | |
| | Instantaneous_Speed | Directionality | 0.05536192 | 0.02694151 | NB_I | |
| | Instantaneous_Speed | Directionality | -0.0133527 | 0.55576944 | NB_NI | NS |
| 33 | Instantaneous_Speed | Ellipse_A | 0.04056222 | 0.25092566 | B_I | |
| | Instantaneous_Speed | Ellipse_A | -0.0010171 | 0.9697133 | B_NI | |
| | Instantaneous_Speed | Ellipse_A | 0.02997916 | 0.29862288 | NB_I | |
| | Instantaneous_Speed | Ellipse_A | -0.0026178 | 0.91914887 | NB_NI | NS |
| 34 | Instantaneous_Speed | Ellipse_B | -0.0080517 | 0.81979273 | B_I | |
| | Instantaneous_Speed | Ellipse_B | -0.0007087 | 0.97889389 | B_NI | |
| | Instantaneous_Speed | Ellipse_B | 0.01492679 | 0.60485521 | NB_I | |
| | Instantaneous_Speed | Ellipse_B | -0.006902 | 0.78898591 | NB_NI | NS |
| 35 | Instantaneous_Speed | Ellipse_C | 0.01200098 | 0.73419001 | B_I | |
| | Instantaneous_Speed | Ellipse_C | 0.01238902 | 0.64372625 | B_NI | |
| | Instantaneous_Speed | Ellipse_C | -0.0546316 | 0.05808008 | NB_I | |
| | Instantaneous_Speed | Ellipse_C | 0.03408169 | 0.18620106 | NB_NI | NS |
| 36 | Instantaneous_Speed | Elongation | 0.02700499 | 0.44475129 | B_I | |
| | Instantaneous_Speed | Elongation | 0.00963921 | 0.71896965 | B_NI | |
| | Instantaneous_Speed | Elongation | 0.12571403 | 1.21E-05 | NB_I | |
| | Instantaneous_Speed | Elongation | 0.00714551 | 0.78172576 | NB_NI | NS |
| 37 | Instantaneous_Speed | Flatness_Ratio | 0.01026208 | 0.7715456 | B_I | |
| | Instantaneous_Speed | Flatness_Ratio | 0.0082319 | 0.75861581 | B_NI | |
| | Instantaneous_Speed | Flatness_Ratio | 0.03898017 | 0.17648085 | NB_I | |
| | Instantaneous_Speed | Flatness_Ratio | -0.0019845 | 0.93866516 | NB_NI | NS |
| 38 | Instantaneous_Speed | Max_Feret_Diameter | 0.12447563 | 0.00040691 | B_I | |
| | Instantaneous_Speed | Max_Feret_Diameter | 0.24145927 | 0 | B_NI | |
| | Instantaneous_Speed | Max_Feret_Diameter | 0.18091476 | 2.56E-10 | NB_I | |
| | Instantaneous_Speed | Max_Feret_Diameter | 0.11882038 | 3.77E-06 | NB_NI | * |
| 39 | Instantaneous_Speed | Roundness_. | -0.1214891 | 0.00056039 | B_I | |
| | Instantaneous_Speed | Roundness_. | -0.1549447 | 5.91E-09 | B_NI | |
| | Instantaneous_Speed | Roundness_. | -0.1731074 | 1.48E-09 | NB_I | |
| | Instantaneous_Speed | Roundness_. | -0.0661203 | 0.01026945 | NB_NI | * |
| 40 | Instantaneous_Speed | Sphericity_. | -0.0660722 | 0.0612849 | B_I | |
| | Instantaneous_Speed | Sphericity_. | -0.1614091 | 1.32E-09 | B_NI | |
| | Instantaneous_Speed | Sphericity_. | -0.114509 | 6.83E-05 | NB_I | |
| | Instantaneous_Speed | Sphericity_. | -0.0838177 | 0.00113089 | NB_NI | * |
| 41 | Instantaneous_Speed | Surface_Area | 0.03551306 | 0.31485252 | B_I | |
| | Instantaneous_Speed | Surface_Area | 0.20415029 | 1.33E-14 | B_NI | |
| | Instantaneous_Speed | Surface_Area | 0.0653233 | 0.02340851 | NB_I | |
| | Instantaneous_Speed | Surface_Area | 0.11194217 | 1.34E-05 | NB_NI | * |
| 42 | Instantaneous_Speed | Total_displacement | 0.31955203 | 0 | B_I | |
| | Instantaneous_Speed | Total_displacement | 0.23442675 | 0 | B_NI | |
| | Instantaneous_Speed | Total_displacement | 0.26047119 | 0 | NB_I | |
| | Instantaneous_Speed | Total_displacement | 0.22336558 | 0 | NB_NI | * |
| 43 | Instantaneous_Speed | Total_distance | 0.28181186 | 0 | B_I | |
| | Instantaneous_Speed | Total_distance | 0.21650349 | 0 | B_NI | |
| | Instantaneous_Speed | Total_distance | 0.33511588 | 0 | NB_I | |

| | | | | | | |
|---|---|---|---|---|---|---|
| | Instantaneous_Speed | Total_distance | 0.27951902 | 0 | NB_NI | * |
| 44 | Instantaneous_Speed | Volume | 0.0416734 | 0.23816631 | B_I | |
| | Instantaneous_Speed | Volume | 0.13898326 | 1.85E-07 | B_NI | |
| | Instantaneous_Speed | Volume | 0.03102822 | 0.28202581 | NB_I | |
| | Instantaneous_Speed | Volume | 0.10878737 | 2.33E-05 | NB_NI | * |
| 45 | Instantaneous_Speed | X_size | 0.13156758 | 0.00018496 | B_I | |
| | Instantaneous_Speed | X_size | 0.24000408 | 0 | B_NI | |
| | Instantaneous_Speed | X_size | 0.18785654 | 5.03E-11 | NB_I | |
| | Instantaneous_Speed | X_size | 0.1123905 | 1.23E-05 | NB_NI | * |
| 46 | Instantaneous_Speed | Y_size | 0.02314061 | 0.51259172 | B_I | |
| | Instantaneous_Speed | Y_size | 0.0819255 | 0.00218839 | B_NI | |
| | Instantaneous_Speed | Y_size | 0.02282655 | 0.4287503 | NB_I | |
| | Instantaneous_Speed | Y_size | 0.07348767 | 0.0043262 | NB_NI | * |
| 47 | Instantaneous_Speed | Z_size | 0.02568109 | 0.46739892 | B_I | |
| | Instantaneous_Speed | Z_size | 0.14881977 | 2.32E-08 | B_NI | |
| | Instantaneous_Speed | Z_size | 0.01122609 | 0.69717369 | NB_I | |
| | Instantaneous_Speed | Z_size | 0.04796381 | 0.06276289 | NB_NI | NS |
| 48 | Max_Feret_Diameter | Ellipse_A | 0.25375873 | 1.12E-13 | B_I | |
| | Max_Feret_Diameter | Ellipse_A | 0.05565497 | 0.03433096 | B_NI | |
| | Max_Feret_Diameter | Ellipse_A | 0.095915 | 0.00053967 | NB_I | |
| | Max_Feret_Diameter | Ellipse_A | 0.1302067 | 2.53E-07 | NB_NI | * |
| 49 | Max_Feret_Diameter | Ellipse_B | 0.09749666 | 0.00490791 | B_I | |
| | Max_Feret_Diameter | Ellipse_B | 0.01518833 | 0.5638786 | B_NI | |
| | Max_Feret_Diameter | Ellipse_B | 0.12351413 | 8.09E-06 | NB_I | |
| | Max_Feret_Diameter | Ellipse_B | 0.03786833 | 0.13528649 | NB_NI | NS |
| 50 | Max_Feret_Diameter | Ellipse_C | -0.0187733 | 0.58891323 | B_I | |
| | Max_Feret_Diameter | Ellipse_C | -0.0350365 | 0.18300273 | B_NI | |
| | Max_Feret_Diameter | Ellipse_C | -0.0216764 | 0.43521942 | NB_I | |
| | Max_Feret_Diameter | Ellipse_C | 0.03302832 | 0.19272229 | NB_NI | NS |
| 51 | Max_Feret_Diameter | Elongation | 0.31559172 | 0 | B_I | |
| | Max_Feret_Diameter | Elongation | 0.11530654 | 1.10E-05 | B_NI | |
| | Max_Feret_Diameter | Elongation | 0.22562689 | 2.22E-16 | NB_I | |
| | Max_Feret_Diameter | Elongation | 0.19426528 | 1.07E-14 | NB_NI | * |
| 52 | Max_Feret_Diameter | Flatness_Ratio | 0.15608375 | 6.17E-06 | B_I | |
| | Max_Feret_Diameter | Flatness_Ratio | 0.04416628 | 0.09318179 | B_NI | |
| | Max_Feret_Diameter | Flatness_Ratio | 0.28700814 | 0 | NB_I | |
| | Max_Feret_Diameter | Flatness_Ratio | 0.12913622 | 3.17E-07 | NB_NI | * |
| 53 | Roundness_. | Convexity_. | 0.57060558 | 0 | B_I | |
| | Roundness_. | Convexity_. | 0.66741312 | 0 | B_NI | |
| | Roundness_. | Convexity_. | 0.62570381 | 0 | NB_I | |
| | Roundness_. | Convexity_. | 0.53401101 | 0 | NB_NI | * |
| 54 | Roundness_. | Ellipse_A | -0.1865747 | 6.03E-08 | B_I | |
| | Roundness_. | Ellipse_A | -0.0932183 | 0.00038609 | B_NI | |
| | Roundness_. | Ellipse_A | -0.107154 | 0.00010976 | NB_I | |
| | Roundness_. | Ellipse_A | -0.1031006 | 4.59E-05 | NB_NI | * |
| 55 | Roundness_. | Ellipse_B | -0.0592236 | 0.08797622 | B_I | |
| | Roundness_. | Ellipse_B | -0.0093786 | 0.72159029 | B_NI | |
| | Roundness_. | Ellipse_B | -0.1014484 | 0.00025135 | NB_I | |
| | Roundness_. | Ellipse_B | -0.0412622 | 0.10362162 | NB_NI | NS |
| 56 | Roundness_. | Ellipse_C | 0.05374611 | 0.12159103 | B_I | |
| | Roundness_. | Ellipse_C | -0.0209381 | 0.42626561 | B_NI | |
| | Roundness_. | Ellipse_C | 0.03060014 | 0.27061363 | NB_I | |
| | Roundness_. | Ellipse_C | 0.01566536 | 0.53678727 | NB_NI | NS |
| 57 | Roundness_. | Elongation | -0.1916774 | 2.56E-08 | B_I | |
| | Roundness_. | Elongation | -0.1245853 | 2.01E-06 | B_NI | |
| | Roundness_. | Elongation | -0.1865087 | 1.27E-11 | NB_I | |
| | Roundness_. | Elongation | -0.1473594 | 5.16E-09 | NB_NI | * |
| 58 | Roundness_. | Flatness_Ratio | -0.1365048 | 7.89E-05 | B_I | |

| | | | | | | |
|---|---|---|---|---|---|---|
| | Roundness_. | Flatness_Ratio | -0.036335 | 0.16729545 | B_NI | |
| | Roundness_. | Flatness_Ratio | -0.2992585 | 0 | NB_I | |
| | Roundness_. | Flatness_Ratio | -0.12594 | 6.19E-07 | NB_NI | * |
| 59 | Roundness_. | Max_Feret_Diameter | -0.5789781 | 0 | B_I | |
| | Roundness_. | Max_Feret_Diameter | -0.6375035 | 0 | B_NI | |
| | Roundness_. | Max_Feret_Diameter | -0.7151635 | 0 | NB_I | |
| | Roundness_. | Max_Feret_Diameter | -0.650316 | 0 | NB_NI | * |
| 60 | Sphericity_. | Convexity_. | 0.7716229 | 0 | B_I | |
| | Sphericity_. | Convexity_. | 0.84727472 | 0 | B_NI | |
| | Sphericity_. | Convexity_. | 0.7729547 | 0 | NB_I | |
| | Sphericity_. | Convexity_. | 0.68545753 | 0 | NB_NI | * |
| 61 | Sphericity_. | Ellipse_A | -0.1327492 | 0.00012404 | B_I | |
| | Sphericity_. | Ellipse_A | -0.0476482 | 0.07008692 | B_NI | |
| | Sphericity_. | Ellipse_A | -0.1024595 | 0.00021768 | NB_I | |
| | Sphericity_. | Ellipse_A | -0.1018641 | 5.66E-05 | NB_NI | * |
| 62 | Sphericity_. | Ellipse_B | -0.020235 | 0.56023228 | B_I | |
| | Sphericity_. | Ellipse_B | -0.003432 | 0.89625495 | B_NI | |
| | Sphericity_. | Ellipse_B | -0.1207995 | 1.28E-05 | NB_I | |
| | Sphericity_. | Ellipse_B | -0.0283073 | 0.26429432 | NB_NI | NS |
| 63 | Sphericity_. | Ellipse_C | 0.02390745 | 0.49130003 | B_I | |
| | Sphericity_. | Ellipse_C | -0.1126811 | 1.75E-05 | B_NI | |
| | Sphericity_. | Ellipse_C | 0.00472396 | 0.86498599 | NB_I | |
| | Sphericity_. | Ellipse_C | -0.0517177 | 0.04130565 | NB_NI | NS |
| 64 | Sphericity_. | Elongation | 0.00329699 | 0.92439444 | B_I | |
| | Sphericity_. | Elongation | -0.0099765 | 0.70464936 | B_NI | |
| | Sphericity_. | Elongation | -0.0819973 | 0.00311342 | NB_I | |
| | Sphericity_. | Elongation | -0.0918489 | 0.00028451 | NB_NI | * |
| 65 | Sphericity_. | Flatness_Ratio | -0.0936642 | 0.00689356 | B_I | |
| | Sphericity_. | Flatness_Ratio | -0.0337747 | 0.19928749 | B_NI | |
| | Sphericity_. | Flatness_Ratio | -0.3471782 | 0 | NB_I | |
| | Sphericity_. | Flatness_Ratio | -0.1379655 | 4.61E-08 | NB_NI | * |
| 66 | Sphericity_. | Max_Feret_Diameter | -0.4136333 | 0 | B_I | |
| | Sphericity_. | Max_Feret_Diameter | -0.6126686 | 0 | B_NI | |
| | Sphericity_. | Max_Feret_Diameter | -0.6561826 | 0 | NB_I | |
| | Sphericity_. | Max_Feret_Diameter | -0.6742815 | 0 | NB_NI | * |
| 67 | Sphericity_. | Roundness_. | 0.64870644 | 0 | B_I | |
| | Sphericity_. | Roundness_. | 0.72973853 | 0 | B_NI | |
| | Sphericity_. | Roundness_. | 0.66522938 | 0 | NB_I | |
| | Sphericity_. | Roundness_. | 0.72048014 | 0 | NB_NI | * |
| 68 | Surface_Area | Convexity_. | -0.679974 | 0 | B_I | |
| | Surface_Area | Convexity_. | -0.7836588 | 0 | B_NI | |
| | Surface_Area | Convexity_. | -0.7530592 | 0 | NB_I | |
| | Surface_Area | Convexity_. | -0.7518147 | 0 | NB_NI | * |
| 69 | Surface_Area | Ellipse_A | 0.15604641 | 6.20E-06 | B_I | |
| | Surface_Area | Ellipse_A | 0.04236478 | 0.10733109 | B_NI | |
| | Surface_Area | Ellipse_A | 0.08910769 | 0.00131058 | NB_I | |
| | Surface_Area | Ellipse_A | 0.09027191 | 0.00036185 | NB_NI | * |
| 70 | Surface_Area | Ellipse_B | 0.01339054 | 0.6999069 | B_I | |
| | Surface_Area | Ellipse_B | 0.00317531 | 0.90397521 | B_NI | |
| | Surface_Area | Ellipse_B | 0.13050997 | 2.39E-06 | NB_I | |
| | Surface_Area | Ellipse_B | 0.03592582 | 0.15650927 | NB_NI | NS |
| 71 | Surface_Area | Ellipse_C | 0.04938 | 0.1549665 | B_I | |
| | Surface_Area | Ellipse_C | 0.19372568 | 1.08E-13 | B_NI | |
| | Surface_Area | Ellipse_C | 0.06333176 | 0.02250148 | NB_I | |
| | Surface_Area | Ellipse_C | 0.16860437 | 2.15E-11 | NB_NI | * |
| 72 | Surface_Area | Elongation | -0.0673141 | 0.05241035 | B_I | |
| | Surface_Area | Elongation | -0.0150166 | 0.56829396 | B_NI | |
| | Surface_Area | Elongation | -0.0101548 | 0.7147302 | NB_I | |

| | | | | | | |
|---|---|---|---|---|---|---|
| | Surface_Area | Elongation | 0.02811559 | 0.26754375 | NB_NI | NS |
| 73 | Surface_Area | Flatness_Ratio | 0.08061173 | 0.02012093 | B_I | |
| | Surface_Area | Flatness_Ratio | 0.02695807 | 0.30563916 | B_NI | |
| | Surface_Area | Flatness_Ratio | 0.33756283 | 0 | NB_I | |
| | Surface_Area | Flatness_Ratio | 0.12359816 | 9.99E-07 | NB_NI | * |
| 74 | Surface_Area | Max_Feret_Diameter | 0.52018273 | 0 | B_I | |
| | Surface_Area | Max_Feret_Diameter | 0.67970234 | 0 | B_NI | |
| | Surface_Area | Max_Feret_Diameter | 0.68499994 | 0 | NB_I | |
| | Surface_Area | Max_Feret_Diameter | 0.74007779 | 0 | NB_NI | * |
| 75 | Surface_Area | Roundness_. | -0.4671117 | 0 | B_I | |
| | Surface_Area | Roundness_. | -0.5517236 | 0 | B_NI | |
| | Surface_Area | Roundness_. | -0.5386282 | 0 | NB_I | |
| | Surface_Area | Roundness_. | -0.5328427 | 0 | NB_NI | * |
| 76 | Surface_Area | Sphericity_. | -0.777285 | 0 | B_I | |
| | Surface_Area | Sphericity_. | -0.8453241 | 0 | B_NI | |
| | Surface_Area | Sphericity_. | -0.823664 | 0 | NB_I | |
| | Surface_Area | Sphericity_. | -0.7931702 | 0 | NB_NI | * |
| 77 | Surface_Area | Volume | 0.7590819 | 0 | B_I | |
| | Surface_Area | Volume | 0.8433035 | 0 | B_NI | |
| | Surface_Area | Volume | 0.85335535 | 0 | NB_I | |
| | Surface_Area | Volume | 0.89580256 | 0 | NB_NI | * |
| 78 | Total_displacement | Convexity_. | -0.0779463 | 0.02719522 | B_I | |
| | Total_displacement | Convexity_. | -0.0514118 | 0.05480088 | B_NI | |
| | Total_displacement | Convexity_. | -0.1590166 | 2.90E-08 | NB_I | |
| | Total_displacement | Convexity_. | 0.04327767 | 0.09317581 | NB_NI | NS |
| 79 | Total_displacement | Directionality | 0.21773058 | 2.93E-14 | B_I | |
| | Total_displacement | Directionality | 0.18973792 | 2.00E-15 | B_NI | |
| | Total_displacement | Directionality | 0.25922379 | 0 | NB_I | |
| | Total_displacement | Directionality | 0.10096861 | 7.97E-06 | NB_NI | * |
| 80 | Total_displacement | Ellipse_A | -0.0575848 | 0.10297351 | B_I | |
| | Total_displacement | Ellipse_A | -0.0042724 | 0.87328544 | B_NI | |
| | Total_displacement | Ellipse_A | 0.01776043 | 0.53811077 | NB_I | |
| | Total_displacement | Ellipse_A | -0.0261617 | 0.31030058 | NB_NI | NS |
| 81 | Total_displacement | Ellipse_B | -0.0119404 | 0.73548189 | B_I | |
| | Total_displacement | Ellipse_B | -0.0207344 | 0.43887765 | B_NI | |
| | Total_displacement | Ellipse_B | 0.00979237 | 0.73428012 | NB_I | |
| | Total_displacement | Ellipse_B | -0.0301782 | 0.24182891 | NB_NI | NS |
| 82 | Total_displacement | Ellipse_C | -0.0277892 | 0.43163589 | B_I | |
| | Total_displacement | Ellipse_C | 0.01994998 | 0.45639309 | B_NI | |
| | Total_displacement | Ellipse_C | -0.0194272 | 0.50065298 | NB_I | |
| | Total_displacement | Ellipse_C | -0.062091 | 0.01595668 | NB_NI | * |
| 83 | Total_displacement | Elongation | -0.1472546 | 2.80E-05 | B_I | |
| | Total_displacement | Elongation | 0.041201 | 0.12388473 | B_NI | |
| | Total_displacement | Elongation | 0.07557487 | 0.00870624 | NB_I | |
| | Total_displacement | Elongation | 0.03666059 | 0.15502855 | NB_NI | NS |
| 84 | Total_displacement | Flatness_Ratio | 0.02159754 | 0.54111168 | B_I | |
| | Total_displacement | Flatness_Ratio | -0.0245889 | 0.35860082 | B_NI | |
| | Total_displacement | Flatness_Ratio | 0.04956569 | 0.08558973 | NB_I | |
| | Total_displacement | Flatness_Ratio | -0.0286974 | 0.2657213 | NB_NI | NS |
| 85 | Total_displacement | Max_Feret_Diameter | -0.1732762 | 7.81E-07 | B_I | |
| | Total_displacement | Max_Feret_Diameter | 0.24498929 | 0 | B_NI | |
| | Total_displacement | Max_Feret_Diameter | 0.15587388 | 5.44E-08 | NB_I | |
| | Total_displacement | Max_Feret_Diameter | 0.05226362 | 0.04256967 | NB_NI | * |
| 86 | Total_displacement | Roundness_. | 0.01409196 | 0.69009576 | B_I | |
| | Total_displacement | Roundness_. | -0.1372616 | 2.63E-07 | B_NI | |
| | Total_displacement | Roundness_. | -0.1895765 | 3.32E-11 | NB_I | |
| | Total_displacement | Roundness_. | -0.0313791 | 0.22359489 | NB_NI | NS |
| 87 | Total_displacement | Sphericity_. | -0.0358027 | 0.3109196 | B_I | |

| | | | | | | |
|---|---|---|---|---|---|---|
| | Total_displacement | Sphericity_. | -0.1171309 | 1.15E-05 | B_NI | |
| | Total_displacement | Sphericity_. | -0.0913832 | 0.00150235 | NB_I | |
| | Total_displacement | Sphericity_. | -0.0813258 | 0.00158507 | NB_NI | * |
| 88 | Total_displacement | Surface_Area | -0.0179103 | 0.61231032 | B_I | |
| | Total_displacement | Surface_Area | 0.17789288 | 2.17E-11 | B_NI | |
| | Total_displacement | Surface_Area | 0.03526543 | 0.22141501 | NB_I | |
| | Total_displacement | Surface_Area | 0.02297227 | 0.37300054 | NB_NI | NS |
| 89 | Total_displacement | Total_distance | 0.61494023 | 0 | B_I | |
| | Total_displacement | Total_distance | 0.57750356 | 0 | B_NI | |
| | Total_displacement | Total_distance | 0.64122641 | 0 | NB_I | |
| | Total_displacement | Total_distance | 0.50540245 | 0 | NB_NI | * |
| 90 | Total_displacement | Volume | -0.0281307 | 0.42599627 | B_I | |
| | Total_displacement | Volume | 0.18230699 | 6.74E-12 | B_NI | |
| | Total_displacement | Volume | 0.01747338 | 0.54470074 | NB_I | |
| | Total_displacement | Volume | -0.0337971 | 0.18990605 | NB_NI | NS |
| 91 | Total_displacement | X_size | -0.1927982 | 3.66E-08 | B_I | |
| | Total_displacement | X_size | 0.24584679 | 0 | B_NI | |
| | Total_displacement | X_size | 0.14791402 | 2.53E-07 | NB_I | |
| | Total_displacement | X_size | 0.04367143 | 0.09023341 | NB_NI | NS |
| 92 | Total_displacement | Y_size | 0.02839779 | 0.42161416 | B_I | |
| | Total_displacement | Y_size | 0.06790353 | 0.01115702 | B_NI | |
| | Total_displacement | Y_size | 0.07471308 | 0.00950378 | NB_I | |
| | Total_displacement | Y_size | 0.00610689 | 0.81281443 | NB_NI | NS |
| 93 | Total_displacement | Z_size | 0.10011142 | 0.00451701 | B_I | |
| | Total_displacement | Z_size | 0.13354054 | 5.52E-07 | B_NI | |
| | Total_displacement | Z_size | -0.0161326 | 0.57600095 | NB_I | |
| | Total_displacement | Z_size | -0.0326828 | 0.20493617 | NB_NI | NS |
| 94 | Total_distance | Convexity_. | -0.0254758 | 0.47096766 | B_I | |
| | Total_distance | Convexity_. | -0.1603265 | 1.70E-09 | B_NI | |
| | Total_distance | Convexity_. | -0.139061 | 1.27E-06 | NB_I | |
| | Total_distance | Convexity_. | -0.0461281 | 0.07352332 | NB_NI | NS |
| 95 | Total_distance | Directionality | -0.3796833 | 0 | B_I | |
| | Total_distance | Directionality | -0.3663 | 0 | B_NI | |
| | Total_distance | Directionality | -0.2940732 | 0 | NB_I | |
| | Total_distance | Directionality | -0.481795 | 0 | NB_NI | * |
| 96 | Total_distance | Ellipse_A | -0.0302802 | 0.39149008 | B_I | |
| | Total_distance | Ellipse_A | 0.00287256 | 0.91460525 | B_NI | |
| | Total_distance | Ellipse_A | 0.01351303 | 0.63948504 | NB_I | |
| | Total_distance | Ellipse_A | -0.008606 | 0.73860409 | NB_NI | NS |
| 97 | Total_distance | Ellipse_B | 0.05031779 | 0.15428749 | B_I | |
| | Total_distance | Ellipse_B | -0.0212183 | 0.42826825 | B_NI | |
| | Total_distance | Ellipse_B | -0.0104349 | 0.7175686 | NB_I | |
| | Total_distance | Ellipse_B | -0.032192 | 0.21182355 | NB_NI | NS |
| 98 | Total_distance | Ellipse_C | 0.00915713 | 0.79556656 | B_I | |
| | Total_distance | Ellipse_C | 0.01931446 | 0.4708673 | B_NI | |
| | Total_distance | Ellipse_C | -0.0364883 | 0.20579858 | NB_I | |
| | Total_distance | Ellipse_C | 0.0276357 | 0.28381943 | NB_NI | NS |
| 99 | Total_distance | Elongation | 0.00736406 | 0.83495176 | B_I | |
| | Total_distance | Elongation | 0.01305722 | 0.62594533 | B_NI | |
| | Total_distance | Elongation | 0.11781207 | 4.17E-05 | NB_I | |
| | Total_distance | Elongation | 0.06565519 | 0.01081782 | NB_NI | * |
| 100 | Total_distance | Flatness_Ratio | 0.04180372 | 0.23670037 | B_I | |
| | Total_distance | Flatness_Ratio | -0.0169018 | 0.52805134 | B_NI | |
| | Total_distance | Flatness_Ratio | 0.01406754 | 0.62580299 | NB_I | |
| | Total_distance | Flatness_Ratio | -0.0479242 | 0.06298032 | NB_NI | NS |
| 101 | Total_distance | Max_Feret_Diameter | -0.1215815 | 0.00055493 | B_I | |
| | Total_distance | Max_Feret_Diameter | 0.18301138 | 5.58E-12 | B_NI | |
| | Total_distance | Max_Feret_Diameter | 0.22036779 | 1.04E-14 | NB_I | |

| | | | | | | |
|---|---|---|---|---|---|---|
| | Total_distance | Max_Feret_Diameter | 0.08423042 | 0.00106846 | NB_NI | * |
| 102 | Total_distance | Roundness_. | 0.05799698 | 0.10052833 | B_I | |
| | Total_distance | Roundness_. | -0.0737547 | 0.00583358 | B_NI | |
| | Total_distance | Roundness_. | -0.1913467 | 2.16E-11 | NB_I | |
| | Total_distance | Roundness_. | -0.037371 | 0.14717915 | NB_NI | NS |
| 103 | Total_distance | Sphericity_. | 0.04881756 | 0.16696411 | B_I | |
| | Total_distance | Sphericity_. | -0.1587417 | 2.47E-09 | B_NI | |
| | Total_distance | Sphericity_. | -0.1627079 | 1.36E-08 | NB_I | |
| | Total_distance | Sphericity_. | -0.124466 | 1.27E-06 | NB_NI | * |
| 104 | Total_distance | Surface_Area | -0.145172 | 3.64E-05 | B_I | |
| | Total_distance | Surface_Area | 0.20524755 | 9.55E-15 | B_NI | |
| | Total_distance | Surface_Area | 0.07831734 | 0.00655065 | NB_I | |
| | Total_distance | Surface_Area | 0.13605763 | 1.16E-07 | NB_NI | * |
| 105 | Total_distance | Volume | -0.1652571 | 2.50E-06 | B_I | |
| | Total_distance | Volume | 0.12121696 | 5.58E-06 | B_NI | |
| | Total_distance | Volume | 0.0063814 | 0.82493793 | NB_I | |
| | Total_distance | Volume | 0.11180627 | 1.37E-05 | NB_NI | * |
| 106 | Total_distance | X_size | -0.1311588 | 0.00019377 | B_I | |
| | Total_distance | X_size | 0.18639854 | 2.22E-12 | B_NI | |
| | Total_distance | X_size | 0.22064883 | 9.77E-15 | NB_I | |
| | Total_distance | X_size | 0.06521327 | 0.0113628 | NB_NI | * |
| 107 | Total_distance | Y_size | 0.01422558 | 0.68731137 | B_I | |
| | Total_distance | Y_size | 0.03197831 | 0.23246097 | B_NI | |
| | Total_distance | Y_size | 0.052989 | 0.0660578 | NB_I | |
| | Total_distance | Y_size | 0.07147203 | 0.00552227 | NB_NI | * |
| 107 | Total_distance | Z_size | -0.0798675 | 0.02361615 | B_I | |
| | Total_distance | Z_size | 0.17265293 | 8.36E-11 | B_NI | |
| | Total_distance | Z_size | 0.04435681 | 0.12397961 | NB_I | |
| | Total_distance | Z_size | 0.07377354 | 0.00417705 | NB_NI | * |
| 108 | Volume | Convexity_. | -0.3963951 | 0 | B_I | |
| | Volume | Convexity_. | -0.6717472 | 0 | B_NI | |
| | Volume | Convexity_. | -0.6076085 | 0 | NB_I | |
| | Volume | Convexity_. | -0.65258 | 0 | NB_NI | * |
| 109 | Volume | Ellipse_A | 0.15105042 | 1.23E-05 | B_I | |
| | Volume | Ellipse_A | 0.0507524 | 0.05366871 | B_NI | |
| | Volume | Ellipse_A | 0.07057302 | 0.01098076 | NB_I | |
| | Volume | Ellipse_A | 0.07062133 | 0.00530544 | NB_NI | * |
| 110 | Volume | Ellipse_B | 0.01651026 | 0.63460281 | B_I | |
| | Volume | Ellipse_B | 0.00268462 | 0.91875819 | B_NI | |
| | Volume | Ellipse_B | 0.12622899 | 5.08E-06 | NB_I | |
| | Volume | Ellipse_B | 0.04991084 | 0.04894514 | NB_NI | * |
| 111 | Volume | Ellipse_C | 0.05942061 | 0.08692313 | B_I | |
| | Volume | Ellipse_C | 0.22103509 | 0 | B_NI | |
| | Volume | Ellipse_C | 0.13499029 | 1.06E-06 | NB_I | |
| | Volume | Ellipse_C | 0.21643096 | 0 | NB_NI | * |
| 112 | Volume | Elongation | -0.055895 | 0.10737083 | B_I | |
| | Volume | Elongation | 0.00935244 | 0.72233572 | B_NI | |
| | Volume | Elongation | -0.0550845 | 0.04723746 | NB_I | |
| | Volume | Elongation | -0.0095717 | 0.70588016 | NB_NI | NS |
| 113 | Volume | Flatness_Ratio | 0.08040741 | 0.02043979 | B_I | |
| | Volume | Flatness_Ratio | 0.01436425 | 0.58522045 | B_NI | |
| | Volume | Flatness_Ratio | 0.23797901 | 0 | NB_I | |
| | Volume | Flatness_Ratio | 0.09820076 | 0.00010396 | NB_NI | * |
| 114 | Volume | Max_Feret_Diameter | 0.5585984 | 0 | B_I | |
| | Volume | Max_Feret_Diameter | 0.64901799 | 0 | B_NI | |
| | Volume | Max_Feret_Diameter | 0.58631051 | 0 | NB_I | |
| | Volume | Max_Feret_Diameter | 0.61614788 | 0 | NB_NI | * |
| 115 | Volume | Roundness_. | -0.2391934 | 2.82E-12 | B_I | |

| | | | | | | |
|---|---|---|---|---|---|---|
| | Volume | Roundness_. | -0.4536803 | 0 | B_NI | |
| | Volume | Roundness_. | -0.3938958 | 0 | NB_I | |
| | Volume | Roundness_. | -0.3560166 | 0 | NB_NI | * |
| 116 | Volume | Sphericity_. | -0.3001768 | 0 | B_I | |
| | Volume | Sphericity_. | -0.6149626 | 0 | B_NI | |
| | Volume | Sphericity_. | -0.510987 | 0 | NB_I | |
| | Volume | Sphericity_. | -0.5172471 | 0 | NB_NI | * |
| 117 | X_displacement | Convexity_. | -0.0837574 | 0.01759989 | B_I | |
| | X_displacement | Convexity_. | -0.236919 | 0 | B_NI | |
| | X_displacement | Convexity_. | -0.1003386 | 0.00048883 | NB_I | |
| | X_displacement | Convexity_. | -0.0936075 | 0.00027517 | NB_NI | * |
| 118 | X_displacement | Directionality | -0.0261706 | 0.36665617 | B_I | |
| | X_displacement | Directionality | 0.02572283 | 0.28605251 | B_NI | |
| | X_displacement | Directionality | 0.03516287 | 0.16016124 | NB_I | |
| | X_displacement | Directionality | -0.0163508 | 0.47064427 | NB_NI | NS |
| 119 | X_displacement | Ellipse_A | 0.06786104 | 0.05457858 | B_I | |
| | X_displacement | Ellipse_A | 0.00104629 | 0.96884445 | B_NI | |
| | X_displacement | Ellipse_A | 0.01678389 | 0.56069113 | NB_I | |
| | X_displacement | Ellipse_A | 0.00736678 | 0.77514561 | NB_NI | NS |
| 120 | X_displacement | Ellipse_B | 0.00312547 | 0.92953525 | B_I | |
| | X_displacement | Ellipse_B | 0.00345122 | 0.89748964 | B_NI | |
| | X_displacement | Ellipse_B | -0.0026917 | 0.92566278 | NB_I | |
| | X_displacement | Ellipse_B | -0.0046457 | 0.85704541 | NB_NI | NS |
| 121 | X_displacement | Ellipse_C | 0.03303025 | 0.34989978 | B_I | |
| | X_displacement | Ellipse_C | -0.0003776 | 0.98875246 | B_NI | |
| | X_displacement | Ellipse_C | -0.0802092 | 0.00535689 | NB_I | |
| | X_displacement | Ellipse_C | 0.024284 | 0.34632021 | NB_NI | NS |
| 122 | X_displacement | Elongation | 0.0945793 | 0.00731929 | B_I | |
| | X_displacement | Elongation | 0.01858705 | 0.48774025 | B_NI | |
| | X_displacement | Elongation | 0.14753059 | 2.72E-07 | NB_I | |
| | X_displacement | Elongation | 0.02243534 | 0.38427863 | NB_NI | NS |
| 123 | X_displacement | Flatness_Ratio | 0.01219411 | 0.73007772 | B_I | |
| | X_displacement | Flatness_Ratio | 0.01616051 | 0.5463057 | B_NI | |
| | X_displacement | Flatness_Ratio | 0.021316 | 0.45993544 | NB_I | |
| | X_displacement | Flatness_Ratio | 0.00419098 | 0.87090733 | NB_NI | NS |
| 124 | X_displacement | Instantaneous_Speed | 0.72540057 | 0 | B_I | |
| | X_displacement | Instantaneous_Speed | 0.78461832 | 0 | B_NI | |
| | X_displacement | Instantaneous_Speed | 0.95762151 | 0 | NB_I | |
| | X_displacement | Instantaneous_Speed | 0.90243965 | 0 | NB_NI | * |
| 125 | X_displacement | Max_Feret_Diameter | 0.18432504 | 1.44E-07 | B_I | |
| | X_displacement | Max_Feret_Diameter | 0.3253879 | 0 | B_NI | |
| | X_displacement | Max_Feret_Diameter | 0.19727159 | 4.99E-12 | NB_I | |
| | X_displacement | Max_Feret_Diameter | 0.13599288 | 1.17E-07 | NB_NI | * |
| 126 | X_displacement | Roundness_. | -0.1324708 | 0.00016681 | B_I | |
| | X_displacement | Roundness_. | -0.1736825 | 6.44E-11 | B_NI | |
| | X_displacement | Roundness_. | -0.1720588 | 1.87E-09 | NB_I | |
| | X_displacement | Roundness_. | -0.0870849 | 0.00071661 | NB_NI | * |
| 127 | X_displacement | Sphericity_. | -0.0675817 | 0.05558369 | B_I | |
| | X_displacement | Sphericity_. | -0.2008429 | 3.60E-14 | B_NI | |
| | X_displacement | Sphericity_. | -0.1262657 | 1.11E-05 | NB_I | |
| | X_displacement | Sphericity_. | -0.0977061 | 0.00014608 | NB_NI | * |
| 128 | X_displacement | Surface_Area | 0.04204888 | 0.23395995 | B_I | |
| | X_displacement | Surface_Area | 0.25398767 | 0 | B_NI | |
| | X_displacement | Surface_Area | 0.06982786 | 0.01537692 | NB_I | |
| | X_displacement | Surface_Area | 0.10633887 | 3.55E-05 | NB_NI | * |
| 129 | X_displacement | Total_displacement | 0.0835858 | 0.00387877 | B_I | |
| | X_displacement | Total_displacement | 0.15292509 | 1.78E-10 | B_NI | |
| | X_displacement | Total_displacement | 0.24583839 | 0 | NB_I | |

| | | | | | | |
|---|---|---|---|---|---|---|
| | X_displacement | Total_displacement | 0.20037277 | 0 | NB_NI | * |
| 130 | X_displacement | Total_distance | 0.20914097 | 3.01E-13 | B_I | |
| | X_displacement | Total_distance | 0.21328136 | 0 | B_NI | |
| | X_displacement | Total_distance | 0.30722967 | 0 | NB_I | |
| | X_displacement | Total_distance | 0.24764036 | 0 | NB_NI | * |
| 131 | X_displacement | Volume | 0.05022683 | 0.15503471 | B_I | |
| | X_displacement | Volume | 0.17506562 | 4.52E-11 | B_NI | |
| | X_displacement | Volume | 0.02408772 | 0.40368076 | NB_I | |
| | X_displacement | Volume | 0.09158213 | 0.00037287 | NB_NI | * |
| 132 | X_displacement | X_size | 0.19200042 | 4.17E-08 | B_I | |
| | X_displacement | X_size | 0.32821482 | 0 | B_NI | |
| | X_displacement | X_size | 0.21272305 | 8.70E-14 | NB_I | |
| | X_displacement | X_size | 0.13720129 | 9.03E-08 | NB_NI | * |
| 133 | X_displacement | Y_displacement | 0.3229531 | 0 | B_I | |
| | X_displacement | Y_displacement | 0.30260524 | 0 | B_NI | |
| | X_displacement | Y_displacement | 0.49484107 | 0 | NB_I | |
| | X_displacement | Y_displacement | 0.34156653 | 0 | NB_NI | * |
| 134 | X_displacement | Y_size | -0.0076642 | 0.82832839 | B_I | |
| | X_displacement | Y_size | 0.076633 | 0.00417149 | B_NI | |
| | X_displacement | Y_size | -0.0128318 | 0.65646117 | NB_I | |
| | X_displacement | Y_size | 0.03488667 | 0.17601019 | NB_NI | NS |
| 135 | X_displacement | Z_displacement | 0.16088536 | 2.33E-08 | B_I | |
| | X_displacement | Z_displacement | 0.20694508 | 0 | B_NI | |
| | X_displacement | Z_displacement | 0.34467441 | 0 | NB_I | |
| | X_displacement | Z_displacement | 0.25308207 | 0 | NB_NI | * |
| 136 | X_displacement | Z_size | 0.02145085 | 0.54386334 | B_I | |
| | X_displacement | Z_size | 0.16507801 | 5.47E-10 | B_NI | |
| | X_displacement | Z_size | -0.0209539 | 0.4675947 | NB_I | |
| | X_displacement | Z_size | 0.02483697 | 0.33544627 | NB_NI | NS |
| 137 | X_position | Convexity_. | 0.0017165 | 0.96059473 | B_I | |
| | X_position | Convexity_. | 0.24172266 | 0 | B_NI | |
| | X_position | Convexity_. | -0.0389264 | 0.1610315 | NB_I | |
| | X_position | Convexity_. | 0.14899302 | 3.48E-09 | NB_NI | * |
| 138 | X_position | Directionality | 0.01948434 | 0.50316748 | B_I | |
| | X_position | Directionality | -0.1212391 | 4.87E-07 | B_NI | |
| | X_position | Directionality | 0.12803075 | 4.33E-07 | NB_I | |
| | X_position | Directionality | -0.0147541 | 0.51820979 | NB_NI | NS |
| 139 | X_position | Ellipse_A | 0.07549246 | 0.02955104 | B_I | |
| | X_position | Ellipse_A | -0.0126664 | 0.63033332 | B_NI | |
| | X_position | Ellipse_A | -0.0036324 | 0.89597871 | NB_I | |
| | X_position | Ellipse_A | -0.0718761 | 0.00454626 | NB_NI | * |
| 140 | X_position | Ellipse_B | -0.0305823 | 0.3786017 | B_I | |
| | X_position | Ellipse_B | 0.00735419 | 0.77992668 | B_NI | |
| | X_position | Ellipse_B | 0.01394243 | 0.61576951 | NB_I | |
| | X_position | Ellipse_B | 0.0225563 | 0.37376461 | NB_NI | NS |
| 141 | X_position | Ellipse_C | -0.0331577 | 0.33974763 | B_I | |
| | X_position | Ellipse_C | 0.01884022 | 0.47407285 | B_NI | |
| | X_position | Ellipse_C | -0.0192888 | 0.48747631 | NB_I | |
| | X_position | Ellipse_C | -0.0159599 | 0.52915765 | NB_NI | NS |
| 142 | X_position | Elongation | -0.0205241 | 0.55463965 | B_I | |
| | X_position | Elongation | 0.00185482 | 0.94381862 | B_NI | |
| | X_position | Elongation | -0.0717503 | 0.00971407 | NB_I | |
| | X_position | Elongation | -0.0651978 | 0.01007329 | NB_NI | * |
| 143 | X_position | Flatness_Ratio | 0.00114909 | 0.97361472 | B_I | |
| | X_position | Flatness_Ratio | -0.002263 | 0.93148362 | B_NI | |
| | X_position | Flatness_Ratio | 0.14911965 | 6.76E-08 | NB_I | |
| | X_position | Flatness_Ratio | -0.0176858 | 0.48557936 | NB_NI | NS |
| 144 | X_position | Instantaneous_Speed | 0.04506248 | 0.12136685 | B_I | |

| | | | | | | |
|---|---|---|---|---|---|---|
| | X_position | Instantaneous_Speed | 0.03938207 | 0.10332845 | B_NI | |
| | X_position | Instantaneous_Speed | 0.03406261 | 0.18041222 | NB_I | |
| | X_position | Instantaneous_Speed | 0.09509934 | 3.00E-05 | NB_NI | * |
| 145 | X_position | Max_Feret_Diameter | 0.06599186 | 0.05722848 | B_I | |
| | X_position | Max_Feret_Diameter | -0.1333242 | 3.62E-07 | B_NI | |
| | X_position | Max_Feret_Diameter | 0.12144617 | 1.15E-05 | NB_I | |
| | X_position | Max_Feret_Diameter | -0.0499802 | 0.04863134 | NB_NI | * |
| | X_position | Roundness_. | -0.1035516 | 0.00280229 | B_I | |
| 146 | X_position | Roundness_. | 0.12667029 | 1.35E-06 | B_NI | |
| | X_position | Roundness_. | -0.1066232 | 0.00011876 | NB_I | |
| | X_position | Roundness_. | -0.0587047 | 0.02052783 | NB_NI | * |
| | X_position | Sphericity_. | -0.0333858 | 0.33643444 | B_I | |
| 147 | X_position | Sphericity_. | 0.19262892 | 1.49E-13 | B_NI | |
| | X_position | Sphericity_. | 0.02583753 | 0.35230544 | NB_I | |
| | X_position | Sphericity_. | -0.0278392 | 0.27227668 | NB_NI | NS |
| 148 | X_position | Surface_Area | 0.0443208 | 0.20183435 | B_I | |
| | X_position | Surface_Area | -0.1798748 | 5.57E-12 | B_NI | |
| | X_position | Surface_Area | 0.05581634 | 0.04437107 | NB_I | |
| | X_position | Surface_Area | -0.0027638 | 0.91322746 | NB_NI | NS |
| 149 | X_position | Total_displacement | 0.0206707 | 0.47752534 | B_I | |
| | X_position | Total_displacement | -0.0301246 | 0.2128312 | B_NI | |
| | X_position | Total_displacement | 0.07201315 | 0.00458654 | NB_I | |
| | X_position | Total_displacement | 0.13246025 | 5.68E-09 | NB_NI | * |
| 150 | X_position | Total_distance | 0.0863812 | 0.00294444 | B_I | |
| | X_position | Total_distance | 0.00446414 | 0.85356201 | B_NI | |
| | X_position | Total_distance | 0.07270502 | 0.00420937 | NB_I | |
| | X_position | Total_distance | 0.11005192 | 1.34E-06 | NB_NI | * |
| 151 | X_position | Volume | 0.0427515 | 0.21828237 | B_I | |
| | X_position | Volume | -0.1345281 | 2.83E-07 | B_NI | |
| | X_position | Volume | 0.09962392 | 0.00032475 | NB_I | |
| | X_position | Volume | -0.0007061 | 0.97778961 | NB_NI | NS |
| | X_position | X_displacement | 0.06432801 | 0.02693214 | B_I | |
| 152 | X_position | X_displacement | 0.01664795 | 0.49121643 | B_NI | |
| | X_position | X_displacement | 0.00213749 | 0.93303172 | NB_I | |
| | X_position | X_displacement | 0.06167749 | 0.00686369 | NB_NI | * |
| 153 | X_position | X_size | 0.08198252 | 0.01809159 | B_I | |
| | X_position | X_size | -0.1391735 | 1.08E-07 | B_NI | |
| | X_position | X_size | 0.10176107 | 0.00024045 | NB_I | |
| | X_position | X_size | -0.0459238 | 0.07004771 | NB_NI | NS |
| 154 | X_position | Y_displacement | 0.0740719 | 0.01081892 | B_I | |
| | X_position | Y_displacement | 0.00808356 | 0.73820666 | B_NI | |
| | X_position | Y_displacement | 0.03349605 | 0.18777332 | NB_I | |
| | X_position | Y_displacement | -0.0193796 | 0.39604895 | NB_NI | NS |
| 155 | X_position | Y_position | 0.83897406 | 0 | B_I | |
| | X_position | Y_position | -0.7357979 | 0 | B_NI | |
| | X_position | Y_position | 0.61870921 | 0 | NB_I | |
| | X_position | Y_position | -0.8159056 | 0 | NB_NI | * |
| 156 | X_position | Y_size | -0.0269652 | 0.43757332 | B_I | |
| | X_position | Y_size | -0.0773162 | 0.00326193 | B_NI | |
| | X_position | Y_size | 0.07826997 | 0.0047799 | NB_I | |
| | X_position | Y_size | -0.1799357 | 8.50E-13 | NB_NI | * |
| 157 | X_position | Z_displacement | -0.0113994 | 0.69529587 | B_I | |
| | X_position | Z_displacement | 0.04928207 | 0.04146398 | B_NI | |
| | X_position | Z_displacement | 0.12368532 | 1.05E-06 | NB_I | |
| | X_position | Z_displacement | 0.13205031 | 6.33E-09 | NB_NI | * |
| 158 | X_position | Z_position | 0.51612884 | 0 | B_I | |
| | X_position | Z_position | 0.49193862 | 0 | B_NI | |
| | X_position | Z_position | 0.0371339 | 0.12883665 | NB_I | |

| | | | | | | |
|---|---|---|---|---|---|---|
| | X_position | Z_position | 0.47213495 | 0 | NB_NI | * |
| 159 | X_position | Z_size | 0.03177905 | 0.36021831 | B_I | |
| | X_position | Z_size | -0.0329464 | 0.21053777 | B_NI | |
| | X_position | Z_size | 0.01583759 | 0.56862412 | NB_I | |
| | X_position | Z_size | 0.25742665 | 0 | NB_NI | * |
| 160 | X_size | Convexity_. | -0.3825619 | 0 | B_I | |
| | X_size | Convexity_. | -0.5569238 | 0 | B_NI | |
| | X_size | Convexity_. | -0.6121866 | 0 | NB_I | |
| | X_size | Convexity_. | -0.5538925 | 0 | NB_NI | * |
| 161 | X_size | Ellipse_A | 0.25427777 | 9.95E-14 | B_I | |
| | X_size | Ellipse_A | 0.04948191 | 0.05995296 | B_NI | |
| | X_size | Ellipse_A | 0.10129439 | 0.00025688 | NB_I | |
| | X_size | Ellipse_A | 0.12858112 | 3.57E-07 | NB_NI | * |
| 162 | X_size | Ellipse_B | 0.0907517 | 0.00885581 | B_I | |
| | X_size | Ellipse_B | 0.00679059 | 0.79640628 | B_NI | |
| | X_size | Ellipse_B | 0.11066882 | 6.45E-05 | NB_I | |
| | X_size | Ellipse_B | 0.03719821 | 0.1423415 | NB_NI | NS |
| | X_size | Ellipse_C | -0.0333953 | 0.3362976 | B_I | |
| 163 | X_size | Ellipse_C | -0.0629407 | 0.01667903 | B_NI | |
| | X_size | Ellipse_C | -0.0244279 | 0.37920214 | NB_I | |
| | X_size | Ellipse_C | 0.02905183 | 0.25193061 | NB_NI | NS |
| | X_size | Elongation | 0.31479287 | 0 | B_I | |
| 164 | X_size | Elongation | 0.09758563 | 0.00020201 | B_NI | |
| | X_size | Elongation | 0.24878742 | 0 | NB_I | |
| | X_size | Elongation | 0.18992326 | 4.13E-14 | NB_NI | * |
| | X_size | Flatness_Ratio | 0.15256858 | 9.99E-06 | B_I | |
| 165 | X_size | Flatness_Ratio | 0.0375882 | 0.15311662 | B_NI | |
| | X_size | Flatness_Ratio | 0.26999488 | 0 | NB_I | |
| | X_size | Flatness_Ratio | 0.12684309 | 5.13E-07 | NB_NI | * |
| 166 | X_size | Max_Feret_Diameter | 0.98184139 | 0 | B_I | |
| | X_size | Max_Feret_Diameter | 0.98242795 | 0 | B_NI | |
| | X_size | Max_Feret_Diameter | 0.97633493 | 0 | NB_I | |
| | X_size | Max_Feret_Diameter | 0.99104005 | 0 | NB_NI | * |
| 167 | X_size | Roundness_. | -0.5832303 | 0 | B_I | |
| | X_size | Roundness_. | -0.6230311 | 0 | B_NI | |
| | X_size | Roundness_. | -0.6831214 | 0 | NB_I | |
| | X_size | Roundness_. | -0.6392599 | 0 | NB_NI | * |
| 168 | X_size | Sphericity_. | -0.4226823 | 0 | B_I | |
| | X_size | Sphericity_. | -0.6153148 | 0 | B_NI | |
| | X_size | Sphericity_. | -0.6493569 | 0 | NB_I | |
| | X_size | Sphericity_. | -0.6678365 | 0 | NB_NI | * |
| 169 | X_size | Surface_Area | 0.52985263 | 0 | B_I | |
| | X_size | Surface_Area | 0.66473925 | 0 | B_NI | |
| | X_size | Surface_Area | 0.67218614 | 0 | NB_I | |
| | X_size | Surface_Area | 0.72532225 | 0 | NB_NI | * |
| 170 | X_size | Volume | 0.56276989 | 0 | B_I | |
| | X_size | Volume | 0.59567428 | 0 | B_NI | |
| | X_size | Volume | 0.55646795 | 0 | NB_I | |
| | X_size | Volume | 0.59545761 | 0 | NB_NI | * |
| 171 | X_size | Y_size | 0.26347646 | 1.15E-14 | B_I | |
| | X_size | Y_size | 0.19329596 | 1.23E-13 | B_NI | |
| | X_size | Y_size | 0.22320573 | 4.44E-16 | NB_I | |
| | X_size | Y_size | 0.20940618 | 0 | NB_NI | * |
| 172 | X_size | Z_size | 0.28425169 | 0 | B_I | |
| | X_size | Z_size | 0.42753845 | 0 | B_NI | |
| | X_size | Z_size | 0.20247044 | 1.79E-13 | NB_I | |
| | X_size | Z_size | 0.43547565 | 0 | NB_NI | * |
| 173 | Y_displacement | Convexity_. | -0.1447994 | 3.81E-05 | B_I | |

| | | | | | | |
|---|---|---|---|---|---|---|
| | Y_displacement | Convexity_. | -0.0855964 | 0.00136876 | B_NI | |
| | Y_displacement | Convexity_. | -0.097208 | 0.00073151 | NB_I | |
| | Y_displacement | Convexity_. | -0.0947316 | 0.00023187 | NB_NI | * |
| 174 | Y_displacement | Directionality | 0.12689258 | 1.11E-05 | B_I | |
| | Y_displacement | Directionality | 0.0440213 | 0.06780354 | B_NI | |
| | Y_displacement | Directionality | 0.07915965 | 0.001546 | NB_I | |
| | Y_displacement | Directionality | -0.0212038 | 0.3494793 | NB_NI | NS |
| 175 | Y_displacement | Ellipse_A | 0.01847497 | 0.60114194 | B_I | |
| | Y_displacement | Ellipse_A | 0.02572329 | 0.33685425 | B_NI | |
| | Y_displacement | Ellipse_A | 0.010953 | 0.70418936 | NB_I | |
| | Y_displacement | Ellipse_A | -0.0093564 | 0.71675229 | NB_NI | NS |
| 176 | Y_displacement | Ellipse_B | -0.0102809 | 0.77113838 | B_I | |
| | Y_displacement | Ellipse_B | 0.00583325 | 0.82762028 | B_NI | |
| | Y_displacement | Ellipse_B | 0.07450783 | 0.00970308 | NB_I | |
| | Y_displacement | Ellipse_B | -0.0138178 | 0.59208887 | NB_NI | NS |
| 177 | Y_displacement | Ellipse_C | 0.03113598 | 0.37823805 | B_I | |
| | Y_displacement | Ellipse_C | 0.07214165 | 0.00700652 | B_NI | |
| | Y_displacement | Ellipse_C | 0.03446079 | 0.23214262 | NB_I | |
| | Y_displacement | Ellipse_C | 0.04960655 | 0.05426959 | NB_NI | NS |
| 180 | Y_displacement | Elongation | -0.0449796 | 0.20292674 | B_I | |
| | Y_displacement | Elongation | 0.04386621 | 0.10135972 | B_NI | |
| | Y_displacement | Elongation | 0.03822243 | 0.18504667 | NB_I | |
| | Y_displacement | Elongation | 0.00216536 | 0.93308681 | NB_NI | NS |
| 181 | Y_displacement | Flatness_Ratio | 0.01449951 | 0.6816169 | B_I | |
| | Y_displacement | Flatness_Ratio | 0.00109007 | 0.9675415 | B_NI | |
| | Y_displacement | Flatness_Ratio | 0.03807391 | 0.18676101 | NB_I | |
| 182 | Y_displacement | Flatness_Ratio | -0.0140726 | 0.58527801 | NB_NI | NS |
| | Y_displacement | Instantaneous_Speed | 0.52149123 | 0 | B_I | |
| | Y_displacement | Instantaneous_Speed | 0.52632517 | 0 | B_NI | |
| | Y_displacement | Instantaneous_Speed | 0.64084816 | 0 | NB_I | |
| | Y_displacement | Instantaneous_Speed | 0.5198853 | 0 | NB_NI | * |
| 183 | Y_displacement | Max_Feret_Diameter | -0.0145342 | 0.68089772 | B_I | |
| | Y_displacement | Max_Feret_Diameter | 0.05763865 | 0.03128578 | B_NI | |
| | Y_displacement | Max_Feret_Diameter | 0.0812816 | 0.00477092 | NB_I | |
| | Y_displacement | Max_Feret_Diameter | 0.04202956 | 0.10301351 | NB_NI | NS |
| 184 | Y_displacement | Roundness_. | -0.0704451 | 0.04597843 | B_I | |
| | Y_displacement | Roundness_. | -0.0519675 | 0.05222942 | B_NI | |
| | Y_displacement | Roundness_. | -0.1010589 | 0.00044481 | NB_I | |
| | Y_displacement | Roundness_. | 0.00543596 | 0.83305823 | NB_NI | NS |
| 185 | Y_displacement | Sphericity_. | -0.0379563 | 0.28269351 | B_I | |
| | Y_displacement | Sphericity_. | -0.0380872 | 0.15494058 | B_NI | |
| | Y_displacement | Sphericity_. | -0.0249217 | 0.38759611 | NB_I | |
| | Y_displacement | Sphericity_. | 0.0226771 | 0.37917493 | NB_NI | NS |
| 186 | Y_displacement | Surface_Area | 0.01307839 | 0.71134968 | B_I | |
| | Y_displacement | Surface_Area | 0.09155026 | 0.00061541 | B_NI | |
| | Y_displacement | Surface_Area | 0.04236401 | 0.14180323 | NB_I | |
| | Y_displacement | Surface_Area | 0.04346826 | 0.09174215 | NB_NI | NS |
| 187 | Y_displacement | Total_displacement | 0.1747483 | 1.25E-09 | B_I | |
| | Y_displacement | Total_displacement | 0.09050405 | 0.00016948 | B_NI | |
| | Y_displacement | Total_displacement | 0.20429765 | 0 | NB_I | |
| | Y_displacement | Total_displacement | 0.12415544 | 3.82E-08 | NB_NI | * |
| 188 | Y_displacement | Total_distance | 0.15025945 | 1.87E-07 | B_I | |
| | Y_displacement | Total_distance | 0.10816073 | 6.85E-06 | B_NI | |
| | Y_displacement | Total_distance | 0.20753777 | 0 | NB_I | |
| | Y_displacement | Total_distance | 0.16951357 | 4.93E-14 | NB_NI | * |
| 189 | Y_displacement | Volume | -0.0009728 | 0.97804191 | B_I | |
| | Y_displacement | Volume | 0.06672499 | 0.01264559 | B_NI | |
| | Y_displacement | Volume | 0.07537606 | 0.00888474 | NB_I | |

| | | | | | | |
|---|---|---|---|---|---|---|
| | Y_displacement | Volume | 0.0681618 | 0.00814375 | NB_NI | * |
| 190 | Y_displacement | X_size | -0.0264605 | 0.45398938 | B_I | |
| | Y_displacement | X_size | 0.0405003 | 0.13041149 | B_NI | |
| | Y_displacement | X_size | 0.05782078 | 0.04486688 | NB_I | |
| | Y_displacement | X_size | 0.01813021 | 0.48202057 | NB_NI | NS |
| 191 | Y_displacement | Y_size | 0.14841919 | 2.41E-05 | B_I | |
| | Y_displacement | Y_size | 0.14487532 | 5.42E-08 | B_NI | |
| | Y_displacement | Y_size | 0.14727399 | 2.85E-07 | NB_I | |
| | Y_displacement | Y_size | 0.22137728 | 0 | NB_NI | * |
| 192 | Y_displacement | Z_displacement | 0.28781745 | 0 | B_I | |
| | Y_displacement | Z_displacement | 0.2637693 | 0 | B_NI | |
| | Y_displacement | Z_displacement | 0.3762632 | 0 | NB_I | |
| | Y_displacement | Z_displacement | 0.21129616 | 0 | NB_NI | * |
| 193 | Y_displacement | Z_size | -0.0023125 | 0.94783318 | B_I | |
| | Y_displacement | Z_size | -0.0050837 | 0.84948691 | B_NI | |
| | Y_displacement | Z_size | 0.04350531 | 0.13136931 | NB_I | |
| | Y_displacement | Z_size | -0.0362407 | 0.15981511 | NB_NI | NS |
| 194 | Y_position | Convexity_. | -0.0090399 | 0.79470439 | B_I | |
| | Y_position | Convexity_. | -0.1557122 | 2.64E-09 | B_NI | |
| | Y_position | Convexity_. | 0.11761916 | 2.15E-05 | NB_I | |
| | Y_position | Convexity_. | -0.2700889 | 0 | NB_NI | * |
| 195 | Y_position | Directionality | 0.05303368 | 0.06823738 | B_I | |
| | Y_position | Directionality | 0.06001198 | 0.01300984 | B_NI | |
| | Y_position | Directionality | -0.0075893 | 0.76542866 | NB_I | |
| | Y_position | Directionality | -0.0081496 | 0.72118889 | NB_NI | NS |
| 196 | Y_position | Ellipse_A | 0.08635417 | 0.01276555 | B_I | |
| | Y_position | Ellipse_A | 0.08525268 | 0.00117471 | B_NI | |
| | Y_position | Ellipse_A | -0.0369336 | 0.18358235 | NB_I | |
| | Y_position | Ellipse_A | 0.07503712 | 0.00304976 | NB_NI | * |
| 197 | Y_position | Ellipse_B | -0.0290521 | 0.4029285 | B_I | |
| | Y_position | Ellipse_B | 0.03237888 | 0.2185073 | B_NI | |
| | Y_position | Ellipse_B | 0.01766566 | 0.52484774 | NB_I | |
| | Y_position | Ellipse_B | -0.0096262 | 0.70428518 | NB_NI | NS |
| 198 | Y_position | Ellipse_C | -0.0319521 | 0.35760668 | B_I | |
| | Y_position | Ellipse_C | 0.06309033 | 0.0164216 | B_NI | |
| | Y_position | Ellipse_C | 0.01991862 | 0.47337223 | NB_I | |
| | Y_position | Ellipse_C | 0.01906374 | 0.45223322 | NB_NI | NS |
| 199 | Y_position | Elongation | -0.085705 | 0.01345651 | B_I | |
| | Y_position | Elongation | 0.10925072 | 3.14E-05 | B_NI | |
| | Y_position | Elongation | -0.1827433 | 3.28E-11 | NB_I | |
| | Y_position | Elongation | 0.08434518 | 0.00086402 | NB_NI | * |
| 200 | Y_position | Flatness_Ratio | 0.0255332 | 0.46230332 | B_I | |
| | Y_position | Flatness_Ratio | 0.03683439 | 0.16153143 | B_NI | |
| | Y_position | Flatness_Ratio | 0.10855383 | 8.90E-05 | NB_I | |
| | Y_position | Flatness_Ratio | 0.03068462 | 0.22624353 | NB_NI | NS |
| 201 | Y_position | Instantaneous_Speed | 0.0432588 | 0.13701492 | B_I | |
| | Y_position | Instantaneous_Speed | -0.0180839 | 0.45460401 | B_NI | |
| | Y_position | Instantaneous_Speed | -0.0022828 | 0.92849196 | NB_I | |
| | Y_position | Instantaneous_Speed | -0.0561249 | 0.01390897 | NB_NI | * |
| 202 | Y_position | Max_Feret_Diameter | -0.0149821 | 0.66627651 | B_I | |
| | Y_position | Max_Feret_Diameter | 0.10997219 | 2.78E-05 | B_NI | |
| | Y_position | Max_Feret_Diameter | -0.1792944 | 7.71E-11 | NB_I | |
| | Y_position | Max_Feret_Diameter | 0.16270266 | 1.06E-10 | NB_NI | * |
| 203 | Y_position | Roundness_. | -0.0628714 | 0.0700694 | B_I | |
| | Y_position | Roundness_. | -0.0694854 | 0.00821307 | B_NI | |
| | Y_position | Roundness_. | 0.17548664 | 1.94E-10 | NB_I | |
| | Y_position | Roundness_. | -0.0586325 | 0.02068394 | NB_NI | * |
| 204 | Y_position | Sphericity_. | -0.1009857 | 0.00356609 | B_I | |

| | | | | | | |
|---|---|---|---|---|---|---|
| | Y_position | Sphericity_. | -0.0202206 | 0.44229151 | B_NI | |
| | Y_position | Sphericity_. | 0.18472792 | 1.99E-11 | NB_I | |
| | Y_position | Sphericity_. | -0.025589 | 0.31294254 | NB_NI | NS |
| 205 | Y_position | Surface_Area | 0.13203648 | 0.00013496 | B_I | |
| | Y_position | Surface_Area | 0.07632051 | 0.0036853 | B_NI | |
| | Y_position | Surface_Area | -0.0315067 | 0.2566665 | NB_I | |
| | Y_position | Surface_Area | 0.12202046 | 1.37E-06 | NB_NI | * |
| 206 | Y_position | Total_displacement | 0.07783048 | 0.00740202 | B_I | |
| | Y_position | Total_displacement | 0.00473258 | 0.8448646 | B_NI | |
| | Y_position | Total_displacement | -0.0421126 | 0.09766057 | NB_I | |
| | Y_position | Total_displacement | -0.0901916 | 7.57E-05 | NB_NI | * |
| 207 | Y_position | Total_distance | 0.0614103 | 0.03469043 | B_I | |
| | Y_position | Total_distance | 0.02574391 | 0.28706297 | B_NI | |
| | Y_position | Total_distance | 0.0203432 | 0.42380812 | NB_I | |
| | Y_position | Total_distance | -0.032504 | 0.15453259 | NB_NI | NS |
| 208 | Y_position | Volume | 0.09143674 | 0.00835412 | B_I | |
| | Y_position | Volume | 0.11481627 | 1.20E-05 | B_NI | |
| | Y_position | Volume | 0.07260011 | 0.00888225 | NB_I | |
| | Y_position | Volume | 0.15233234 | 1.53E-09 | NB_NI | * |
| 209 | Y_position | X_displacement | 0.04213508 | 0.14752272 | B_I | |
| | Y_position | X_displacement | 0.02289409 | 0.34378898 | B_NI | |
| | Y_position | X_displacement | -0.0514841 | 0.04283343 | NB_I | |
| | Y_position | X_displacement | -0.0330739 | 0.14742689 | NB_NI | NS |
| 210 | Y_position | X_size | -0.0014483 | 0.96674755 | B_I | |
| | Y_position | X_size | 0.09052467 | 0.00056807 | B_NI | |
| | Y_position | X_size | -0.139461 | 4.55E-07 | NB_I | |
| | Y_position | X_size | 0.14638963 | 6.51E-09 | NB_NI | * |
| 211 | Y_position | Y_displacement | 0.04239488 | 0.14504029 | B_I | |
| | Y_position | Y_displacement | -0.00231 | 0.92391121 | B_NI | |
| | Y_position | Y_displacement | 0.04688012 | 0.06518241 | NB_I | |
| | Y_position | Y_displacement | 0.04048505 | 0.07613965 | NB_NI | NS |
| 212 | Y_position | Y_size | -0.0403639 | 0.24511526 | B_I | |
| | Y_position | Y_size | 0.22867066 | 0 | B_NI | |
| | Y_position | Y_size | -0.1855625 | 1.61E-11 | NB_I | |
| | Y_position | Y_size | 0.30785042 | 0 | NB_NI | * |
| 213 | Y_position | Z_displacement | 0.01568198 | 0.58999663 | B_I | |
| | Y_position | Z_displacement | -0.0573159 | 0.01770453 | B_NI | |
| | Y_position | Z_displacement | 0.16456009 | 7.33E-11 | NB_I | |
| | Y_position | Z_displacement | -0.108559 | 1.86E-06 | NB_NI | * |
| 214 | Y_position | Z_position | 0.58845812 | 0 | B_I | |
| | Y_position | Z_position | -0.435741 | 0 | B_NI | |
| | Y_position | Z_position | 0.42776811 | 0 | NB_I | |
| | Y_position | Z_position | -0.5887895 | 0 | NB_NI | * |
| 215 | Y_position | Z_size | 0.14932591 | 1.54E-05 | B_I | |
| | Y_position | Z_size | -0.1097955 | 2.86E-05 | B_NI | |
| | Y_position | Z_size | 0.22519164 | 2.22E-16 | NB_I | |
| | Y_position | Z_size | -0.2251239 | 0 | NB_NI | * |
| 216 | Y_size | Convexity_. | -0.3643106 | 0 | B_I | |
| | Y_size | Convexity_. | -0.5488325 | 0 | B_NI | |
| | Y_size | Convexity_. | -0.4267379 | 0 | NB_I | |
| | Y_size | Convexity_. | -0.6358825 | 0 | NB_NI | * |
| 217 | Y_size | Ellipse_A | 0.09002703 | 0.0094156 | B_I | |
| | Y_size | Ellipse_A | 0.07844363 | 0.00283632 | B_NI | |
| | Y_size | Ellipse_A | -0.003166 | 0.90927412 | NB_I | |
| | Y_size | Ellipse_A | 0.06383684 | 0.01175301 | NB_NI | * |
| 218 | Y_size | Ellipse_B | 0.09391559 | 0.00674404 | B_I | |
| | Y_size | Ellipse_B | 0.03893462 | 0.13892113 | B_NI | |
| | Y_size | Ellipse_B | 0.09798172 | 0.00040751 | NB_I | |

| | | | | | | |
|---|---|---|---|---|---|---|
| | Y_size | Ellipse_B | 0.02956524 | 0.2436436 | NB_NI | NS |
| 219 | Y_size | Ellipse_C | 0.12637667 | 0.00025996 | B_I | |
| | Y_size | Ellipse_C | 0.26454815 | 0 | B_NI | |
| | Y_size | Ellipse_C | 0.06165793 | 0.02632584 | NB_I | |
| | Y_size | Ellipse_C | 0.18170646 | 5.03E-13 | NB_NI | * |
| 220 | Y_size | Elongation | 0.02406265 | 0.48849044 | B_I | |
| | Y_size | Elongation | 0.10789913 | 3.93E-05 | B_NI | |
| | Y_size | Elongation | -0.0846335 | 0.00227554 | NB_I | |
| | Y_size | Elongation | 0.03444101 | 0.17436414 | NB_NI | NS |
| 221 | Y_size | Flatness_Ratio | 0.11258338 | 0.00115038 | B_I | |
| | Y_size | Flatness_Ratio | 0.03643921 | 0.16607997 | B_NI | |
| | Y_size | Flatness_Ratio | 0.08832963 | 0.00144513 | NB_I | |
| | Y_size | Flatness_Ratio | 0.06831978 | 0.00700081 | NB_NI | * |
| 222 | Y_size | Max_Feret_Diameter | 0.39248237 | 0 | B_I | |
| | Y_size | Max_Feret_Diameter | 0.33096346 | 0 | B_NI | |
| | Y_size | Max_Feret_Diameter | 0.396231 | 0 | NB_I | |
| | Y_size | Max_Feret_Diameter | 0.30214635 | 0 | NB_NI | * |
| 223 | Y_size | Roundness_. | -0.0703514 | 0.04261296 | B_I | |
| | Y_size | Roundness_. | -0.302013 | 0 | B_NI | |
| | Y_size | Roundness_. | -0.3047717 | 0 | NB_I | |
| | Y_size | Roundness_. | -0.1679076 | 2.60E-11 | NB_NI | * |
| 224 | Y_size | Sphericity_. | -0.0185265 | 0.59382211 | B_I | |
| | Y_size | Sphericity_. | -0.3087258 | 0 | B_NI | |
| | Y_size | Sphericity_. | -0.1919978 | 3.05E-12 | NB_I | |
| | Y_size | Sphericity_. | -0.1727445 | 6.77E-12 | NB_NI | * |
| 225 | Y_size | Surface_Area | 0.18916206 | 3.92E-08 | B_I | |
| | Y_size | Surface_Area | 0.47028765 | 0 | B_NI | |
| | Y_size | Surface_Area | 0.34576344 | 0 | NB_I | |
| | Y_size | Surface_Area | 0.41947472 | 0 | NB_NI | * |
| 226 | Y_size | Volume | 0.3265523 | 0 | B_I | |
| | Y_size | Volume | 0.62389576 | 0 | B_NI | |
| | Y_size | Volume | 0.47104338 | 0 | NB_I | |
| | Y_size | Volume | 0.4974491 | 0 | NB_NI | * |
| 227 | Y_size | Z_size | 0.06965223 | 0.04471926 | B_I | |
| | Y_size | Z_size | 0.2160597 | 0 | B_NI | |
| | Y_size | Z_size | 0.01259492 | 0.65029851 | NB_I | |
| | Y_size | Z_size | 0.18757248 | 8.53E-14 | NB_NI | * |
| 228 | Z_displacement | Convexity_. | -0.0898156 | 0.0108868 | B_I | |
| | Z_displacement | Convexity_. | -0.0476061 | 0.07538324 | B_NI | |
| | Z_displacement | Convexity_. | -0.0894107 | 0.0019 | NB_I | |
| | Z_displacement | Convexity_. | -0.0385375 | 0.13495465 | NB_NI | NS |
| 229 | Z_displacement | Directionality | 0.09198988 | 0.00147583 | B_I | |
| | Z_displacement | Directionality | 0.06169264 | 0.01044798 | B_NI | |
| | Z_displacement | Directionality | 0.0459391 | 0.06645057 | NB_I | |
| | Z_displacement | Directionality | 0.0170577 | 0.45167472 | NB_NI | NS |
| 230 | Z_displacement | Ellipse_A | -0.013493 | 0.70262764 | B_I | |
| | Z_displacement | Ellipse_A | -0.0165861 | 0.53578751 | B_NI | |
| | Z_displacement | Ellipse_A | 0.05958635 | 0.03870982 | NB_I | |
| | Z_displacement | Ellipse_A | -0.0099678 | 0.69911752 | NB_NI | NS |
| 231 | Z_displacement | Ellipse_B | -0.0188742 | 0.59330108 | B_I | |
| | Z_displacement | Ellipse_B | -0.0050193 | 0.85136937 | B_NI | |
| | Z_displacement | Ellipse_B | 0.00424632 | 0.88298246 | NB_I | |
| | Z_displacement | Ellipse_B | 0.0049412 | 0.84805863 | NB_NI | NS |
| 232 | Z_displacement | Ellipse_C | -0.0485556 | 0.16925533 | B_I | |
| | Z_displacement | Ellipse_C | -0.0004599 | 0.98630304 | B_NI | |
| | Z_displacement | Ellipse_C | -0.005771 | 0.84144804 | NB_I | |
| | Z_displacement | Ellipse_C | 0.0464601 | 0.07147255 | NB_NI | NS |
| 233 | Z_displacement | Elongation | -0.0692301 | 0.04986862 | B_I | |

| | | | | | | |
|---|---|---|---|---|---|---|
| | Z_displacement | Elongation | -0.0273432 | 0.30730133 | B_NI | |
| | Z_displacement | Elongation | 0.00828311 | 0.77402081 | NB_I | |
| | Z_displacement | Elongation | -0.0249036 | 0.33415011 | NB_NI | NS |
| 234 | Z_displacement | Flatness_Ratio | 0.00366485 | 0.91741517 | B_I | |
| | Z_displacement | Flatness_Ratio | -0.00381 | 0.88690232 | B_NI | |
| | Z_displacement | Flatness_Ratio | 0.07493181 | 0.00929543 | NB_I | |
| | Z_displacement | Flatness_Ratio | -0.0007769 | 0.97596719 | NB_NI | NS |
| 235 | Z_displacement | Instantaneous_Speed | 0.76406467 | 0 | B_I | |
| | Z_displacement | Instantaneous_Speed | 0.71656597 | 0 | B_NI | |
| | Z_displacement | Instantaneous_Speed | 0.54354805 | 0 | NB_I | |
| | Z_displacement | Instantaneous_Speed | 0.58212286 | 0 | NB_NI | * |
| 236 | Z_displacement | Max_Feret_Diameter | -0.0107798 | 0.76036498 | B_I | |
| | Z_displacement | Max_Feret_Diameter | 0.01485513 | 0.57919087 | B_NI | |
| | Z_displacement | Max_Feret_Diameter | 0.0603302 | 0.03633915 | NB_I | |
| | Z_displacement | Max_Feret_Diameter | 0.02520291 | 0.32837217 | NB_NI | NS |
| 237 | Z_displacement | Roundness_. | -0.0456222 | 0.19654223 | B_I | |
| | Z_displacement | Roundness_. | -0.0619545 | 0.02061504 | B_NI | |
| | Z_displacement | Roundness_. | -0.0852019 | 0.00308935 | NB_I | |
| | Z_displacement | Roundness_. | -0.0033425 | 0.89687854 | NB_NI | NS |
| 238 | Z_displacement | Sphericity_. | -0.0275043 | 0.43637414 | B_I | |
| | Z_displacement | Sphericity_. | -0.0347377 | 0.19458451 | B_NI | |
| | Z_displacement | Sphericity_. | -0.0418702 | 0.14650993 | NB_I | |
| | Z_displacement | Sphericity_. | -0.026351 | 0.30680988 | NB_NI | NS |
| 239 | Z_displacement | Surface_Area | 0.00859762 | 0.80780557 | B_I | |
| | Z_displacement | Surface_Area | 0.02056029 | 0.44273153 | B_NI | |
| | Z_displacement | Surface_Area | 0.02568517 | 0.37321721 | NB_I | |
| | Z_displacement | Surface_Area | 0.05486646 | 0.03325008 | NB_NI | * |
| 240 | Z_displacement | Total_displacement | 0.36223918 | 0 | B_I | |
| | Z_displacement | Total_displacement | 0.19754413 | 0 | B_NI | |
| | Z_displacement | Total_displacement | 0.09809548 | 8.62E-05 | NB_I | |
| | Z_displacement | Total_displacement | 0.12118422 | 8.04E-08 | NB_NI | * |
| 241 | Z_displacement | Total_distance | 0.19473867 | 1.19E-11 | B_I | |
| | Z_displacement | Total_distance | 0.112863 | 2.67E-06 | B_NI | |
| | Z_displacement | Total_distance | 0.24061663 | 0 | NB_I | |
| | Z_displacement | Total_distance | 0.14763784 | 5.77E-11 | NB_NI | * |
| 242 | Z_displacement | Volume | 0.01434855 | 0.68475285 | B_I | |
| | Z_displacement | Volume | 0.02080246 | 0.43737608 | B_NI | |
| | Z_displacement | Volume | 0.02058515 | 0.47546737 | NB_I | |
| | Z_displacement | Volume | 0.06899006 | 0.00740012 | NB_NI | * |
| 243 | Z_displacement | X_size | -0.0036861 | 0.91693758 | B_I | |
| | Z_displacement | X_size | 0.01332763 | 0.61881012 | B_NI | |
| | Z_displacement | X_size | 0.05210233 | 0.07072597 | NB_I | |
| | Z_displacement | X_size | 0.01323877 | 0.6077025 | NB_NI | NS |
| 244 | Z_displacement | Y_size | 0.00843623 | 0.81134475 | B_I | |
| | Z_displacement | Y_size | 0.00547013 | 0.83819777 | B_NI | |
| | Z_displacement | Y_size | 0.04386859 | 0.12817595 | NB_I | |
| | Z_displacement | Y_size | 0.04023715 | 0.11856449 | NB_NI | NS |
| 245 | Z_displacement | Z_size | 0.06398207 | 0.06996939 | B_I | |
| | Z_displacement | Z_size | 0.0929959 | 0.00050321 | B_NI | |
| | Z_displacement | Z_size | 0.1239402 | 1.61E-05 | NB_I | |
| | Z_displacement | Z_size | 0.10521815 | 4.29E-05 | NB_NI | * |
| 246 | Z_position | Convexity_. | -0.104123 | 0.00265394 | B_I | |
| | Z_position | Convexity_. | 0.17853464 | 8.02E-12 | B_NI | |
| | Z_position | Convexity_. | -0.0542645 | 0.05063202 | NB_I | |
| | Z_position | Convexity_. | 0.21044987 | 0 | NB_NI | * |
| 247 | Z_position | Directionality | -0.0143231 | 0.62261821 | B_I | |
| | Z_position | Directionality | 0.02810295 | 0.24516364 | B_NI | |
| | Z_position | Directionality | -0.143975 | 1.27E-08 | NB_I | |

| | | | | | | |
|---|---|---|---|---|---|---|
| | Z_position | Directionality | 0.03219118 | 0.15854161 | NB_NI | NS |
| 248 | Z_position | Ellipse_A | 0.09299795 | 0.00730438 | B_I | |
| | Z_position | Ellipse_A | -0.0405517 | 0.12323612 | B_NI | |
| | Z_position | Ellipse_A | 0.01316404 | 0.63561792 | NB_I | |
| | Z_position | Ellipse_A | -0.0592184 | 0.0194462 | NB_NI | * |
| 249 | Z_position | Ellipse_B | -0.0254337 | 0.46405013 | B_I | |
| | Z_position | Ellipse_B | -0.0059074 | 0.82241166 | B_NI | |
| | Z_position | Ellipse_B | -0.0063031 | 0.82052385 | NB_I | |
| | Z_position | Ellipse_B | 0.02321842 | 0.3598979 | NB_NI | NS |
| 250 | Z_position | Ellipse_C | -0.0758459 | 0.02879548 | B_I | |
| | Z_position | Ellipse_C | 0.00801493 | 0.76073212 | B_NI | |
| | Z_position | Ellipse_C | -0.0030909 | 0.91141573 | NB_I | |
| | Z_position | Ellipse_C | 0.01094496 | 0.66607483 | NB_NI | NS |
| 251 | Z_position | Elongation | -0.0161578 | 0.6418508 | B_I | |
| | Z_position | Elongation | -0.0357673 | 0.17403363 | B_NI | |
| | Z_position | Elongation | -0.1127506 | 4.68E-05 | NB_I | |
| | Z_position | Elongation | -0.0556735 | 0.02803824 | NB_NI | * |
| 252 | Z_position | Flatness_Ratio | 0.00896315 | 0.79640871 | B_I | |
| | Z_position | Flatness_Ratio | -0.0070886 | 0.78768117 | B_NI | |
| | Z_position | Flatness_Ratio | 0.06834295 | 0.01378755 | NB_I | |
| | Z_position | Flatness_Ratio | -0.0092488 | 0.71536507 | NB_NI | NS |
| 253 | Z_position | Instantaneous_Speed | 0.00200885 | 0.94497293 | B_I | |
| | Z_position | Instantaneous_Speed | 0.08923525 | 0.00021824 | B_NI | |
| | Z_position | Instantaneous_Speed | -0.04001 | 0.11559438 | NB_I | |
| | Z_position | Instantaneous_Speed | -0.0119224 | 0.60160636 | NB_NI | NS |
| 254 | Z_position | Max_Feret_Diameter | 0.00830997 | 0.81095398 | B_I | |
| | Z_position | Max_Feret_Diameter | -0.0371277 | 0.15821706 | B_NI | |
| | Z_position | Max_Feret_Diameter | -0.1623901 | 4.00E-09 | NB_I | |
| | Z_position | Max_Feret_Diameter | -0.0345039 | 0.1735784 | NB_NI | NS |
| 255 | Z_position | Roundness_. | -0.2280962 | 2.87E-11 | B_I | |
| | Z_position | Roundness_. | 0.04303877 | 0.10185046 | B_NI | |
| | Z_position | Roundness_. | 0.09884749 | 0.0003617 | NB_I | |
| | Z_position | Roundness_. | -8.27E-05 | 0.9973993 | NB_NI | NS |
| 256 | Z_position | Sphericity_. | -0.3983388 | 0 | B_I | |
| | Z_position | Sphericity_. | -0.0356408 | 0.17556245 | B_NI | |
| | Z_position | Sphericity_. | -0.2918854 | 0 | NB_I | |
| | Z_position | Sphericity_. | -0.1598596 | 2.24E-10 | NB_NI | * |
| 257 | Z_position | Surface_Area | 0.26030105 | 2.44E-14 | B_I | |
| | Z_position | Surface_Area | -0.0083421 | 0.7512819 | B_NI | |
| | Z_position | Surface_Area | 0.2312552 | 0 | NB_I | |
| | Z_position | Surface_Area | 0.03112429 | 0.21965746 | NB_NI | NS |
| 258 | Z_position | Total_displacement | 0.05033485 | 0.083537 | B_I | |
| | Z_position | Total_displacement | 0.23790011 | 0 | B_NI | |
| | Z_position | Total_displacement | -0.2866096 | 0 | NB_I | |
| | Z_position | Total_displacement | -0.0836011 | 0.00024517 | NB_NI | * |
| 259 | Z_position | Total_distance | -0.0102209 | 0.72545129 | B_I | |
| | Z_position | Total_distance | 0.10609356 | 1.09E-05 | B_NI | |
| | Z_position | Total_distance | -0.1781352 | 1.67E-12 | NB_I | |
| | Z_position | Total_distance | -0.0796799 | 0.00047463 | NB_NI | * |
| 260 | Z_position | Volume | -0.0492868 | 0.15574796 | B_I | |
| | Z_position | Volume | -0.1379517 | 1.39E-07 | B_NI | |
| | Z_position | Volume | 0.07946566 | 0.00417348 | NB_I | |
| | Z_position | Volume | -0.0448271 | 0.07701185 | NB_NI | NS |
| 261 | Z_position | X_displacement | -0.0179786 | 0.53673039 | B_I | |
| | Z_position | X_displacement | 0.05741238 | 0.01751391 | B_NI | |
| | Z_position | X_displacement | -0.047898 | 0.05955321 | NB_I | |
| | Z_position | X_displacement | -0.0315565 | 0.16691613 | NB_NI | NS |
| 262 | Z_position | X_size | 0.02735097 | 0.43104252 | B_I | |

| | | | | | | |
|---|---|---|---|---|---|---|
| | Z_position | X_size | 0.00355118 | 0.89267347 | B_NI | |
| | Z_position | X_size | -0.1078543 | 9.89E-05 | NB_I | |
| | Z_position | X_size | -0.020321 | 0.4229685 | NB_NI | NS |
| 263 | Z_position | Y_displacement | -0.0441797 | 0.12884203 | B_I | |
| | Z_position | Y_displacement | 0.00881407 | 0.71553359 | B_NI | |
| | Z_position | Y_displacement | -0.054363 | 0.03245637 | NB_I | |
| | Z_position | Y_displacement | -0.1074427 | 2.38E-06 | NB_NI | * |
| 264 | Z_position | Y_size | -0.2806962 | 0 | B_I | |
| | Z_position | Y_size | -0.2963332 | 0 | B_NI | |
| | Z_position | Y_size | -0.3568068 | 0 | NB_I | |
| | Z_position | Y_size | -0.3283893 | 0 | NB_NI | * |
| 265 | Z_position | Z_displacement | 0.02875642 | 0.32304036 | B_I | |
| | Z_position | Z_displacement | 0.08010295 | 0.000909 | B_NI | |
| | Z_position | Z_displacement | 0.06038779 | 0.01749271 | NB_I | |
| | Z_position | Z_displacement | 0.07029294 | 0.00205691 | NB_NI | * |
| 266 | Z_position | Z_size | 0.18241729 | 1.19E-07 | B_I | |
| | Z_position | Z_size | 0.05190779 | 0.04844032 | B_NI | |
| | Z_position | Z_size | 0.46914664 | 0 | NB_I | |
| | Z_position | Z_size | 0.38170755 | 0 | NB_NI | * |
| 267 | Z_size | Convexity_. | -0.5765371 | 0 | B_I | |
| | Z_size | Convexity_. | -0.6411263 | 0 | B_NI | |
| | Z_size | Convexity_. | -0.5395641 | 0 | NB_I | |
| | Z_size | Convexity_. | -0.516305 | 0 | NB_NI | * |
| 268 | Z_size | Ellipse_A | 0.08065324 | 0.02005668 | B_I | |
| | Z_size | Ellipse_A | 0.02990956 | 0.25569677 | B_NI | |
| | Z_size | Ellipse_A | 0.08310807 | 0.002731 | NB_I | |
| | Z_size | Ellipse_A | 0.04867108 | 0.05484529 | NB_NI | NS |
| 269 | Z_size | Ellipse_B | -0.0210209 | 0.5450958 | B_I | |
| | Z_size | Ellipse_B | -0.0178057 | 0.49868961 | B_NI | |
| | Z_size | Ellipse_B | 0.11762238 | 2.15E-05 | NB_I | |
| | Z_size | Ellipse_B | 0.05319412 | 0.03583644 | NB_NI | * |
| 270 | Z_size | Ellipse_C | -0.0118693 | 0.73261033 | B_I | |
| | Z_size | Ellipse_C | 0.07249799 | 0.00581426 | B_NI | |
| | Z_size | Ellipse_C | 0.09443013 | 0.00065815 | NB_I | |
| | Z_size | Ellipse_C | 0.09210499 | 0.00027351 | NB_NI | * |
| 271 | Z_size | Elongation | -0.1764559 | 3.07E-07 | B_I | |
| | Z_size | Elongation | -0.0596577 | 0.02329128 | B_NI | |
| | Z_size | Elongation | -0.1280637 | 3.68E-06 | NB_I | |
| | Z_size | Elongation | -0.0070911 | 0.7797959 | NB_NI | NS |
| 272 | Z_size | Flatness_Ratio | 0.07809864 | 0.02435986 | B_I | |
| | Z_size | Flatness_Ratio | 0.00638346 | 0.80836812 | B_NI | |
| | Z_size | Flatness_Ratio | 0.32977107 | 0 | NB_I | |
| | Z_size | Flatness_Ratio | 0.12612966 | 5.95E-07 | NB_NI | * |
| 273 | Z_size | Max_Feret_Diameter | 0.30630404 | 0 | B_I | |
| | Z_size | Max_Feret_Diameter | 0.46684104 | 0 | B_NI | |
| | Z_size | Max_Feret_Diameter | 0.19839235 | 5.49E-13 | NB_I | |
| | Z_size | Max_Feret_Diameter | 0.46546972 | 0 | NB_NI | * |
| 274 | Z_size | Roundness_. | -0.3716531 | 0 | B_I | |
| | Z_size | Roundness_. | -0.3728855 | 0 | B_NI | |
| | Z_size | Roundness_. | -0.2274328 | 0 | NB_I | |
| | Z_size | Roundness_. | -0.3485386 | 0 | NB_NI | * |
| 275 | Z_size | Sphericity_. | -0.6032296 | 0 | B_I | |
| | Z_size | Sphericity_. | -0.5408847 | 0 | B_NI | |
| | Z_size | Sphericity_. | -0.5511211 | 0 | NB_I | |
| | Z_size | Sphericity_. | -0.6347367 | 0 | NB_NI | * |
| 276 | Z_size | Surface_Area | 0.65023386 | 0 | B_I | |
| | Z_size | Surface_Area | 0.55742836 | 0 | B_NI | |
| | Z_size | Surface_Area | 0.59497285 | 0 | NB_I | |

| | | | | | | |
|---|---|---|---|---|---|---|
| | Z_size | Surface_Area | 0.65235782 | 0 | NB_NI | * |
| 277 | Z_size | Volume | 0.54021883 | 0 | B_I | |
| | Z_size | Volume | 0.5812946 | 0 | B_NI | |
| | Z_size | Volume | 0.50244659 | 0 | NB_I | |
| | Z_size | Volume | 0.5400306 | 0 | NB_NI | * |

**Table S4**

| | pvalue | measurement | coefficient | Significance |
|---|---|---|---|---|
| Dataset | 2.99E-49 | Convexity_. | -0.2684621 | * |
| Blebbistatin:Injured | 0.00074531 | Convexity_. | -0.2029283 | * |
| Injured | 0.04132771 | Convexity_. | -0.0810242 | * |
| Time | 0.00292086 | Convexity_. | 0.04326495 | * |
| Intercept | 6.78E-28 | Convexity_. | 0.36132029 | * |
| Blebbistatin | 3.14E-32 | Convexity_. | 0.78355844 | * |
| Time | 4.77E-59 | Directionality | -0.235059 | * |
| Blebbistatin:Injured | 0.00118093 | Directionality | -0.1919919 | * |
| Intercept | 7.61E-05 | Directionality | -0.1277067 | * |
| Dataset | 0.06196489 | Directionality | -0.0330185 | |
| Blebbistatin | 0.02741446 | Directionality | 0.14277386 | * |
| Injured | 3.34E-27 | Directionality | 0.42470977 | * |
| Blebbistatin | 9.69E-05 | Ellipse_A | -0.2610483 | * |
| Intercept | 0.01409592 | Ellipse_A | -0.0818779 | * |
| Time | 1.37E-07 | Ellipse_A | -0.0780023 | * |
| Dataset | 0.0133774 | Ellipse_A | 0.04524616 | * |
| Blebbistatin:Injured | 0.12478215 | Ellipse_A | 0.09390008 | |
| Injured | 5.09E-06 | Ellipse_A | 0.18439439 | * |
| Blebbistatin | 9.81E-11 | Ellipse_B | -0.4351633 | * |
| Intercept | 2.89E-06 | Ellipse_B | -0.1566485 | * |
| Time | 0.01634314 | Ellipse_B | -0.0356024 | * |
| Blebbistatin:Injured | 0.10292861 | Ellipse_B | 0.1000522 | |
| Injured | 0.00504113 | Ellipse_B | 0.11362825 | * |
| Dataset | 6.04E-11 | Ellipse_B | 0.12027468 | * |
| Blebbistatin:Injured | 2.58E-09 | Ellipse_C | -0.3619508 | * |
| Intercept | 4.21E-05 | Ellipse_C | -0.1355418 | * |
| Dataset | 0.5557221 | Ellipse_C | -0.0106837 | |
| Time | 0.56351725 | Ellipse_C | -0.0084654 | |
| Injured | 0.9778472 | Ellipse_C | -0.0011119 | |
| Blebbistatin | 3.20E-13 | Ellipse_C | 0.48482055 | * |
| Dataset | 0.00034001 | Elongation | -0.0662163 | * |
| Blebbistatin:Injured | 0.4425107 | Elongation | -0.0474438 | |
| Time | 0.09081346 | Elongation | -0.0252483 | |
| Injured | 0.25645335 | Elongation | 0.04628952 | |
| Intercept | 0.13428603 | Elongation | 0.05043436 | |
| Blebbistatin | 0.00130735 | Elongation | 0.21735071 | * |
| Blebbistatin | 1.21E-30 | Flatness_Ratio | -0.7634366 | * |
| Intercept | 0.16643402 | Flatness_Ratio | -0.0454157 | |
| Time | 0.01303836 | Flatness_Ratio | -0.0361272 | * |
| Injured | 0.01939642 | Flatness_Ratio | 0.09294819 | * |
| Dataset | 1.51E-11 | Flatness_Ratio | 0.12172796 | * |
| Blebbistatin:Injured | 1.34E-09 | Flatness_Ratio | 0.36572358 | * |
| Blebbistatin | 3.69E-25 | Instantaneous_Speed | -0.6743438 | * |
| Intercept | 0.04869588 | Instantaneous_Speed | -0.0635599 | * |

| | | | | |
|---|---|---|---|---|
| Dataset | 3.26E-07 | Instantaneous_Speed | 0.09043614 | * |
| Time | 8.61E-19 | Instantaneous_Speed | 0.12703957 | * |
| Injured | 3.44E-11 | Instantaneous_Speed | 0.25934927 | * |
| Blebbistatin:Injured | 3.15E-06 | Instantaneous_Speed | 0.27593209 | * |
| Blebbistatin | 6.51E-13 | Max_Feret_Diameter | -0.4799751 | * |
| Time | 3.52E-13 | Max_Feret_Diameter | -0.1072572 | * |
| Intercept | 0.47186004 | Max_Feret_Diameter | -0.0238677 | |
| Dataset | 2.86E-05 | Max_Feret_Diameter | 0.07620236 | * |
| Injured | 0.0496869 | Max_Feret_Diameter | 0.07887268 | * |
| Blebbistatin:Injured | 0.01049105 | Max_Feret_Diameter | 0.15579541 | * |
| Blebbistatin:Injured | 0.00058895 | Roundness_. | -0.2095222 | * |
| Dataset | 2.41E-16 | Roundness_. | -0.1499146 | * |
| Injured | 0.21626088 | Roundness_. | 0.04974518 | |
| Intercept | 0.11404607 | Roundness_. | 0.05249321 | |
| Time | 2.95E-05 | Roundness_. | 0.06156378 | * |
| Blebbistatin | 2.09E-26 | Roundness_. | 0.71319268 | * |
| Blebbistatin:Injured | 6.50E-16 | Sphericity_. | -0.4834809 | * |
| Dataset | 5.05E-61 | Sphericity_. | -0.2982664 | * |
| Time | 0.07579286 | Sphericity_. | -0.0255858 | |
| Injured | 0.12345878 | Sphericity_. | 0.06064622 | |
| Intercept | 1.40E-23 | Sphericity_. | 0.3271167 | * |
| Blebbistatin | 6.85E-51 | Sphericity_. | 0.9910482 | * |
| Blebbistatin | 1.44E-22 | Surface_Area | -0.6495598 | * |
| Intercept | 1.24E-25 | Surface_Area | -0.3468232 | * |
| Time | 0.00327822 | Surface_Area | -0.0429498 | * |
| Blebbistatin:Injured | 0.01552902 | Surface_Area | 0.14625473 | * |
| Injured | 0.00015428 | Surface_Area | 0.15107624 | * |
| Dataset | 2.61E-36 | Surface_Area | 0.22931805 | * |
| Blebbistatin | 2.09E-18 | Total_displacement | -0.470841 | * |
| Intercept | 2.53E-06 | Total_displacement | -0.1257619 | * |
| Dataset | 0.00952721 | Total_displacement | 0.03796736 | * |
| Blebbistatin:Injured | 0.21677069 | Total_displacement | 0.06048831 | |
| Time | 0 | Total_displacement | 0.50049162 | * |
| Injured | 6.40E-70 | Total_displacement | 0.58157773 | * |
| Intercept | 1.24E-13 | Volume | -0.2472828 | * |
| Blebbistatin:Injured | 0.00222076 | Volume | -0.1867291 | * |
| Blebbistatin | 0.01481887 | Volume | -0.1626877 | * |
| Time | 5.99E-12 | Volume | -0.1016939 | * |
| Dataset | 1.13E-07 | Volume | 0.09691294 | * |
| Injured | 1.84E-12 | Volume | 0.28460101 | * |
| Blebbistatin | 1.48E-17 | X_displacement | -0.5629057 | * |
| Intercept | 0.48059101 | X_displacement | 0.0231056 | |
| Dataset | 0.00050962 | X_displacement | 0.06247925 | * |
| Injured | 0.00349069 | X_displacement | 0.11593435 | * |
| Time | 1.69E-20 | X_displacement | 0.13539793 | * |
| Blebbistatin:Injured | 0.00044837 | X_displacement | 0.21104785 | * |

| | | | | |
|---|---|---|---|---|
| Blebbistatin:Injured | 6.67E-15 | X_position | -0.4725283 | * |
| Blebbistatin | 8.00E-08 | X_position | -0.3554811 | * |
| Intercept | 7.06E-19 | X_position | -0.2937124 | * |
| Time | 0.68858853 | X_position | 0.00585397 | |
| Injured | 0.00203714 | X_position | 0.12318022 | * |
| Dataset | 7.58E-28 | X_position | 0.19899551 | * |
| Blebbistatin | 6.56E-14 | X_size | -0.5010008 | * |
| Time | 2.89E-13 | X_size | -0.1077415 | * |
| Injured | 0.8350012 | X_size | 0.00837548 | |
| Intercept | 0.78138785 | X_size | 0.00921345 | |
| Dataset | 4.83E-05 | X_size | 0.07405802 | * |
| Blebbistatin:Injured | 0.00029658 | X_size | 0.22052323 | * |
| Blebbistatin | 4.60E-07 | Y_displacement | -0.3296475 | * |
| Intercept | 0.0053783 | Y_displacement | -0.090589 | * |
| Dataset | 0.07015114 | Y_displacement | 0.03231419 | |
| Time | 1.14E-22 | Y_displacement | 0.14206158 | * |
| Blebbistatin:Injured | 0.00111544 | Y_displacement | 0.19463881 | * |
| Injured | 8.84E-16 | Y_displacement | 0.31800233 | * |
| Intercept | 1.72E-235 | Y_position | -0.703603 | * |
| Blebbistatin:Injured | 1.41E-13 | Y_position | -0.2748928 | * |
| Dataset | 1.49E-22 | Y_position | -0.1088347 | * |
| Time | 0.64528463 | Y_position | -0.0041219 | |
| Blebbistatin | 2.95E-57 | Y_position | 0.6557604 | * |
| Injured | 0 | Y_position | 1.73728264 | * |
| Blebbistatin:Injured | 6.78E-12 | Y_size | -0.4129694 | * |
| Intercept | 2.59E-30 | Y_size | -0.3771216 | * |
| Blebbistatin | 0.00796616 | Y_size | -0.1742958 | * |
| Time | 0.00052956 | Y_size | -0.0502928 | * |
| Dataset | 2.52E-11 | Y_size | 0.12001954 | * |
| Injured | 1.36E-45 | Y_size | 0.56797878 | * |
| Blebbistatin | 1.17E-07 | Z_displacement | -0.3504209 | * |
| Intercept | 0.00045356 | Z_displacement | -0.1154945 | * |
| Dataset | 0.00026137 | Z_displacement | 0.06594469 | * |
| Blebbistatin:Injured | 0.265347 | Z_displacement | 0.06725186 | |
| Time | 3.41E-22 | Z_displacement | 0.14206607 | * |
| Injured | 2.05E-10 | Z_displacement | 0.253986 | * |
| Injured | 1.26E-41 | Z_position | -0.5201777 | * |
| Dataset | 5.00E-80 | Z_position | -0.3334145 | * |
| Time | 0.0277476 | Z_position | -0.0306925 | * |
| Blebbistatin:Injured | 3.10E-15 | Z_position | 0.45664032 | * |
| Intercept | 1.20E-57 | Z_position | 0.51065591 | * |
| Blebbistatin | 8.90E-51 | Z_position | 0.95793721 | * |
| Blebbistatin | 0.00110497 | Z_size | -0.2200903 | * |
| Injured | 0.00087832 | Z_size | -0.135465 | * |
| Time | 0.45047573 | Z_size | 0.01123904 | |
| Dataset | 0.46374357 | Z_size | 0.01350125 | |

| | | | | |
|---|---|---|---|---|
| Intercept | 0.06110341 | Z_size | 0.06293508 | |
| Blebbistatin:Injured | 3.10E-09 | Z_size | 0.36597103 | * |
